# Sex-related structural alterations across common epilepsies: a worldwide ENIGMA study

**DOI:** 10.64898/2026.06.06.730611

**Authors:** Huantao Wen, Bin Wan, Taha Gholipour, Ke Xie, Judy Chen, Bianca Serio, Meike D. Hettwer, Marina K.M. Alvim, Donatello Arienzo, Rafael Batista João, Tobias Bauer, Andrea Bernasconi, Neda Bernasconi, Paolo Bonanni, Maria Eugenia Caligiuri, Fernando Cendes, Raphaël Christin, Luis Concha, Arielle Dascal, Edward Davoodi Boyd, Orrin Devinsky, Niels K.N. Focke, Francesco Fortunato, Marian Galovic, Antonio Gambardella, Renzo Guerrini, Sean N Hatton, Mirana Iliza, Sara Inati, Victoria Ives-Deliperi, Robert-Paul Juster, Jonathan K. Kleen, Mohamad Z Koubeissi, Angelo Labate, Sara Larivière, Meng Law, Matteo Lenge, Stefano Meletti, Patrick B. Moloney, Marlo Naish, Alexander Ngo, Terence J. O’Brien, Raluca Pana, Sofia Panzeri, Heath Pardoe, Erik H.U. Rauf, Antonella Riva, Raúl Rodríguez-Cruces, Jessica Royer, Theodor Rüber, Ella Sahlas, Lucas Scárdua-Silva, Kai Michael Schubert, Leigh N. Sepeta, Mariasavina Severino, Benjamin Sinclair, Hamid Soltanian-Zadeh, Travis Stoub, Pasquale Striano, Sophia I. Thomopoulos, Nina V. Valientes, Anna Elisabetta Vaudano, Lucy Vivash, Clarissa L. Yasuda, Paul M. Thompson, Sanjay M. Sisodiya, Carrie R. McDonald, Boris C. Bernhardt, Sofie L. Valk

**Affiliations:** Max Planck Institute for Human Cognitive and Brain Sciences, Leipzig, Germany; Institute of Neuroscience and Medicine (INM-7: Brain and Behaviour), Research Center Jülich, Jülich, Germany; Faculty of Life Science, Leipzig University, Leipzig, Germany; Department of Psychiatry, University Hospitals of Genève, Thonex, Switzerland; Synapsy Center for Neuroscience and Mental Health Research, University of Genève, Genève, Switzerland; Department of Neurosciences, 9300 Campus Point Drive, #7740, La Jolla, 92037, CA, USA; Centre for Excellence in Epilepsy at the Neuro (CEEN) and McConnell Brain Imaging Centre (BIC), Montreal Neurological Institute and Hospital, McGill University, Montreal, QC, Canada; Max Planck School of Cognition, Leipzig, Germany; Institute of Systems Neuroscience, Heinrich Heine University Düsseldorf, Düsseldorf, Germany; Penn Lifespan Informatics & Neuroimaging Center, Department of Psychiatry, Perelman School of Medicine, University of Pennsylvania, Philadelphia, Pennsylvania, USA; Department of Neurology, University of Campinas-UNICAMP, Rua Tessalia Vieira de Camargo 126, Campinas, 13083-887, SP, Brazil; Department of Radiation Medicine and Applied Sciences, University of California, San Diego, 9500 Gilman Drive, La Jolla, 92093, CA, USA; Brazilian Institute of Neuroscience and Neurotechnology, Rua Vital Brasil 251, Campinas, 13083888, SP, Brazil; Department of Neuroradiology, University Hospital Bonn, Venusberg-Campus 1, 53127, Bonn, Germany; Department of Epileptology, University Hospital Bonn, Venusberg-Campus 1, 53127, Bonn, Germany; German Center for Neurodegenerative Diseases (DZNE), Venusberg-Campus 1, Bonn, 53127, Germany; Neuroimaging of Epilepsy Laboratory, Montreal Neurological Institute, McGill University, Montreal, QC, Canada; Scientific Institute IRCCS E.Medea, Epilepsy and Clinical Neurophysiology Unit, Conegliano, Italy; Neuroscience Research Center, Department of Medical and Surgical Sciences, University “Magna Græcia" of Catanzaro, Catanzaro, Italy; Department of Neurology, 505 Parnassus Avenue, San Francisco, 94143, CA, USA; Instituto de Neurobiología, Universidad Nacional Autónoma de México, Querétaro, Mexico; Department of Neurology, Henry Ford Health, 2799 W Grand Blvd, Detriot, 48202, MI, USA; Epilepsy Center, Neurology Department, NYU Grossman School of Medicine, 223 E 34 Street, New York, 10016, NY, USA; Department of Neurology, Langone School of Medicine, New York University, New York, NY, USA; University Medical Center Göttingen, Clinic for Neurology, Göttingen, Germany; Institute of Neurology, Department of Medical and Surgical Sciences, Magna Graecia University of Catanzaro, Italy, Viale Europa, Catanzaro, 88100, Italy; Department of Neurology, Clinical Neuroscience Center, University Hospital and University of Zurich, Frauenklinikstrasse 26, Zurich, 8091, Switzerland; Department of Medical and Surgical Sciences, Institute of Neurology, Magna Graecia University, Viale Europa, Catanzaro, 88100, Calabria, Italy; Neuroscience Research Center, Magna Graecia University, Viale Europa, Catanzaro, 88100, Calabria, Italy; Neuroscience and Human Genetics Department, Meyer Children’s Hospital IRCCS, viale Pieraccini 24, Florence, 50139, Italy; University of Florence, Piazza San Marco, 450121 Florence, Italy; Department of Psychology, Bishop’s University, Sherbrooke, QC, Canada; Neurophysiology of Epilepsy Unit, National Institutes of Neurological Disorders and Stroke, National Institutes of Health, 10 Center Drive, Rm 7-5680, Bethesda, 20892, MD, USA; Department of Psychiatry, Neuroscience Institute, University of Cape Town, Anzio Road, Cape Town, 7800, Western Cape, South Africa; Department of Psychiatry and Addiction, University of Montreal, Montreal, H3T1J4, QC, Canada; Department of Neurology and Rehabilitation Medicine. George Washington University,Washington, USA; Neurology Clinic, Department of Neuroscience, Biomedicine and Advanced Diagnostic, University of Palermo, Palermo, Italy, Via del vespro 129, Palermo, 90127, PA, Italy; Department of Medical imaging and radiation sciences, Faculty of Medicine and Health Sciences, Université de Sherbrooke, Sherbrooke, QC, Canada; Department of Neuroscience, School of Translational Medicine, The Alfred Hospital, Monash University, 99 Commercial Road, Melbourne, 3004, Victoria, Australia; Biomedical Metabolic and Neural Sciences, OCB Hospital, Via giardini 1355, Modena, 41126, Italy; Neurophysiology Unit and epilepsy Centre, OCB Hospital, Modena, 41126, Italy; Research Department of Epilepsy, UCL Queen Square Institute of Neurology, Queen Square, London, WC1N 3BG, United Kingdom; Departments of Medicine and Neurology, The Royal Melbourne Hospital, The University of Melbourne, Royal Parade, Parkville, 3052, Victoria, Australia; Neuroradiology Unit, IRCCS Istituto Giannina Gaslini, via Gaslini 5, Genoa, 16147, Italy; New York University Grossman School of Medicine, 223 E 34th St, New York, 10016, NY, USA; Florey Institute of Neuroscience and Mental Health, 245 Burgundy St, Heidelberg, 3084, Victoria, Australia; Department of Neurosciences, Rehabilitation, Ophthalmology, Genetics, Maternal and Child Health, University of Genoa, Via Gerolamo Gaslini 5, Genoa, 16147, Genoa, Italy; Multimodal Imaging and Connectome Analysis Laboratory, McConnell Brain Imaging Centre, Montreal Neurological Institute and Hospital, McGill University, Montreal, QC, Canada; German Center for Neurodegenerative Diseases (DZNE), Campus Venusberg 99, Bonn, 53127, Germany; Center for Medical Data Usability and Translation, University of Bonn, Konviktstr. 4, 53115 Bonn, Germany; Neuroimaging Laboratory, University of Campinas (UNICAMP), Brazil, Campinas, Sâo Paulo, Brazil; Department of Neuropsychology, Children’s National Hospital, 111 Michigan Avenue NW Washington, 20010, USA; School of Translational Medicine, Monash University, 99 Commercial Road, Melbourne, 3004, VIC, Australia; Departments of Research Administration and Radiology, Henry Ford Health, 1 Ford Place, Detriot, 48202, MI, USA; School of Electrical and Computer Eng., North Kargar Ave., Tehran, 1439957131, Tehran, Iran; Neurological Sciences Rush University Medical Center, Chicago, IL, USA; IRCCS G. Gaslini, full member of Epicare, Genova, Italy, Genova, Italy; Imaging Genetics Center, Mark and Mary Stevens Neuroimaging and Informatics Institute, Keck School of Medicine, University of Southern California, 4676 Admiralty Way, Marina del Rey, 90033, CA, USA; Department of Biomedical, Metabolic and Neuronal sciences, University of Modena and Reggio-Emilia, Modena, Italy; Department of Neurology, Alfred Health, Melbourne, VIC, Australia; Neurology Department, University of Campinas (UNICAMP), Campinas, São Paulo, Brazil; Chalfont Centre for Epilepsy, Chesham Lane, Chalfont St Peter, SL9 0RJ, Bucks, UK; Department of Psychiatry, University of California, San Diego, 9500 Gilman Drive, La Jolla, 92093, CA, USA

## Abstract

Epilepsy is characterized by widespread structural brain alterations extending beyond the epileptic zone, involving both cortical and subcortical regions. Importantly, the clinical manifestation of epilepsy, including seizure types, psychiatric comorbidities, and treatment responses, has been shown to differ between sexes. However, sex differences in structural alterations in epilepsy have been seldomly reported in neuroimaging studies, partly due to limited sample sizes and single-center designs. Here, we systematically investigated sex differences in common epilepsies and their related clinical variables using structural neuroimaging biomarkers in an international multi-center cohort of 1,253 epilepsy patients and 1,077 healthy controls. We studied cortical thickness and subcortical volume in two types of epilepsy: temporal lobe epilepsy (TLE) and genetic generalized epilepsy (GGE). Both male and female patients with TLE showed widespread cortical and subcortical thinning compared with controls. In GGE, when compared separately to controls, male patients showed only subtle structural alterations, whereas female patients exhibited more widespread structural alterations. Sex-stratified analyses revealed some variation in the extent and distribution of cortical thickness and subcortical volume alterations between male and female patients in both epilepsy cohorts. Yet, we did not find significant sex-by-diagnosis interaction effects in TLE and GGE. Similarly, no significant interaction effects were observed between sex and age of onset or disease duration in either patient group. Overall, although we observed some differences in regional cortical thickness and subcortical volume between male and female patients with epilepsy, we did not find significant sex-by-diagnosis interactions. Our findings indicate that sex differences in behavioral and clinical outcomes of epilepsy may involve biological or functional processes that require further investigation.

## Introduction

Structural alterations have been consistently reported across both focal and generalized epilepsy in neuroimaging studies. These alterations have extended beyond the epileptic zone, involving widespread cortical and subcortical regions^1–5^. In temporal lobe epilepsy (TLE), pronounced volume loss has been observed in the hippocampus and thalamus, along with cortical thinning in entorhinal, parietal, frontal, and other neocortical regions^6^. Genetic generalized epilepsy (GGE) is characterized by thalamic atrophy and cortical thinning involving frontal, central, and temporal areas^7^. Importantly, grey matter alterations in both TLE and GGE are not static but change along the course of the disease. Evidence from longitudinal and cross-sectional studies indicates that longer disease duration is associated with more pronounced grey matter atrophy in TLE^8,9^. In GGE, longer epilepsy duration is also associated with more pronounced grey matter loss, especially within the areas involving thalamocortical networks^10,11^. These widespread and progressive grey matter changes might also be associated with poor seizure control, as well as cognitive and behavioural abnormalities across individuals with epilepsy^8,12,13^.

Crucially, the clinical presentation of epilepsy, including seizure types, psychiatric and cognitive comorbidities, as well as treatment outcomes, has been shown to differ between sexes^14–16^. Epidemiological studies indicate that GGE is more prevalent among females, whereas structural focal epilepsies, such as TLE, occur more frequently in males^17–19^. Psychiatric comorbidities also show marked sex differences, with female patients more likely to experience depression and anxiety compared to males^20^, and risk factors contributing to the psychiatric comorbidities of epilepsy may also differ across male and female patients^21^. In terms of cognitive impairments in epilepsy, male and female patients have also been reported to perform differently in visual and verbal memory^22,23^ and exhibit different emotional responses^21^. These differences exist in both pre- and post-surgical treatments, and might influence the surgical outcomes among epilepsy patients^24^.

Sex differences in the clinical presentation of epilepsy may be attributed to both gender-based social-environmental factors and sex-based biological mechanisms. Epidemiological studies indicate that males have a higher lifetime risk of epilepsy, which might be attributed to their occupations and exposure to risk factors, such as head trauma and alcohol use^25^. From a biological perspective, sex differences in epilepsy risk factors have been reported at multiple levels, including genetic, connectomic, and neuroendocrine processes. Sex differences in gene expression may differentially modulate neuronal excitability by regulating neurotransmitter synthesis and synaptic transmission, thereby contributing to sex differences in seizure susceptibility^15,26^. Sex-specific differences in brain connectivity may also contribute to differential epilepsy vulnerability^27^. For instance, males exhibit higher intrahemispheric connectivity, whereas females exhibit stronger interhemispheric connectivity, which may partly explain the higher prevalence of generalized epilepsies among women^28^. Moreover, neuroendocrine influences might also contribute to the sex difference of epilepsy, as sex hormones critically influence neuronal excitability, gene expression, and brain plasticity^29^. In addition, hormonal fluctuations across the lifespan may shape sex-specific trajectories of brain structural and functional organization^30–32^ and differentially modulate neuronal excitability in males and females, particularly during critical periods such as puberty, pregnancy, and menopause^33,34^. Together, these neuroendocrine influences may contribute to sex differences in epilepsy susceptibility and clinical expression^35^.

Despite evidence suggesting the relevance of considering sex differences in epilepsy, neuroimaging studies investigating the role of sex in structural alterations in epilepsy are limited. In TLE, male patients exhibit greater brain atrophy than female patients in ipsilateral hippocampus^36^. Similarly, female patients with TLE associated with mesial temporal sclerosis (TLE-MTS) have been reported to show more pronounced structural alterations in temporal regions, whereas male patients with MTS display greater abnormalities in frontal areas^37,38^. However, these findings are mostly derived from small sample sizes and single-site designs, which may limit the statistical power to detect the sex-specific structural alteration patterns in epilepsy and require replication in larger, multisite cohorts.

The present study systematically investigates sex differences in structural brain alterations in epilepsy using data from the Enhancing Neuro Imaging Genetics through Meta Analysis Epilepsy Consortium (ENIGMA)-Epilepsy consortium, a large-scale, international multi-center cohort. To increase the statistical power, we integrated the ENIGMA dataset with two additional independent datasets. We compared cortical thickness and subcortical volume derived from structural MRI between male and female patients with TLE and GGE. To estimate whether males and females exhibited distinct spatial patterns of cortical and subcortical alterations in TLE and GGE, we first conducted sex-stratified case-control comparisons. We then tested for significant sex-by-diagnosis interactions. In addition, we investigated whether disease duration or age of onset would differentially influence cortical thickness and subcortical volume in male and female patients. Similar to case-control analyses, we further evaluated sex-by-disease duration and sex-by-age of onset interactions to determine their associations with cortical thickness and subcortical volume.

## Results

### Data samples

Our main analyses included patients with TLE (*n* = 1,038) and GGE (*n* = 215), together with their corresponding site-matched healthy controls (*n*_TLE-HC_ = 705 and *n*_GGE-HC_ = 372, respectively). No significant sex differences were observed in age within each diagnostic group, or in the side of epileptic focus among TLE patients. All participants were aged between 18 and 65 years. Details of all participants and their demographic characteristics are summarized in **Table 1**, and sex differences of each participant group are provided in **Table 2**. Details of ENIGMA and two independent datasets are provided in **Supplementary Table 1**. Subject inclusion details are listed in **Materials and Methods**.

**Table 1.**
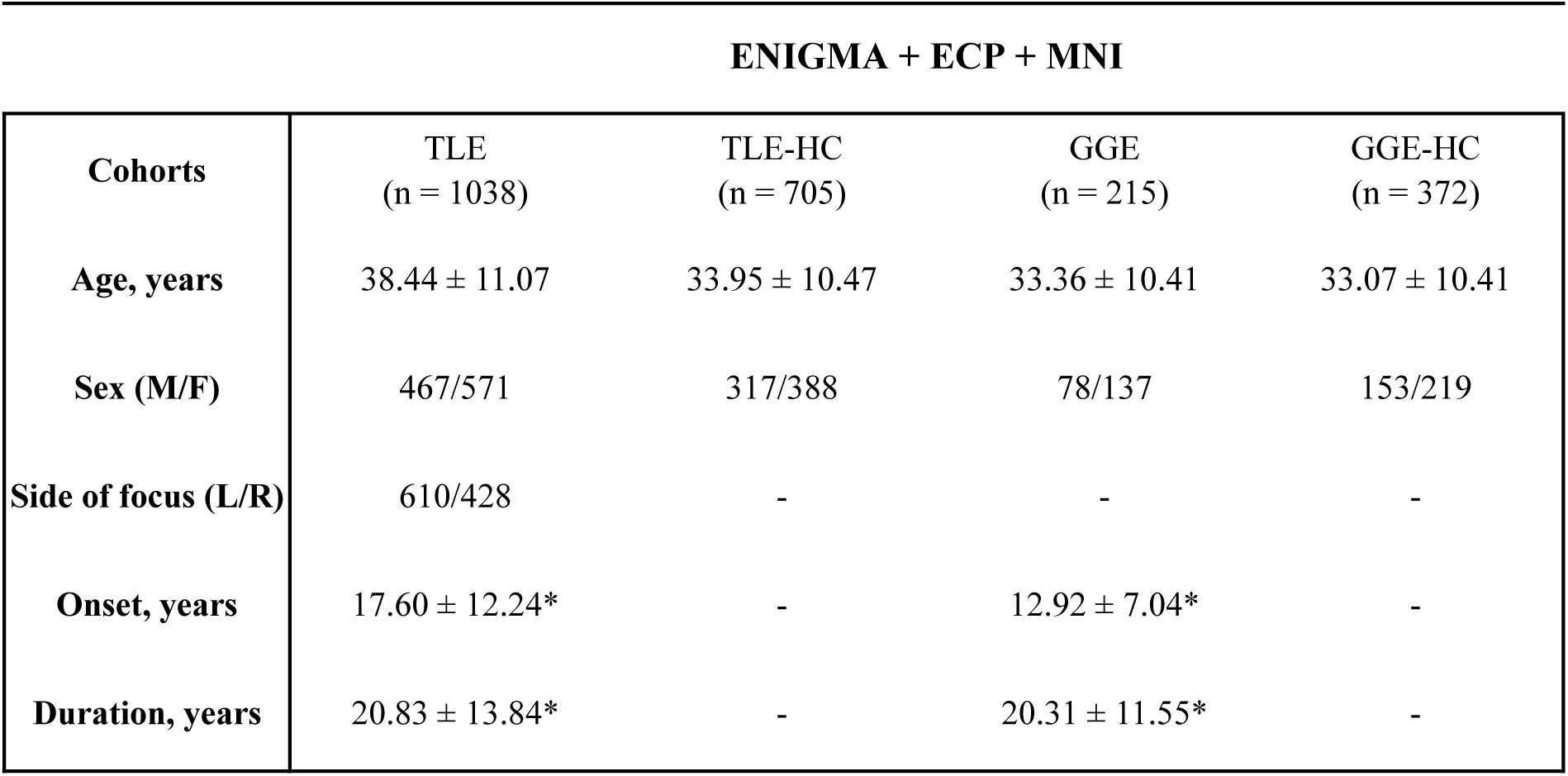
Demographics and clinical information. Demographics are shown for patient subcohorts and their site-matched controls in the merged dataset. Age, age of onset, and disease duration of epilepsy are presented as mean ± SD years. Side of seizure focus (patients with TLE only). TLE: temporal lobe epilepsy; GGE: genetic generalized epilepsy; HC: healthy control; Onset: age of onset; Duration: disease duration; L = left; R = right. *Information is available in 1,015 of 1,038 patients with TLE and 171 of 215 patients with GGE.

**Table 2.** Demographic information comparison across sexes. Statistical comparisons between males and females were shown for patient subcohorts and their site-matched controls in the merged dataset. Age, age of onset, and disease duration of epilepsy are presented as mean ± SD years. Age, age at onset, and disease duration were compared using two-sample t-tests, while side of seizure focus was compared using Chi-square tests, with statistical significance set at *p* < 0.05. TLE: temporal lobe epilepsy; GGE: genetic generalized epilepsy; HC: healthy control; Onset: age of onset; Duration: disease duration; L = left; R = right.

| ENIGMA + ECP + MNI |  |  |  |
| --- | --- | --- | --- |
|  | Male | Female | *P* |
| **Age TLE** | 37.76 ± 11.30 | 38.99 ± 10.86 | 0.076 |
| **Age TLE-HC** | 33.59 ± 10.51 | 34.24 ± 10.44 | 0.409 |
| **Age GGE** | 33.68 ± 10.98 | 33.17 ± 10.11 | 0.739 |
| **Age GGE-HC** | 32.32 ± 10.27 | 33.59 ± 10.51 | 0.246 |
| **Onset TLE** | 17.33 ± 12.16 | 17.82 ± 12.31 | 0.529 |
| **Onset GGE** | 13.23 ± 5.47 | 12.77 ± 7.72 | 0.653 |
| **Duration TLE** | 20.65 ± 13.87 | 20.98 ± 13.83 | 0.712 |
| **Duration GGE** | 20.44 ± 12.71 | 20.25 ± 10.98 | 0.924 |
| **TLE Side of focus (L/R)** | 274/193 | 336/235 | 1.0 |

### Sex-stratified case-control differences in the common epilepsies (Figure 1)

We first examined differences between individuals with epilepsy and healthy controls in males and females. Both males and females showed widespread cortical and subcortical atrophy in TLE compared to healthy controls. We found significant negative effects in the bilateral superior parietal, paracentral, postcentral, precentral lobes, bilateral thalamus, and hippocampus in both male (*d* = -0.243 to -0.165, *P*_FDR_ < 0.001) and female (*d* = -0.283 to -0.183, *P*_FDR_ < 0.001) patients. In GGE, male patients exhibited primarily atrophy in right paracentral, bilateral precentral, and transverse temporal areas (*d* = -0.300 to -0.197, *P*_FDR_ < 0.05), and increased cortical thickness in left rostral middle frontal and right entorhinal areas (*d* = 0.211 to 0.242, *P*_FDR_ < 0.05). Female patients exhibited more extensive cortical and subcortical atrophy, mainly within the left hemisphere and predominantly in temporal regions, inferior parietal and lateral occipital lobes (*d* = -0.216 to -0.147, *P*_FDR_ < 0.05), as well as bilateral striatum, hippocampus, and amygdala (*d* = -0.255 to -0.110, *P*_FDR_ < 0.001), with no areas of increased volume or cortical thickness. Altogether, our sex-stratified analyses revealed that both males and females showed extensive structural changes in TLE, with slightly more widespread effects in females relative to sex-matched controls. In GGE, sex-stratified comparisons with controls revealed subtle cortical thinning in males but more pronounced and widespread cortical thinning in females. Detailed statistical estimates are shown in **Supplementary Tables 3 and 4**.

**Figure 1.**
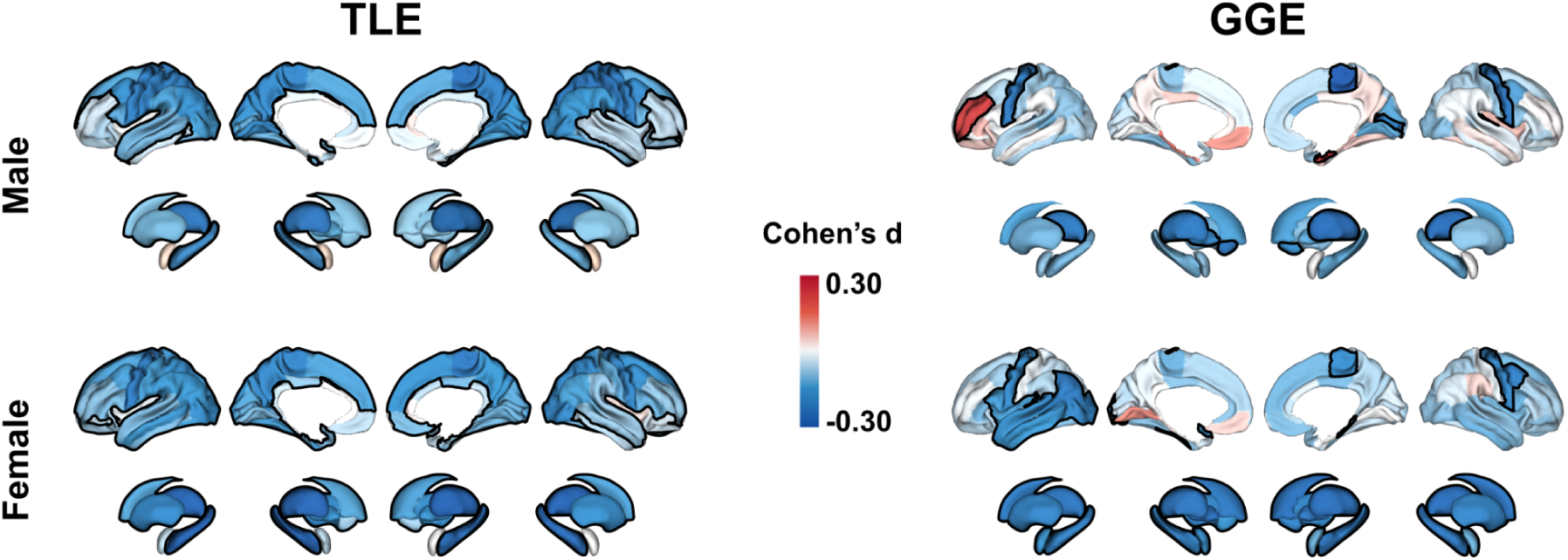
Case-control differences in common epilepsies within males and females. Cohen’s d values for case-control differences in cortical thickness and subcortical volume in TLE and GGE are displayed for males (top) and females (bottom). We summarized the effect sizes using Cohen’s d, where a positive d value (red) indicates higher values in epilepsy patients than controls, and a negative d value (blue) reflects higher values in controls relative to TLE/GGE patients. Significant regions are outlined by dark lines with a threshold of *P*_FDR_ < 0.05. TLE: temporal lobe epilepsy; GGE: genetic generalized epilepsy.

### Sex differences in epilepsy and corresponding healthy controls (Figure 2)

We next examined sex differences within epilepsy and their corresponding control groups. In TLE, only subtle differences of sex effects were observed in the cingulate area, whereas males displayed greater subcortical volumes than females, most prominently in the bilateral putamen, pallidum, and thalamus (*d* = 0.162 to 0.29, *P*_FDR_ < 0.001). In GGE, males showed greater cortical thickness than females, particularly in the bilateral temporal cortex, insula, and entorhinal cortex (*d* = 0.218 to 0.334, *P*_FDR_ < 0.05), as well as greater subcortical volumes in the bilateral hippocampus and amygdala (*d* = 0.25 to 0.38, *P*_FDR_ < 0.001). The corresponding control groups of both TLE and GGE showed similar patterns with only minor cortical differences and generally larger subcortical volumes in males (**Supplementary Tables 5 and 6**). Finally, we examined diagnosis-by-sex interaction effects. We found no significant interactions across cortical regions in either TLE (*d* = -0.052 to 0.041) or GGE (*d* = -0.074 to 0.115), as well as subcortical regions (**Supplementary Tables 11 and 12**). Overall, our findings are consistent with the case-control findings within males and females, showing that females exhibit more widespread cortical atrophy than males in GGE. Yet, based on the diagnosis-by-sex interaction analyses, neither cortical nor subcortical atrophy appears to be significantly influenced by sex in either TLE or GGE. Nevertheless, the broader cortical and limbic alterations observed in females with GGE in sex-stratified analyses suggest a possible sex-related structural vulnerability that may lie below the sensitivity of the present interaction models.

**Figure 2.**
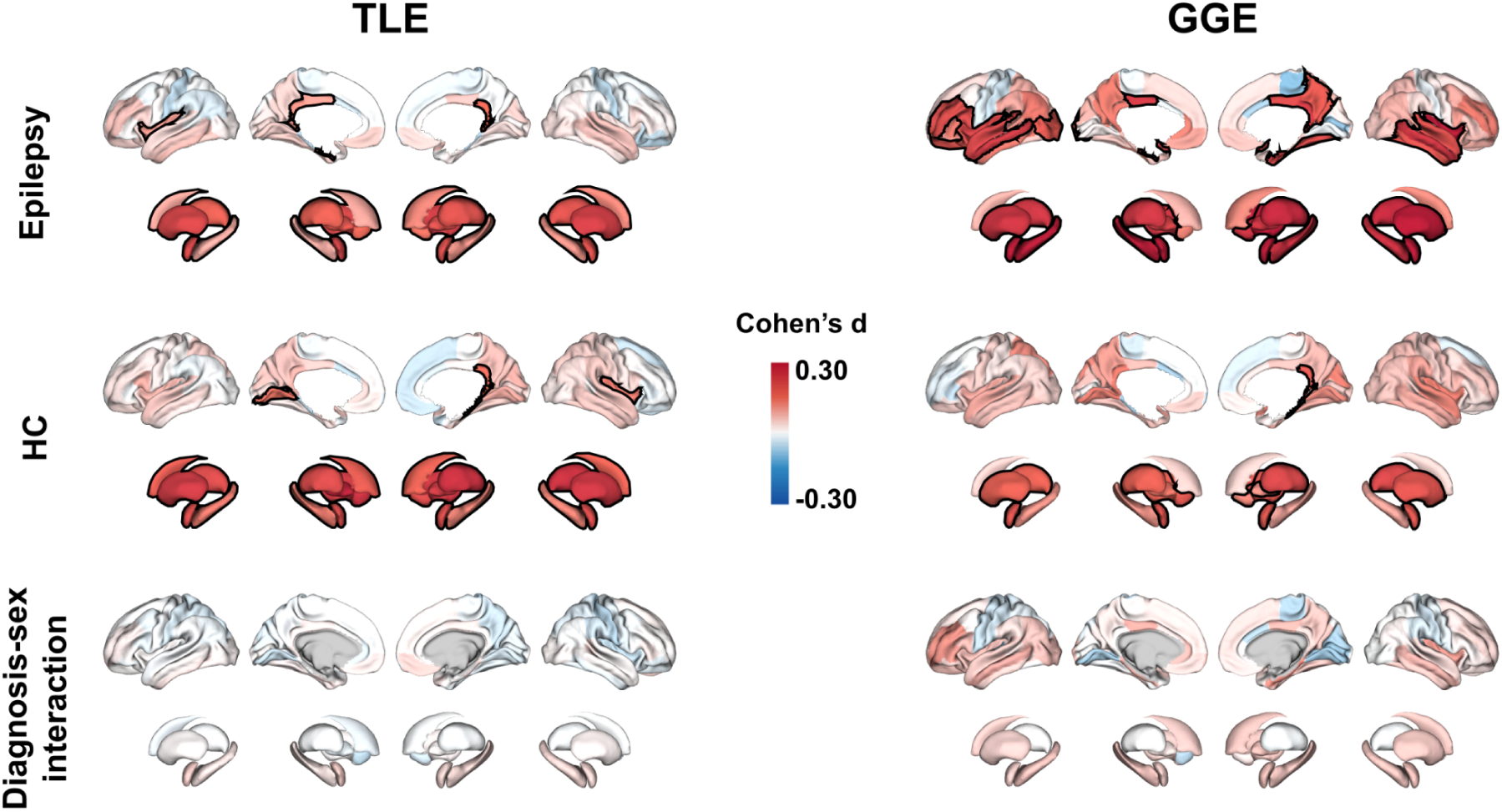
Sex differences in TLE and GGE and corresponding healthy controls. Upper and middle panels show the effect sizes of sex on cortical thickness and subcortical volume in the epilepsy and site-matched control groups. The lower panel depicts diagnosis-by-sex interaction effects. We summarized the effect sizes using Cohen’s d, where a positive d value (red) indicates higher values in male participants than female participants, and a negative d value (blue) reflects higher values in female participants relative to male participants. Significant regions were outlined by dark lines with a threshold of *P*_FDR_ < 0.05. TLE: temporal lobe epilepsy; GGE: genetic generalized epilepsy; HC: healthy control.

### Sex differences in relation to disease duration and age of onset (Figure 3, Figure 4)

We further examine whether structural alterations are differentially associated with disease duration and age of onset across sexes. We first investigated the effects of disease duration on cortical thickness and subcortical volumes in males and females with TLE and GGE separately. In TLE, both males (*d* = -0.310 to 0.004) and females (*d* = -0.248 to 0.034) showed widespread negative associations between disease duration and cortical thickness. Negative associations were also captured in subcortical areas in both males (*d* = -0.349 to -0.101) and females (*d* = -0.344 to -0.109). These findings indicate that longer disease duration is related to greater structural atrophy in both male and female TLE patients. In GGE, disease duration showed more extensive negative effects in cortical regions among female patients. Disease duration was negatively associated with cortical thickness (*d* = -0.577 to -0.418, *P*_FDR_ < 0.05) in male patients in the bilateral isthmus cingulate, left middle temporal, and right superior temporal lobes, whereas female patients displayed wider negative disease duration effects (*d* = -0.442 to -0.231, *P*_FDR_ < 0.05), predominantly in bilateral superior frontal and caudal middle frontal, cingulate, temporal, and insula areas (**Supplementary Table 7**). Both male and female GGE patients displayed widespread negative correlations with disease duration in subcortical volume (**Supplementary Table 8**). To further examine whether disease duration differentially affects cortical and subcortical atrophy across sexes, we tested the disease duration-by-sex interaction effects. We found no significant effects in TLE or GGE in cortical or subcortical regions, suggesting that disease duration influences cortical and subcortical atrophy similarly in males and females (**Supplementary Tables 11 and 12**).

**Figure 3.**
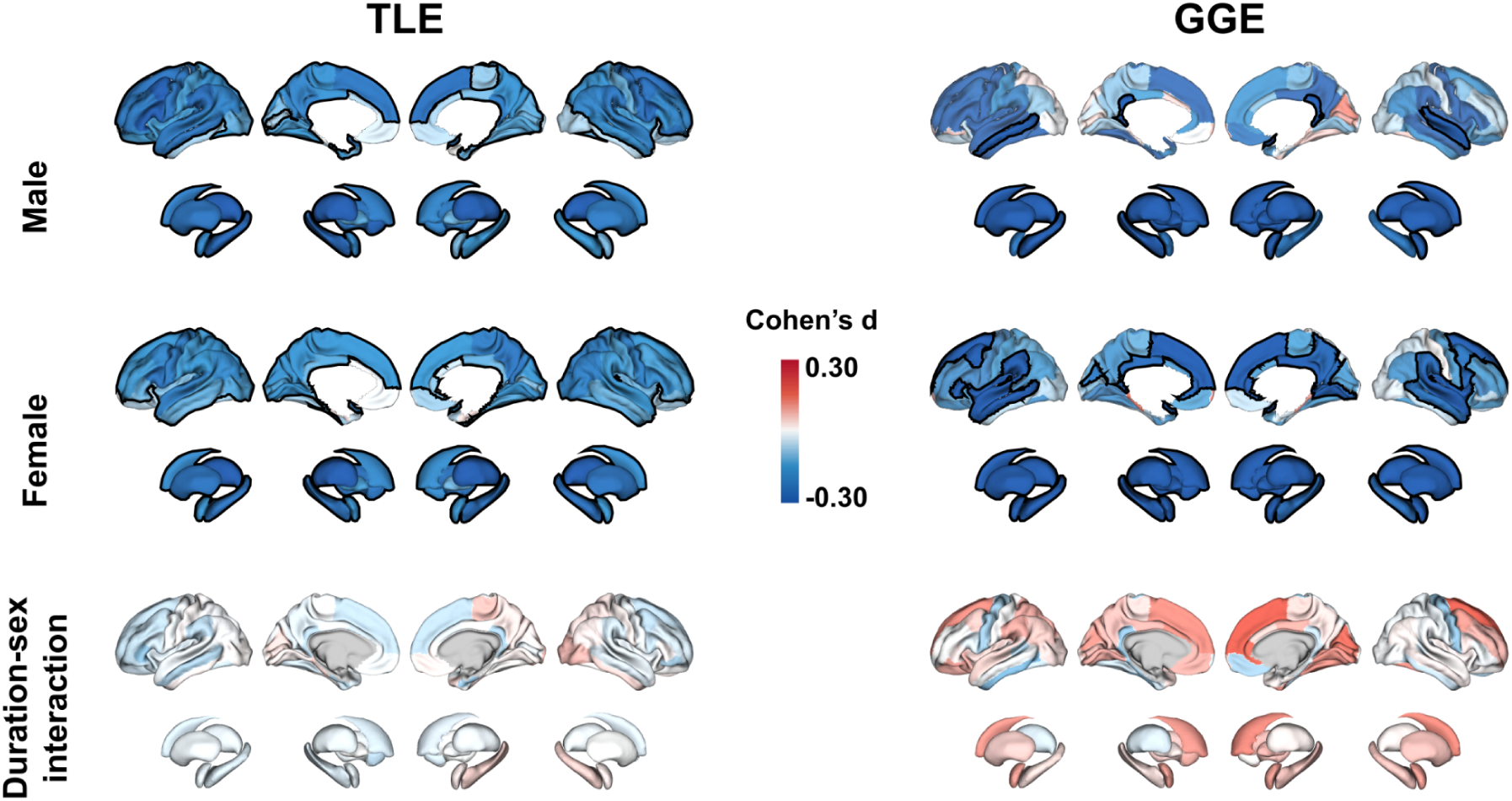
Sex differences in the effect of disease duration. Effect sizes are shown for the main effect of disease duration (upper and middle panels) and for the disease duration-by-sex interaction (lower panel). Positive values (red) denote regions where longer disease duration is associated with greater cortical thickness or where male patients show stronger associations with disease duration. Statistically significant regions are outlined by dark borders (*P*_FDR_ < 0.05). TLE: temporal lobe epilepsy; GGE: genetic generalized epilepsy; Duration: disease duration.

**Figure 4.**
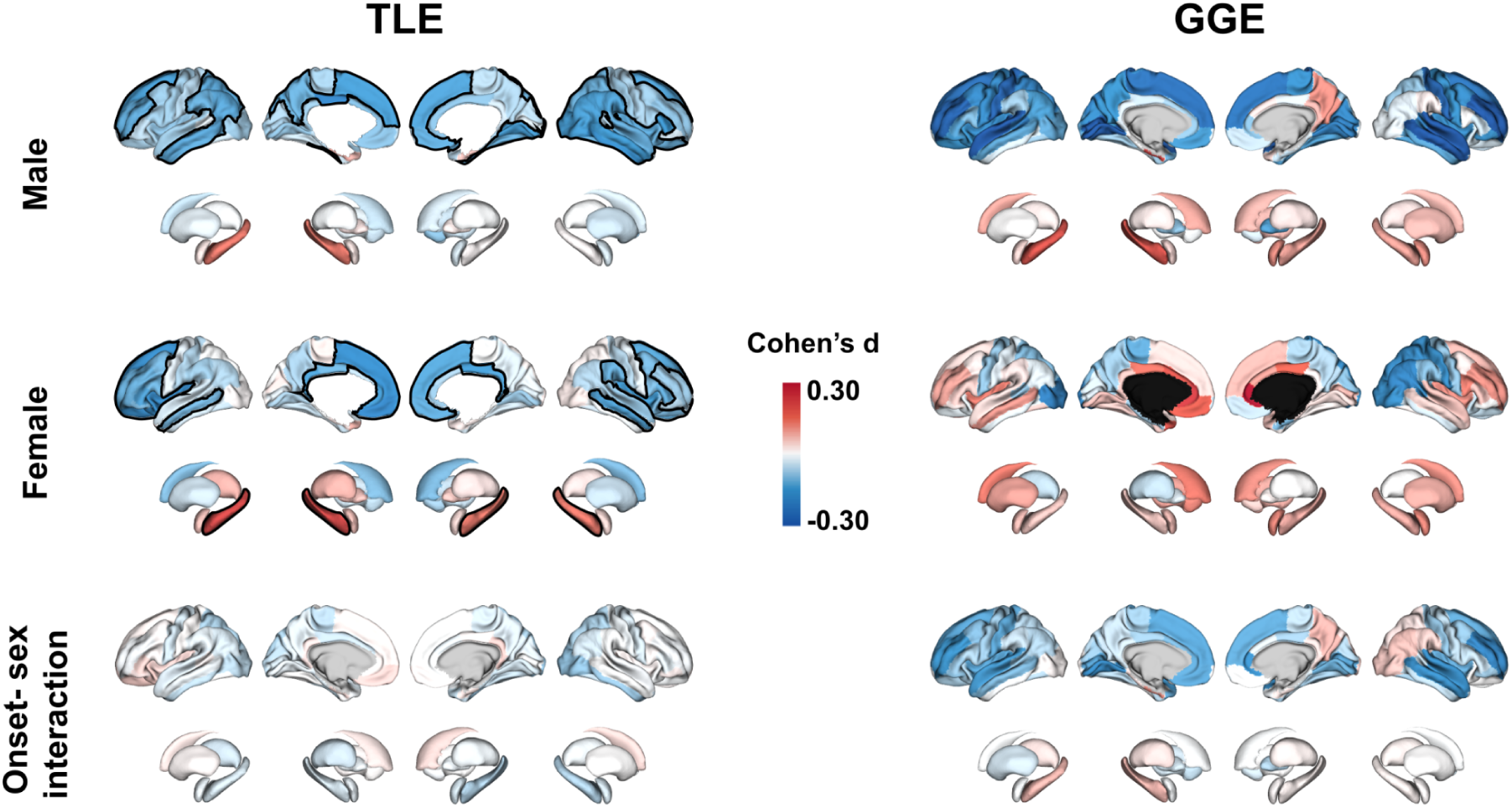
Sex differences in the effect of age of onset. Effect sizes are shown for the main effect of age of onset (upper and middle panels) and for the age of onset-by-sex interaction (lower panel). Positive values (red) indicate regions where a later onset corresponds to greater cortical thickness or where males exhibit stronger age of onset-related effects. Statistically significant regions are outlined by dark borders (*P*_FDR_ < 0.05). TLE: temporal lobe epilepsy; GGE: genetic generalized epilepsy; Onset: age of onset.

Next, we examined whether structural alterations were differentially related to age of onset across sexes in TLE and GGE. In TLE, age of onset showed negative associations with cortical thickness in male patients, mainly in the left posterior cingulate, bilateral temporal, and parietal cortex in male patients (*d* = -0.201 to -0.127, *P*_FDR_ < 0.05). In female patients, negative associations (*d* = -0.188 to -0.108, *P*_FDR_ < 0.05) were mainly found in the bilateral frontal, insula, and isthmus cingulate cortex, while a positive association emerged in the left (*d* = 0.201, *P*_FDR_ < 0.001) and right hippocampus (*d* = 0.129, *P*_FDR_ < 0.05). In GGE, no significant associations with age of onset were detected in either cortical or subcortical regions for males or females (**Supplementary Tables 9 and 10**). Further analyses of age of onset-by-sex interaction effects revealed no significant sex difference in TLE and GGE in cortical or subcortical regions across males or females (**Supplementary Tables 11 and 12**), which indicate the association between age of onset and cortical thickness or subcortical volume did not differ significantly between male and female patients. Together, although the age of onset associated with male and female patients differently in cortical thickness and subcortical volume in sex-stratified analyses in TLE, the interaction analyses suggest that age of onset does not differentially relate to macroscale brain structure in male and female patients.

### Robustness analyses of sex effects

Previous studies also suggested left and right TLE displayed different patterns of structural deficits^6,39^, as well as different cognitive outcomes across TLE patients^40,41^. To account for the effect of lateralization of TLE, we computed diagnosis-by-sex, disease duration-by-sex, and age of onset-by-sex interaction effects separately in left and right TLE. In cortical thickness, effect sizes were small and non-significant for diagnosis-by-sex (*d*_L-TLE_ = -0.048 to 0.052; *d*_R-TLE_ = -0.072 to 0.049), disease duration-by-sex (*d*_L-TLE_ = -0.107 to 0.097; *d*_R-TLE_ = -0.104 to 0.125), and age of onset-by-sex effects (*d*_L-TLE_ = -0.104 to 0.068; *d*_R-TLE_ = -0.094 to 0.085). Similarly, no significant interaction effects were observed in subcortical volumes (**Supplementary Fig. 1, Supplementary Tables 13 and 14**). Additionally, to address the potential influence of lesion effects in TLE patients, we tested the main sex interaction effects above in TLE with and without MTS, respectively. These effects remained non-significant in both TLE-MTS and non-lesional TLE across cortical and subcortical regions (**Supplementary Fig. 2, Supplementary Tables 15 and 16**). These results revealed that lateralization in TLE, and TLE with or without MTS did not relate to sex-related effects in structural alterations in TLE.

Furthermore, we examined the age effects separately in male and female patients with TLE and GGE to see if age affects male and female patients differently in cortical thickness and subcortical volume. In TLE, age was negatively associated with cortical thickness and subcortical volumes in both male and female patients. In males, negative effects (*d* = -0.519 to -0.371, *P*_FDR_ < 0.001) were primarily observed in bilateral frontal, supramarginal, left middle temporal, left posterior cingulate areas, bilateral thalamus, and accumbens. In females, negative effects (*d* = -0.500 to -0.316, *P*_FDR_ < 0.001) were mainly observed in bilateral frontal and parietal areas, as well as putamen and thalamus. In GGE, male patients showed negative age effects (*d* = -0.866 to -0.503, *P*_FDR_ < 0.05) primarily involving left temporal and bilateral isthmus cingulate, insula, thalamus, and putamen areas. In female patients, negative effects (*d* = -0.609 to -0.314, *P*_FDR_ < 0.05) were mainly observed across bilateral superior frontal, superior temporal, right banks of the superior temporal sulcus and left precentral areas, as well as bilateral thalamus and caudate. We next tested age-by-sex interaction effects in TLE and GGE to assess whether age-related associations differed between males and females. No significant age-by-sex interaction effects were detected in TLE and GGE across cortical or subcortical regions (**Supplementary Fig. 3, Supplementary Tables 17 and 18**). These results suggested that age-related effects with cortical and subcortical measures were largely similar between male and female patients in TLE and GGE.

Lastly, to test the finding consistency across three datasets, we examined the diagnosis-by-sex, disease duration-by-sex, and age of onset-by-sex effects in three datasets by fitting the same linear models as in the merged dataset. Consistent with the findings in the merged dataset, all these effects are nonsignificant in three datasets, indicating that sex does not modulate diagnosis, disease duration, and age of onset effects on cortical thickness or subcortical volume in these samples respectively (**Supplementary Fig. 4, Supplementary Tables 19 , 20 , 21, and 22**).

## Discussion

We conducted a comprehensive investigation of sex-related differences in structural brain alterations among individuals with epilepsy. Taken together, this work presents the first large-scale, multisite neuroimaging study to systematically assess sex-related structural differences in common epilepsies. We found no sex-by-diagnosis interactions, nor sex interaction effects with disease duration or age of onset, on cortical thickness or subcortical volume in either TLE or GGE. Nevertheless, sex-stratified case-control analyses revealed more widespread structural alteration patterns in females than in male patients, particularly in GGE. The absence of significant interaction effects should be interpreted specifically within the sensitivity limits of macroscale cortical thickness and subcortical volumetry. These findings do not exclude sex-dependent differences in brain microstructure, network organisation, neurochemistry, or endocrine modulation, which may account for clinically observed sex differences in epilepsy.

Performing case-control analyses on males or females separately in both TLE and GGE, we observed that both male and female patients exhibited widespread cortical and subcortical volume reductions in TLE, and structural alterations were more limited in both sexes in GGE. These findings are consistent with previous ENIGMA-Epilepsy case-control results, which reported widespread cortical and subcortical atrophy in TLE but fewer alterations in GGE^6,7^. However, in sex-stratified comparisons with controls, female patients showed qualitatively more widespread cortical alterations than males in both TLE and GGE, and this difference was more apparent in GGE. Although these differences did not reach statistical significance in diagnosis-by-sex interaction effects, these results might still indicate differentiable structural vulnerability in each sex in GGE.

Notably, sex differences in cortical atrophy in GGE have been rarely described in prior neuroimaging studies. A previous investigation by Ogren et al^42^ reported male GGE patients exhibiting posterior cingulate thickening in the right hemisphere, while females showed only left lateral orbitofrontal thinning. However, their sample size was small (*n* = 53) and they combined patients with both generalized and initially focal temporal lobe onsets, which may have contributed to inconsistent findings across studies. Previous voxel-based morphometry (VBM) studies examining sex differences in TLE have reported that grey matter volume reductions in female patients are predominantly confined to temporal regions, whereas males show more pronounced involvement of frontal areas^37,38^. In contrast, we observed widespread cortical atrophy across bilateral frontal and temporal lobes in female patients with TLE, which did not differ significantly from the extent of atrophy observed in males. This discrepancy might be due to differences in imaging metrics as well as the limited sample sizes (*n* = 120 and *n* = 28) in earlier studies.

The lack of significant diagnosis-by-sex interaction effects on cortical thickness and subcortical volume can be understood in the context of how anatomical sex differences across healthy adult participants. In previous neuroimaging studies of sex differences in brain structure across healthy participants, females generally exhibit greater cortical thickness, particularly in frontal and parietal regions^43,44^, whereas males show higher subcortical volumes, with the largest differences typically observed in the bilateral thalamus, putamen, pallidum^44^. Such sex-related structural variations are thought to be shaped by complex genetic and environmental interactions, which may contribute to differential susceptibility to neuropsychiatric or neurological disorders^44–47^. However, effect sizes of sex differences in grey matter volume across brain regions are generally small to moderate, as reported in large population-based cohort studies^48–50^. Grey matter reductions observed with MRI in epilepsy may reflect underlying pathological changes, including neuronal loss and synaptic reorganization^9,51^, which may be related to mechanisms such as microglial activation, tau deposition, neuroinflammation, and others^52–54^. Currently, how sex is involved in these mechanisms remains unknown. Although animal studies have reported that male mice show greater hippocampal cell loss and increased hippocampal gliosis following development of TLE^55,56^, there is so far no direct pathological evidence in humans demonstrating that males or females with TLE exhibit a higher or lower rate of neuronal loss. On the other hand, the mechanisms of sex differences in epilepsy are widely reported in hormonal modulation and neurotransmitter regulation^15,56^, which may lead to differential susceptibility to epilepsy, such as catamenial epilepsy, the most widely reported sex difference in epilepsy. These observations may suggest that sex differences in epilepsy are more strongly related to neuroendocrine influences, neurochemical modulation, and related neurobiological mechanisms, which may preferentially influence neuronal excitability and network dynamics rather than causing macroscale structural alterations detectable by MRI.

Both male and female patients exhibited widespread negative associations between disease duration and cortical thickness as well as subcortical volume in TLE. These findings align with previous studies reporting that longer disease duration is consistently associated with more pronounced cortical and subcortical atrophy in TLE^6,8,9,12,57^. However, we did not observe any significant sex-related differences in disease duration. Santana et al^37^ reported region-specific cortical thinning patterns, with female patients showing more pronounced temporal lobe changes and males exhibiting greater frontal alterations. Notably, male TLE patients in that study had a shorter disease duration than female patients, implying that apparent sex-related differences in cortical alterations may partly reflect differences in cumulative disease exposure. In contrast, our findings suggest that when disease duration is comparable, the neuroanatomical impact of epilepsy progression is similar between sexes. Our findings align with previous studies that disease duration effects on the brain structure of GGE patients are more limited compared to TLE^6,7,10,58^. This might be due to disease duration has less influence on brain atrophy in GGE compared to focal epilepsy, or differences in location and frequency of epileptiform activity also lead to different neuronal damage than TLE^7^. So far, less evidence on the effects of disease duration comparison between males and females in TLE or GGE. Carlson et al^59^ reported that higher rates of autonomic, psychic, and visual symptoms were observed in female focal epilepsy patients compared to males, and a higher frequency of atonic seizures in male GGE patients, with a sample that females have higher disease duration. However, direct neuroimaging evidence comparing sex differences in the association between disease duration and brain structure remains limited. Our results provide new evidence showing that the association between disease duration and brain structure does not differ by sex in either TLE or GGE, for both cortical thickness and subcortical volume.

Age of onset effects did not differ between males and females in either TLE or GGE. Nonetheless, sex-stratified analyses showed that more widespread cortical thinning was associated with later onset in female patients with TLE. Similar to disease duration, few structural neuroimaging studies have systematically investigated sex differences in age of onset effects in TLE or GGE. A previous study^37^ reported that female TLE patients showed more pronounced temporal cortical thinning, whereas males exhibited stronger frontal atrophy in a cohort where male TLE patients had a higher age of onset than females, suggesting that age of onset might interact differently with sex. Doherty et al^60^ reported that the epileptiform activity in adult patients with focal epilepsy is different between male and female patients, and might be related to age at epilepsy onset. However, these studies did not directly test how the age of onset relates to structural alterations, nor whether such associations differ by sex. In this context, our multisite analyses provide complementary neuroimaging evidence comparing age-of-onset effects in TLE and GGE across males and females.

Several methodological considerations should be acknowledged. First, although data harmonization was applied, site-specific information of subject motion and scanner effects may still influence the observed results. Second, all adult samples were analyzed collectively without stratification by age, age of onset, or disease duration, which could limit the detection of subtle subgroup-specific patterns. In addition, the use of the Desikan-Killiany parcellation, which provides relatively coarse spatial resolution, might have constrained the sensitivity to detect fine-grained cortical effects. Third, we did not include the stratification of GGE in this study. This may overlook syndrome-specific sex dimorphisms, as different GGE subtypes might involve distinct brain networks and pathophysiological mechanisms between genders^20,61^, which might contribute to the structural differences. Furthermore, our study lacked sex-specific endocrine variables, including menstrual cycle phase, menopausal status, hormonal treatment, catamenial seizure patterns, and circulating sex hormone levels, all of which may influence seizure susceptibility as suggested by previous studies^62–64^. This limitation is particularly relevant for the GGE cohort, where sex-stratified interaction models may be underpowered to detect small-to-moderate regional effects despite apparent descriptive differences. However, whether these neuroendocrine factors are associated with structural brain alterations in epilepsy, and whether these associations differ by sex remains unclear. Future studies are necessary to employ higher-resolution parcellations, longitudinal designs, subtype-specific stratification, and the integration of neuroendocrine or hormonal measures to capture sex-specific mechanisms of epilepsy-related brain alterations. Finally, we did not account for social and environmental factors that differ between males and females in this study, such as social stigma, quality of life, and healthcare access. These factors have been linked to the sex difference in psychopathology among epilepsy patients^25,65–67^. Of note, future work should further investigate whether such factors would influence clinical manifestations and structural brain alterations in epilepsy.

In summary, our findings indicate different alteration patterns of cortical thickness and subcortical volume in males and females with TLE and GGE in sex-stratified analyses. However, these differences were not supported by significant sex-by-diagnosis interactions, suggesting that epilepsy-related macroscale structural alterations do not differ between males and females. Moreover, disease duration and age of onset did not differentially affect male and female patients in either TLE or GGE. Instead, clinically relevant sex effects in epilepsy may be reflective of a combination of biological mechanisms, including endocrine regulation, neurotransmitter systems, or large-scale functional network dynamics that diverge between sexes.

## Methods

### Participants

The ENIGMA-Epilepsy consortium comprised 1,117 adults with epilepsy (465 males; mean age ± SD: 37.51 ± 11.04 years) and 536 healthy controls (226 males; mean age ± SD: 34.18 ± 10.88 years). Our main analyses focused on two patient subcohorts with site-matched healthy controls: The TLE cohort included 902 participants encompassing individuals with and without MTS (387 males; mean age ± SD: 38.50 ± 10.97 years) and 536 site-matched healthy controls (226 males; mean age ± SD: 34.18 ± 10.88 years). The GGE cohort comprised 215 adults (78 males; mean age ± SD: 33.36 ± 10.41 years) and 372 site-matched controls (153 males; mean age ± SD: 33.07 ± 10.41 years).

To increase statistical power and improve the robustness of sex-stratified and interaction analyses, we combined the ENIGMA-TLE sample with two independent TLE datasets and corresponding healthy controls: the Epilepsy Connectome Project (ECP)^68^ comprised 77 TLE participants (51 males, mean age ± SD: 38.45 ± 11.92 years) and 69 controls (38 males, mean age ± SD: 34.80 ± 10.84 years), the Montreal Neurological Institute (MNI) Hospital cohort^69^ included 59 TLE participants (29 males, mean age ± SD: 37.49 ± 11.63 years) and 69 controls (53 males, mean age ± SD: 32.13 ± 7.42 years).

All patients were diagnosed according to standardized International League Against Epilepsy (ILAE) criteria. Details of all participating cohorts’ demographic characteristics are summarized in **Supplementary Tables 1**.

### Neuroimaging data processing

Structural T1-weighted MRI scans were acquired across multiple sites in ENIGMA data, with scanner models and acquisition parameters detailed in the previous ENIGMA-Epilepsy study^6^. Acquisition details of the ECP and MNI datasets are listed in their previous studies^68,69^. Image processing for all datasets followed the standardized ENIGMA neuroimaging pipeline (https://enigma.ini.usc.edu/protocols/imaging-protocols/) and was processed within each site. For each participant, cortical thickness and subcortical volume measures were derived from their individual T1-weighted scans using FreeSurfer, with cortical parcellation based on the Desikan-Killiany atlas^70^. To combine the datasets while minimizing site and scanner variability in morphological measures, all data were harmonized using the ComBat toolbox (https://github.com/Jfortin1/ComBatHarmonization)^71^, which statistically adjusts for site effects while preserving age, sex, and diagnosis effects during data harmonization.

### Statistical analyses of cortical thickness and subcortical volume

Biological sex was assigned by clinicians based on clinical records. Demographic variables were compared between males and females. Continuous variables (e.g., age, age at onset, disease duration) were analyzed using two-sample t-tests, while categorical variables (e.g., side of seizure focus) were compared using Chi-square tests, with statistical significance set at *P* < 0.05. Group differences of each brain region were conducted using surface-based linear models implemented in the BrainStat toolbox (https://brainstat.readthedocs.io)^72^, applied to cortical thickness and subcortical volume measures derived from the neuroimaging preprocessing pipeline. To control for multiple comparisons across cortical vertices and subcortical regions, we applied false discovery rate (FDR) correction (*P*_FDR_ < 0.05). Analyses were performed separately for participants with TLE and GGE and their corresponding site-matched healthy controls.

### Effects of diagnosis, age of onset, and disease duration effects

To test the diagnostic effects across male and female participants, we first built linear models that included group and age as main effects, and compared individuals with TLE or GGE against their corresponding healthy control groups within males and females separately. Complementarily, within-diagnosis tests of sex differences were performed by employing linear models that included sex as a main effect to assess sex effects by comparing male and female participants within each diagnostic category (TLE, GGE, and controls). To further evaluate whether diagnostic effects differed between sexes, we modelled the diagnosis-by-sex interaction and controlled for the age, sex, and diagnosis effects in cortical thickness and subcortical volumes, capturing the differential impact of epilepsy versus control status across males and females. Intracranial volume was included as a covariate in all analyses of subcortical volumes.

We then applied linear models to assess the effects of duration of epilepsy and age of onset on cortical thickness and subcortical volume measurements in each patient cohort. Within sex analyses were conducted by fitting linear models separately for male and female participants within TLE and GGE. Linear models included disease duration or age of onset as the main effect. In addition, to test if sex will influence disease duration or age of onset effects in TLE and GGE, we built linear models that included sex, disease duration/age of onset, and their interaction term as main effects, by testing the significance of the interaction term. Intracranial volume was included as a covariate in all analyses of subcortical volumes.

### Sensitivity analyses

#### Robustness analyses of the side of focus, and TLE with or without lesion

To evaluate whether the observed sex-related effect sizes in TLE were influenced by seizure laterality, we examined diagnosis-by-sex, disease duration-by-sex, and age-of-onset-by-sex interaction effects separately in left and right TLE, for both cortical thickness and subcortical volume. Furthermore, to determine whether sex differences in TLE were driven by mesial temporal sclerosis (MTS), we assessed the same interaction effects in TLE with MTS and non-lesional TLE separately.

### Robustness analyses of sex differences in age effects

To evaluate whether age-related structural associations differ by sex in TLE and GGE, and to investigate if disease-related findings were confounded by different aging effects across males and females, we applied linear models to assess the effects of age on cortical thickness and subcortical volume measurements in TLE and GGE. Within sex analyses were conducted by fitting linear models separately for male and female participants within temporal lobe epilepsy (TLE) and genetic generalized epilepsy (GGE), with age included as the main effect. To test if sex will influence age effects differently in TLE and GGE, linear models were fitted in cortical thickness and subcortical volume, including sex, age, and their interaction term as main effects, and testing the significance of the interaction term.

### Consistency analyses across datasets

We repeated the main sex difference analyses across ENIGMA, ECP, and MNI datasets. Specifically, we re-evaluated the diagnosis-by-sex, disease duration-by-sex, and age of onset-by-sex interaction effects by using the same surface-based model as the merged dataset in TLE samples. These analyses provided a quantitative assessment of the cross-dataset reproducibility and robustness of the observed sex-related effects in epilepsy.

## Data availability

Summary statistics of neuroimaging data from the ENIGMA-Epilepsy meta-analysis are publicly available (https://github.com/MICA-MNI/ENIGMA). Access to subject-level neuroimaging data can be requested via the ENIGMA-Epilepsy consortium (enigma.ini.usc.edu/).

## Code availability

Our analysis code and visualization are openly available at GitHub repository (https://github.com/TaoNeuro/ENIGMA-epi-sex-diff). Statistical analyses were carried out using BrainStat (https://github.com/MICA-MNI/BrainStat). Visualizations were based on BrainSpace (v.0.1.2; https://brainspace.readthedocs.io/en/latest/)^73^ and ENIGMA toolbox (v.2.0.2; https://enigma-toolbox.readthedocs.io/en/latest/)^74^. All further analyses are mentioned in the corresponding “Methods” section.

## Acknowledgements

H.W. was funded by the German Federal Ministry of Education and Research (BMBF) and the Max Planck Society through SLV; T.G. was supported by NIH/NINDS K23NS135108; B.S. was funded by the German Federal Ministry of Education and Research (BMBF) and the Max Planck Society through SLV; M.D.H. was funded through a Walter Benjamin Fellowship of the German Research Foundation (HE 10277/1-1); T.B. was funded by the Deutsche Forschungsgemeinschaft (493623632) as Neuro-aCSis Clinician Scientist fellow; A.B. is supported by CIHR MOP-57840; N.B. is supported by CIHR MOP-57840; P.B. was supported by Ricerca Corrente 2025 from Italian Ministry of Health; F.C. was supported by São Paulo Research Foundation (FAPESP) Grant 2013/07559-3; L.C. was supported by Conahcyt (181508, 1782, CF-2023-I-218) and UNAM-DGAPA (IB101712, IG200117, IN213423). Imaging for UNAM cohort was performed at the National Laboratory for MRI; O.D. was supported by Finding A Cure for Epilepsy and Seizures (FACES); M.G. received funding from the Swiss National Science Foundation, Koetser Foundation for Brain Research, Herzog-Egli Foundation, and USZ Innovationspool; R.G. is supported by Current Research 2026, Ministry of Health for IRCCS, PNRR MNESYS project; M.I. is supported by Natural Sciences and Engineering Research Council of Canada (NSERC); S.I. was funded by the National Institute of Neurological Disorders and Stroke Intramural Research Program, National Institutes of Health; S.L. was supported by the Canadian Institutes of Health Research (CIHR), Centre de recherche du CHUS (CRCHUS), Université de Sherbrooke, the Natural Sciences and Engineering Research Council of Canada (NSERC: RGPIN-2025-06138), and the Canada Research Chairs program; M.Le. is supported by Current Research 2026, Ministry of Health for IRCCS; S.M. is supported by the 3TLE multicentric research project (Italian Ministry of Health, NET-2013-02355313); T.J.O. was supported by NHMRC Investigator Grants (APP1176426 & APP2034258); H.P. was supported by the Australian Epilepsy Project which received funding from the Australian Government under the Medical Research Future Fund (Frontier Health and Medical Research Program - Grant Number RFRHPSI000008); R.R.-C. is supported by the Mexican Council of Science and Technology (CONACYT 181508); UNAM-DGAPA (IB201712); J.R. was supported by the Banting Postdoctoral Fellowship; T.R. was supported by the German Ministry of Research, Technology and Space; P.S. was supported by Ministery of Health (Ricerca Corrente 2025); ENIGMA extends gratitude to the NIH Big Data to Knowledge (BD2K) award for its foundational support and contributions to consortium development (U54 EB020403 awarded to PMT). Research reported in this publication was supported by NIH S10OD032285. For a comprehensive list of ENIGMA-related grant support, please visit: https://enigma.ini.usc.edu/about-2/funding/. Sophia I. Thomopoulos was supported by R01NS122827; L.V. is supported by MRFF grants (MRF1172076, MRF1200254, MRF2023250) for unrelated projects; C.L.Y. was supported by CNPQ (403307/2021-0; 445340/2024-0; 315953/2021-7); FAPESP (CEPID BRAINN - 2013/07559-3); P.M.T. was supported by R01NS122827; S.S. was supported by the Epilepsy Society; C.M. is funded by R01 NS122827; B.C.B. acknowledges research support from the National Science and Engineering Research Council of Canada (NSERC RGPIN-2025-05932), CIHR (FDN-154298, PJT-174995, PJT-191853), Sick-Kids Foundation (NI17-039), Helmholtz International BigBrain Analytics and Learning Laboratory (HIBALL), Healthy Brains and Healthy Lives (HBHL), Brain Canada Foundation, FRQS, Tier-2 Canada Research Chairs Program, and The Centre for Excellence in Epilepsy at the Neuro (CEEN); S.L.V. was funded by the Max Planck Society through the Lise Meitner Excellence Program, the Jacobs Foundation Research Fellowship, the Hector Research Career Development Award, and the ERC Starting Grant. We would like to acknowledge the late Professor Dan J. Stein for his longstanding collaboration within ENIGMA-Epilepsy and for his invaluable contributions to psychiatry, neuroscience, and mental health research. His intellectual generosity, mentorship, and unwavering support shaped this work and the broader field in lasting ways. His scientific legacy and his kindness to colleagues continue to inspire us. The views expressed in this article are those of the authors and do not necessarily reflect the position or policy of any funding sources listed here.

