## Supplementary Material for "Sex-related structural alterations across common epilepsies: a worldwide ENIGMA study"

### Supplementary Information

**Supplementary Table 1.** Demographics and clinical details across three datasets
**Supplementary Table 2.** Abbreviation of cortical regions in Desikan-Killiany atlas
**Supplementary Table 3.** Effect size of diagnosis in cortical thickness within males and females
**Supplementary Table 4.** Effect size of diagnosis in subcortical volume within males and females
**Supplementary Table 5.** Effect size of sex in cortical thickness within epilepsy and corresponding healthy controls
**Supplementary Table 6.** Effect size of sex in subcortical volume within epilepsy and corresponding healthy controls
**Supplementary Table 7.** Effect size of disease duration in cortical thickness within male and female patients
**Supplementary Table 8.** Effect size of disease duration in subcortical volume within male and female patients
**Supplementary Table 9.** Effect size of age of onset in cortical thickness within male and female patients
**Supplementary Table 10.** Effect size of age of onset in subcortical volume within male and female patients
**Supplementary Table 11.** Summary of effect size of sex interaction effects in cortical thickness
**Supplementary Table 12.** Summary of effect size of sex interaction effects in subcortical volume
**Supplementary Table 13.** Summary of effect size of sex interaction effects in cortical thickness in left and right TLE
**Supplementary Table 14.** Summary of effect size of sex interaction effects in subcortical volume in left and right TLE
**Supplementary Table 15.** Summary of effect size of sex interaction effects in cortical thickness in MTS and non-lesional TLE patients
**Supplementary Table 16.** Summary of effect size of sex interaction effects in subcortical volume in MTS and non-lesional TLE patients
**Supplementary Table 17.** Effect size of age in cortical thickness in male and female patients with TLE and GGE
**Supplementary Table 18.** Effect size of age in subcortical volume in male and female patients with TLE and GGE

**Supplementary Table 19.** Summary of effect size of sex interaction effects in cortical thickness in TLE in ENIGMA dataset

**Supplementary Table 20.** Summary of effect size of sex interaction effects in subcortical volume in TLE in ENIGMA dataset

**Supplementary Table 21.** Summary of effect size of sex interaction effects in cortical thickness in TLE in ECP and MNI datasets

**Supplementary Table 22.** Summary of effect size of sex interaction effects in subcortical volume in TLE in ECP and MNI datasets

**Supplementary Figure 1.** Subcohort analyses of sex differences across left and right TLE subgroups

**Supplementary Figure 2.** Sex differences in mesial temporal sclerosis (MTS) and non-lesional TLE patients

**Supplementary Figure 3.** Sex differences in the effects of age in TLE and GGE

**Supplementary Figure 4.** Consistency of sex interaction effects analyses in TLE samples across datasets

**Supplementary Table 1.** Demographics and clinical details across three datasets

| | ENIGMA-Consortium | | | | ECP | | MNI | |
|---|---|---|---|---|---|---|---|---|
| **Cohorts** | TLE (n = 902) | TLE-HC (n = 536) | GGE (n = 215) | GGE-HC (n = 372) | TLE (n = 77) | HC (n = 69) | TLE (n = 59) | HC (n = 100) |
| **Age, years** | 38.50 ± 10.97 | 34.18 ± 10.88 | 33.36 ± 10.41 | 33.07 ± 10.41 | 38.45 ± 11.92 | 34.80 ± 10.84 | 37.49 ± 11.63 | 32.13 ± 7.42 |
| **Sex (M/F)** | 387/515 | 226/310 | 78/137 | 153/219 | 51/26 | 38/31 | 29/30 | 53/47 |
| **Side of focus (L/R)** | 516/386 | - | - | - | 57/20 | - | 37/22 | - |
| **Onset, years** | 16.86 ± 11.85 | - | 12.92 ± 7.04 | - | 22.95 ± 13.92 | - | 21.73 ± 13.29 | - |
| **Duration, years** | 21.63 ± 13.91 | - | 20.31 ± 11.55 | - | 15.47 ± 12.77 | - | 15.78 ± 11.47 | - |

Demographics are shown for patient subcohorts and their site-matched controls across three datasets. TLE: temporal lobe epilepsy; GGE: genetic generalized epilepsy; HC: healthy control; Onset: age of onset; Duration: disease duration; L = left; R = right.

**Supplementary Table 2.** Abbreviation of cortical regions in Desikan-Killiany atlas

| ROIs | Hemisphere | Abbreviation | Hemisphere | Abbreviation |
|---|---|---|---|---|
| **Banks of superior temporal sulcus** | Left | L_bankssts | Right | R_bankssts |
| **Caudal anterior cingulate** | Left | L_caudalanteriorcingulate | Right | R_caudalanteriorcingulate |
| **Caudal middle frontal gyrus** | Left | L_caudalmiddlefrontal | Right | R_caudalmiddlefrontal |
| **Cuneus** | Left | L_cuneus | Right | R_cuneus |
| **Entorhinal cortex** | Left | L_entorhinal | Right | R_entorhinal |
| **Fusiform gyrus** | Left | L_fusiform | Right | R_fusiform |
| **Inferior parietal cortex** | Left | L_inferiorparietal | Right | R_inferiorparietal |
| **Inferior temporal gyrus** | Left | L_inferiortemporal | Right | R_inferiortemporal |
| **Isthmus cingulate cortex** | Left | L_isthmuscingulate | Right | R_isthmuscingulate |
| **Lateral occipital cortex** | Left | L_lateraloccipital | Right | R_lateraloccipital |
| **Lateral orbitofrontal cortex** | Left | L_lateralorbitofrontal | Right | R_lateralorbitofrontal |
| **Lingual gyrus** | Left | L_lingual | Right | R_lingual |
| **Medial orbitofrontal** | Left | L_medialorbitofrontal | Right | R_medialorbitofrontal |
| **Middle temporal gyrus** | Left | L_middletemporal | Right | R_middletemporal |
| **Parahippocampal gyrus** | Left | L_parahippocampal | Right | R_parahippocampal |
| **Paracentral lobule** | Left | L_paracentral | Right | R_paracentral |
| **Pars opercularis** | Left | L_parsopercularis | Right | R_parsopercularis |
| **Pars orbitalis** | Left | L_parsorbitalis | Right | R_parsorbitalis |
| **Pars triangularis** | Left | L_parstriangularis | Right | R_parstriangularis |
| **Pericalcarine cortex** | Left | L_pericalcarine | Right | R_pericalcarine |
| **Postcentral gyrus** | Left | L_postcentral | Right | R_postcentral |
| **Posterior cingulate cortex** | Left | L_posteriorcingulate | Right | R_posteriorcingulate |
| **Precentral gyrus** | Left | L_precentral | Right | R_precentral |
| **Precuneus** | Left | L_precuneus | Right | R_precuneus |
| **Rostral anterior cingulate cortex** | Left | L_rostralanteriorcingulate | Right | R_rostralanteriorcingulate |
| **Rostral middle frontal gyrus** | Left | L_rostralmiddlefrontal | Right | R_rostralmiddlefrontal |
| **Superior frontal gyrus** | Left | L_superiorfrontal | Right | R_superiorfrontal |
| **Superior parietal cortex** | Left | L_superiorparietal | Right | R_superiorparietal |
| **Superior temporal gyrus** | Left | L_superiortemporal | Right | R_superiortemporal |
| **Supramarginal gyrus** | Left | L_supramarginal | Right | R_supramarginal |
| **Frontal pole** | Left | L_frontalpole | Right | R_frontalpole |
| **Temporal pole** | Left | L_temporalpole | Right | R_temporalpole |
| **Transverse temporal gyrus** | Left | L_transversetemporal | Right | R_transversetemporal |
| **Insula** | Left | L_insula | Right | R_insula |

**Supplementary Table 3.** Effect size of diagnosis in cortical thickness within males and females

| | TLE | | | | GGE | | | |
|---|---|---|---|---|---|---|---|---|
| | Male | | Female | | Male | | Female | |
| **Regions** | **d** | ***p* (FDR-corrected)** | **d** | ***p* (FDR-corrected)** | **d** | ***p* (FDR-corrected)** | **d** | ***p* (FDR-corrected)** |
| **L_bankssts** | -0.107 | 0.006 | -0.100 | 0.004 | -0.131 | 0.249 | -0.201 | 0.004 |
| **L_caudalanteriorcingulate** | 0.019 | 0.650 | 0.005 | 0.911 | -0.025 | 0.899 | -0.073 | 0.293 |
| **L_caudalmiddlefrontal** | -0.166 | < 0.001 | -0.140 | < 0.001 | -0.073 | 0.601 | -0.088 | 0.208 |
| **L_cuneus** | -0.166 | < 0.001 | -0.129 | < 0.001 | -0.050 | 0.728 | -0.067 | 0.329 |
| **L_entorhinal** | -0.031 | 0.449 | -0.102 | 0.003 | 0.126 | 0.275 | -0.022 | 0.727 |
| **L_fusiform** | -0.112 | 0.004 | -0.145 | < 0.001 | -0.117 | 0.320 | -0.109 | 0.114 |
| **L_inferiorparietal** | -0.170 | < 0.001 | -0.150 | < 0.001 | -0.100 | 0.415 | -0.216 | 0.002 |
| **L_inferiortemporal** | -0.044 | 0.289 | -0.100 | 0.004 | -0.074 | 0.601 | -0.147 | 0.037 |
| **L_isthmuscingulate** | -0.003 | 0.946 | -0.043 | 0.210 | 0.007 | 0.958 | -0.004 | 0.961 |
| **L_lateraloccipital** | -0.142 | < 0.001 | -0.166 | < 0.001 | -0.067 | 0.617 | -0.192 | 0.005 |
| **L_lateralorbitofrontal** | -0.064 | 0.120 | -0.072 | 0.035 | 0.058 | 0.675 | -0.063 | 0.356 |
| **L_lingual** | -0.176 | < 0.001 | -0.099 | 0.004 | -0.005 | 0.958 | 0.130 | 0.060 |
| **L_medialorbitofrontal** | -0.013 | 0.765 | -0.057 | 0.096 | 0.116 | 0.320 | 0.049 | 0.462 |
| **L_middletemporal** | -0.105 | 0.007 | -0.146 | < 0.001 | 0.003 | 0.958 | -0.157 | 0.023 |
| **L_parahippocampal** | -0.047 | 0.254 | -0.040 | 0.237 | 0.171 | 0.075 | 0.025 | 0.727 |
| **L_paracentral** | -0.193 | < 0.001 | -0.200 | < 0.001 | -0.104 | 0.393 | -0.131 | 0.058 |
| **L_parsopercularis** | -0.126 | 0.001 | -0.102 | 0.003 | -0.009 | 0.958 | -0.142 | 0.045 |
| **L_parsorbitalis** | -0.078 | 0.052 | -0.061 | 0.074 | -0.004 | 0.958 | -0.084 | 0.223 |
| **L_parstriangularis** | -0.072 | 0.078 | -0.099 | 0.004 | 0.078 | 0.601 | -0.125 | 0.072 |
| **L_pericalcarine** | -0.121 | 0.002 | -0.119 | < 0.001 | -0.042 | 0.814 | -0.036 | 0.604 |
| **L_postcentral** | -0.211 | < 0.001 | -0.142 | < 0.001 | -0.087 | 0.506 | -0.015 | 0.826 |
| **L_posteriorcingulate** | -0.026 | 0.525 | -0.084 | 0.014 | 0.056 | 0.680 | -0.103 | 0.122 |

| | | | | | | | | |
|---|---|---|---|---|---|---|---|---|
| **L_precentral** | -0.207 | < 0.001 | -0.220 | < 0.001 | -0.227 | 0.009 | -0.182 | 0.008 |
| **L_precuneus** | -0.185 | < 0.001 | -0.186 | < 0.001 | 0.023 | 0.899 | -0.024 | 0.727 |
| **L_rostralanteriorcingulate** | -0.053 | 0.209 | -0.002 | 0.970 | 0.023 | 0.899 | -0.103 | 0.122 |
| **L_rostralmiddlefrontal** | -0.040 | 0.323 | -0.091 | 0.008 | 0.211 | 0.015 | -0.012 | 0.857 |
| **L_superiorfrontal** | -0.129 | < 0.001 | -0.134 | < 0.001 | -0.026 | 0.899 | -0.073 | 0.293 |
| **L_superiorparietal** | -0.209 | < 0.001 | -0.171 | < 0.001 | -0.098 | 0.422 | -0.056 | 0.413 |
| **L_superiortemporal** | -0.109 | 0.005 | -0.101 | 0.004 | -0.071 | 0.601 | -0.165 | 0.015 |
| **L_supramarginal** | -0.136 | < 0.001 | -0.146 | < 0.001 | -0.007 | 0.958 | -0.070 | 0.304 |
| **L_frontalpole** | -0.006 | 0.883 | -0.057 | 0.094 | 0.054 | 0.684 | -0.072 | 0.293 |
| **L_temporalpole** | -0.069 | 0.093 | -0.099 | 0.004 | -0.033 | 0.849 | -0.042 | 0.540 |
| **L_transversetemporal** | -0.170 | < 0.001 | -0.128 | < 0.001 | -0.197 | 0.025 | -0.051 | 0.461 |
| **L_insula** | -0.001 | 0.970 | -0.012 | 0.744 | -0.009 | 0.958 | -0.138 | 0.049 |
| **R_bankssts** | -0.043 | 0.291 | -0.114 | 0.001 | -0.062 | 0.645 | -0.106 | 0.121 |
| **R_caudalanteriorcingulate** | -0.035 | 0.401 | -0.059 | 0.084 | -0.124 | 0.275 | -0.026 | 0.727 |
| **R_caudalmiddlefrontal** | -0.181 | < 0.001 | -0.174 | < 0.001 | -0.168 | 0.078 | -0.213 | 0.002 |
| **R_cuneus** | -0.148 | < 0.001 | -0.126 | < 0.001 | -0.146 | 0.158 | -0.036 | 0.604 |
| **R_entorhinal** | -0.049 | 0.240 | -0.071 | 0.038 | 0.243 | 0.009 | 0.000 | 0.995 |
| **R_fusiform** | -0.137 | < 0.001 | -0.106 | 0.002 | 0.055 | 0.680 | -0.050 | 0.462 |
| **R_inferiorparietal** | -0.151 | < 0.001 | -0.138 | < 0.001 | -0.012 | 0.958 | -0.033 | 0.634 |
| **R_inferiortemporal** | -0.092 | 0.020 | -0.062 | 0.073 | -0.035 | 0.837 | -0.133 | 0.058 |
| **R_isthmuscingulate** | -0.041 | 0.321 | -0.053 | 0.120 | 0.008 | 0.958 | -0.062 | 0.356 |
| **R_lateraloccipital** | -0.168 | < 0.001 | -0.183 | < 0.001 | -0.104 | 0.393 | -0.103 | 0.122 |
| **R_lateralorbitofrontal** | -0.081 | 0.045 | -0.044 | 0.200 | -0.036 | 0.837 | -0.084 | 0.223 |
| **R_lingual** | -0.165 | < 0.001 | -0.111 | 0.001 | -0.146 | 0.158 | -0.003 | 0.973 |
| **R_medialorbitofrontal** | -0.027 | 0.514 | -0.100 | 0.004 | -0.061 | 0.645 | -0.111 | 0.108 |
| **R_middletemporal** | -0.055 | 0.189 | -0.079 | 0.021 | 0.027 | 0.899 | -0.119 | 0.081 |
| **R_parahippocampal** | -0.010 | 0.818 | -0.085 | 0.013 | 0.067 | 0.617 | -0.170 | 0.012 |
| **R_paracentral** | -0.243 | < 0.001 | -0.223 | < 0.001 | -0.300 | < 0.001 | -0.176 | 0.010 |

| | | | | | | | | |
|---|---|---|---|---|---|---|---|---|
| **R_parsopercularis** | -0.116 | 0.003 | -0.118 | < 0.001 | -0.107 | 0.393 | -0.136 | 0.053 |
| **R_parsorbitalis** | -0.051 | 0.212 | -0.033 | 0.337 | -0.071 | 0.601 | -0.105 | 0.121 |
| **R_parstriangularis** | -0.056 | 0.183 | -0.113 | 0.001 | -0.038 | 0.834 | -0.063 | 0.356 |
| **R_pericalcarine** | -0.150 | < 0.001 | -0.114 | 0.001 | -0.213 | 0.015 | -0.058 | 0.397 |
| **R_postcentral** | -0.226 | < 0.001 | -0.121 | < 0.001 | -0.089 | 0.499 | -0.023 | 0.727 |
| **R_posteriorcingulate** | -0.044 | 0.284 | -0.107 | 0.002 | 0.020 | 0.925 | -0.118 | 0.083 |
| **R_precentral** | -0.210 | < 0.001 | -0.196 | < 0.001 | -0.229 | 0.009 | -0.189 | 0.005 |
| **R_precuneus** | -0.206 | < 0.001 | -0.149 | < 0.001 | 0.038 | 0.834 | -0.072 | 0.293 |
| **R_rostralanteriorcingulate** | 0.031 | 0.447 | 0.015 | 0.676 | -0.043 | 0.814 | -0.054 | 0.429 |
| **R_rostralmiddlefrontal** | -0.057 | 0.177 | -0.082 | 0.016 | 0.005 | 0.958 | -0.077 | 0.279 |
| **R_superiorfrontal** | -0.125 | 0.001 | -0.161 | < 0.001 | -0.062 | 0.645 | -0.122 | 0.075 |
| **R_superiorparietal** | -0.225 | < 0.001 | -0.187 | < 0.001 | -0.073 | 0.601 | -0.076 | 0.284 |
| **R_superiortemporal** | -0.060 | 0.149 | -0.076 | 0.026 | -0.025 | 0.899 | -0.101 | 0.127 |
| **R_supramarginal** | -0.147 | < 0.001 | -0.084 | 0.014 | -0.012 | 0.958 | 0.046 | 0.500 |
| **R_frontalpole** | -0.034 | 0.401 | 0.001 | 0.986 | 0.013 | 0.958 | -0.023 | 0.727 |
| **R_temporalpole** | -0.016 | 0.713 | -0.056 | 0.098 | -0.096 | 0.430 | -0.105 | 0.121 |
| **R_transversetemporal** | -0.113 | 0.003 | -0.086 | 0.012 | -0.229 | 0.009 | -0.087 | 0.208 |
| **R_insula** | -0.052 | 0.212 | 0.033 | 0.337 | 0.077 | 0.601 | -0.124 | 0.072 |

**Supplementary Table 4.** Effect size of diagnosis in subcortical volume within males and females

| | TLE | | | | GGE | | | |
|---|---|---|---|---|---|---|---|---|
| | Male | | Female | | Male | | Female | |
| **Regions** | **d** | ***p* (FDR-corrected)** | **d** | ***p* (FDR-corrected)** | **d** | ***p* (FDR-corrected)** | **d** | ***p* (FDR-corrected)** |
| **Left accumbens** | -0.136 | < 0.001 | -0.074 | 0.028 | -0.230 | 0.003 | -0.149 | 0.006 |
| **Left amygdala** | 0.054 | 0.131 | -0.045 | 0.181 | -0.084 | 0.218 | -0.159 | 0.003 |
| **Left caudate** | -0.138 | < 0.001 | -0.131 | < 0.001 | -0.134 | 0.087 | -0.229 | < 0.001 |
| **Left hippocampus** | -0.208 | < 0.001 | -0.282 | < 0.001 | -0.104 | 0.149 | -0.209 | < 0.001 |
| **Left pallidum** | -0.172 | < 0.001 | -0.161 | < 0.001 | -0.203 | 0.008 | -0.255 | < 0.001 |
| **Left putamen** | -0.148 | < 0.001 | -0.183 | < 0.001 | -0.106 | 0.149 | -0.199 | < 0.001 |
| **Left thalamus** | -0.236 | < 0.001 | -0.271 | < 0.001 | -0.225 | 0.003 | -0.235 | < 0.001 |
| **Right accumbens** | -0.116 | 0.001 | -0.072 | 0.029 | -0.174 | 0.024 | -0.206 | < 0.001 |
| **Right amygdala** | 0.067 | 0.066 | 0.003 | 0.921 | 0.003 | 0.969 | -0.110 | 0.038 |
| **Right caudate** | -0.123 | < 0.001 | -0.134 | < 0.001 | -0.134 | 0.087 | -0.247 | < 0.001 |
| **Right hippocampus** | -0.165 | < 0.001 | -0.258 | < 0.001 | -0.121 | 0.106 | -0.232 | < 0.001 |
| **Right pallidum** | -0.126 | < 0.001 | -0.158 | < 0.001 | -0.123 | 0.106 | -0.235 | < 0.001 |
| **Right putamen** | -0.134 | < 0.001 | -0.171 | < 0.001 | -0.090 | 0.199 | -0.198 | < 0.001 |
| **Right thalamus** | -0.254 | < 0.001 | -0.283 | < 0.001 | -0.257 | 0.002 | -0.254 | < 0.001 |

**Supplementary Table 5.** Effect size of sex in cortical thickness within epilepsy and corresponding healthy controls

| | TLE | | TLE-HC | | GGE | | GGE-HC | |
|---|---|---|---|---|---|---|---|---|
| **Regions** | **d** | ***p* (FDR-corrected)** | **d** | ***p* (FDR-corrected)** | **d** | ***p* (FDR-corrected)** | **d** | ***p* (FDR-corrected)** |
| **L_bankssts** | -0.007 | 0.951 | 0.002 | 0.987 | 0.124 | 0.184 | 0.060 | 0.463 |
| **L_caudalanteriorcingulate** | -0.037 | 0.499 | -0.046 | 0.531 | -0.019 | 0.858 | -0.074 | 0.420 |
| **L_caudalmiddlefrontal** | -0.016 | 0.835 | 0.024 | 0.790 | 0.017 | 0.858 | -0.001 | 0.982 |
| **L_cuneus** | 0.026 | 0.692 | 0.066 | 0.408 | 0.089 | 0.338 | 0.075 | 0.420 |
| **L_entorhinal** | 0.119 | 0.004 | 0.052 | 0.486 | 0.218 | 0.014 | 0.063 | 0.445 |
| **L_fusiform** | 0.057 | 0.230 | 0.027 | 0.790 | 0.065 | 0.469 | 0.072 | 0.420 |
| **L_inferiorparietal** | -0.034 | 0.530 | -0.008 | 0.946 | 0.163 | 0.063 | 0.050 | 0.503 |
| **L_inferiortemporal** | 0.076 | 0.142 | 0.024 | 0.790 | 0.159 | 0.070 | 0.075 | 0.420 |
| **L_isthmuscingulate** | 0.109 | 0.010 | 0.087 | 0.171 | 0.157 | 0.070 | 0.150 | 0.061 |
| **L_lateraloccipital** | 0.016 | 0.835 | 0.002 | 0.987 | 0.178 | 0.037 | 0.050 | 0.503 |
| **L_lateralorbitofrontal** | 0.053 | 0.262 | 0.039 | 0.623 | 0.187 | 0.029 | 0.054 | 0.472 |
| **L_lingual** | 0.056 | 0.240 | 0.137 | 0.006 | 0.016 | 0.858 | 0.150 | 0.061 |
| **L_medialorbitofrontal** | 0.058 | 0.230 | 0.022 | 0.801 | 0.106 | 0.277 | 0.054 | 0.472 |
| **L_middletemporal** | 0.061 | 0.230 | 0.044 | 0.540 | 0.233 | 0.011 | 0.072 | 0.420 |
| **L_parahippocampal** | -0.059 | 0.230 | -0.058 | 0.449 | 0.074 | 0.445 | -0.080 | 0.420 |
| **L_paracentral** | -0.024 | 0.720 | -0.018 | 0.840 | -0.014 | 0.858 | -0.040 | 0.616 |
| **L_parsopercularis** | 0.055 | 0.249 | 0.092 | 0.165 | 0.205 | 0.018 | 0.064 | 0.445 |
| **L_parsorbitalis** | 0.002 | 0.988 | 0.022 | 0.801 | 0.118 | 0.217 | 0.037 | 0.641 |
| **L_parstriangularis** | 0.009 | 0.935 | -0.009 | 0.946 | 0.157 | 0.070 | -0.050 | 0.503 |
| **L_pericalcarine** | 0.041 | 0.416 | 0.034 | 0.693 | 0.011 | 0.877 | 0.011 | 0.880 |
| **L_postcentral** | -0.058 | 0.230 | 0.005 | 0.946 | -0.043 | 0.657 | 0.027 | 0.723 |
| **L_posteriorcingulate** | 0.094 | 0.035 | 0.053 | 0.486 | 0.239 | 0.009 | 0.075 | 0.420 |
| **L_precentral** | 0.005 | 0.951 | 0.025 | 0.790 | -0.027 | 0.788 | 0.054 | 0.472 |

| | | | | | | | | |
|---|---|---|---|---|---|---|---|---|
| **L_precuneus** | 0.050 | 0.308 | 0.068 | 0.407 | 0.149 | 0.086 | 0.111 | 0.224 |
| **L_rostralanteriorcingulate** | 0.000 | 0.988 | 0.043 | 0.540 | 0.139 | 0.117 | 0.011 | 0.880 |
| **L_rostralmiddlefrontal** | 0.071 | 0.178 | 0.034 | 0.693 | 0.199 | 0.021 | -0.024 | 0.732 |
| **L_superiorfrontal** | -0.005 | 0.951 | 0.005 | 0.946 | 0.046 | 0.637 | -0.004 | 0.960 |
| **L_superiorparietal** | -0.007 | 0.951 | 0.055 | 0.486 | 0.093 | 0.324 | 0.134 | 0.085 |
| **L_superiortemporal** | 0.042 | 0.416 | 0.065 | 0.408 | 0.210 | 0.017 | 0.102 | 0.275 |
| **L_supramarginal** | -0.029 | 0.633 | -0.018 | 0.840 | 0.099 | 0.307 | 0.033 | 0.681 |
| **L_frontalpole** | 0.024 | 0.722 | -0.032 | 0.727 | 0.074 | 0.445 | -0.058 | 0.469 |
| **L_temporalpole** | 0.014 | 0.835 | -0.014 | 0.896 | 0.070 | 0.458 | 0.062 | 0.445 |
| **L_transversetemporal** | -0.047 | 0.347 | -0.019 | 0.837 | -0.115 | 0.223 | 0.025 | 0.732 |
| **L_insula** | 0.100 | 0.023 | 0.086 | 0.171 | 0.219 | 0.014 | 0.099 | 0.275 |
| **R_bankssts** | 0.076 | 0.142 | 0.013 | 0.901 | 0.101 | 0.294 | 0.057 | 0.469 |
| **R_caudalanteriorcingulate** | -0.001 | 0.988 | -0.026 | 0.790 | -0.072 | 0.449 | 0.027 | 0.723 |
| **R_caudalmiddlefrontal** | -0.016 | 0.835 | 0.008 | 0.946 | 0.098 | 0.307 | 0.041 | 0.614 |
| **R_cuneus** | 0.054 | 0.249 | 0.090 | 0.165 | 0.023 | 0.816 | 0.143 | 0.069 |
| **R_entorhinal** | 0.020 | 0.810 | 0.008 | 0.946 | 0.226 | 0.012 | 0.009 | 0.892 |
| **R_fusiform** | 0.014 | 0.835 | 0.058 | 0.449 | 0.202 | 0.019 | 0.097 | 0.277 |
| **R_inferiorparietal** | 0.019 | 0.810 | 0.035 | 0.693 | 0.089 | 0.338 | 0.070 | 0.425 |
| **R_inferiortemporal** | 0.030 | 0.602 | 0.063 | 0.411 | 0.187 | 0.029 | 0.088 | 0.381 |
| **R_isthmuscingulate** | 0.148 | < 0.001 | 0.143 | 0.006 | 0.239 | 0.009 | 0.177 | 0.024 |
| **R_lateraloccipital** | 0.033 | 0.537 | 0.022 | 0.801 | 0.062 | 0.491 | 0.066 | 0.445 |
| **R_lateralorbitofrontal** | -0.060 | 0.230 | -0.028 | 0.790 | 0.094 | 0.319 | 0.037 | 0.641 |
| **R_lingual** | 0.036 | 0.499 | 0.098 | 0.128 | -0.013 | 0.858 | 0.132 | 0.085 |
| **R_medialorbitofrontal** | 0.044 | 0.390 | -0.044 | 0.540 | 0.069 | 0.462 | 0.020 | 0.788 |
| **R_middletemporal** | 0.068 | 0.185 | 0.059 | 0.449 | 0.244 | 0.009 | 0.100 | 0.275 |
| **R_parahippocampal** | -0.067 | 0.185 | -0.139 | 0.006 | 0.065 | 0.469 | -0.180 | 0.024 |
| **R_paracentral** | -0.025 | 0.711 | 0.001 | 1.000 | -0.103 | 0.289 | 0.009 | 0.892 |
| **R_parsopercularis** | 0.043 | 0.411 | 0.052 | 0.486 | 0.114 | 0.228 | 0.086 | 0.390 |

| | | | | | | | | |
|---|---|---|---|---|---|---|---|---|
| **R_parsorbitalis** | -0.045 | 0.390 | -0.005 | 0.946 | 0.076 | 0.444 | 0.030 | 0.693 |
| **R_parstriangularis** | 0.022 | 0.764 | -0.026 | 0.790 | 0.067 | 0.464 | 0.040 | 0.616 |
| **R_pericalcarine** | 0.011 | 0.889 | 0.049 | 0.497 | -0.095 | 0.319 | 0.059 | 0.467 |
| **R_postcentral** | -0.059 | 0.230 | 0.048 | 0.513 | -0.015 | 0.858 | 0.055 | 0.472 |
| **R_posteriorcingulate** | 0.064 | 0.227 | 0.016 | 0.856 | 0.208 | 0.017 | 0.066 | 0.445 |
| **R_precentral** | -0.015 | 0.835 | 0.030 | 0.772 | 0.028 | 0.788 | 0.069 | 0.434 |
| **R_precuneus** | -0.006 | 0.951 | 0.074 | 0.339 | 0.180 | 0.036 | 0.081 | 0.420 |
| **R_rostralanteriorcingulate** | 0.015 | 0.835 | -0.006 | 0.946 | 0.045 | 0.637 | 0.030 | 0.693 |
| **R_rostralmiddlefrontal** | 0.011 | 0.889 | 0.000 | 1.000 | 0.154 | 0.073 | 0.064 | 0.445 |
| **R_superiorfrontal** | -0.017 | 0.835 | -0.045 | 0.540 | 0.036 | 0.707 | -0.031 | 0.693 |
| **R_superiorparietal** | 0.001 | 0.988 | 0.054 | 0.486 | 0.078 | 0.435 | 0.077 | 0.420 |
| **R_superiortemporal** | 0.070 | 0.178 | 0.068 | 0.407 | 0.196 | 0.021 | 0.138 | 0.077 |
| **R_supramarginal** | -0.007 | 0.951 | 0.063 | 0.411 | 0.041 | 0.657 | 0.106 | 0.256 |
| **R_frontalpole** | -0.041 | 0.416 | -0.012 | 0.908 | 0.056 | 0.537 | 0.018 | 0.793 |
| **R_temporalpole** | -0.002 | 0.988 | -0.051 | 0.489 | 0.041 | 0.657 | 0.024 | 0.732 |
| **R_transversetemporal** | -0.001 | 0.988 | 0.021 | 0.803 | -0.067 | 0.464 | 0.077 | 0.420 |
| **R_insula** | 0.035 | 0.519 | 0.128 | 0.013 | 0.334 | < 0.001 | 0.148 | 0.061 |

**Supplementary Table 6.** Effect size of sex in subcortical volume within epilepsy and corresponding healthy controls

| | TLE | | TLE-HC | | GGE | | GGE-HC | |
|---|---|---|---|---|---|---|---|---|
| **Regions** | **d** | ***p* (FDR-corrected)** | **d** | ***p* (FDR-corrected)** | **d** | ***p* (FDR-corrected)** | **d** | ***p* (FDR-corrected)** |
| **Left accumbens** | 0.148 | < 0.001 | 0.254 | < 0.001 | 0.122 | 0.087 | 0.176 | 0.002 |
| **Left amygdala** | 0.209 | < 0.001 | 0.198 | < 0.001 | 0.375 | < 0.001 | 0.187 | 0.002 |
| **Left caudate** | 0.088 | 0.005 | 0.172 | < 0.001 | 0.075 | 0.275 | 0.043 | 0.440 |
| **Left hippocampus** | 0.081 | 0.009 | 0.128 | < 0.001 | 0.297 | < 0.001 | 0.105 | 0.057 |
| **Left pallidum** | 0.183 | < 0.001 | 0.280 | < 0.001 | 0.221 | 0.002 | 0.134 | 0.018 |
| **Left putamen** | 0.211 | < 0.001 | 0.234 | < 0.001 | 0.248 | < 0.001 | 0.164 | 0.003 |
| **Left thalamus** | 0.162 | < 0.001 | 0.200 | < 0.001 | 0.243 | < 0.001 | 0.164 | 0.003 |
| **Right accumbens** | 0.140 | < 0.001 | 0.205 | < 0.001 | 0.254 | < 0.001 | 0.118 | 0.036 |
| **Right amygdala** | 0.194 | < 0.001 | 0.199 | < 0.001 | 0.385 | < 0.001 | 0.178 | 0.002 |
| **Right caudate** | 0.117 | < 0.001 | 0.160 | < 0.001 | 0.114 | 0.105 | 0.038 | 0.459 |
| **Right hippocampus** | 0.104 | < 0.001 | 0.121 | 0.001 | 0.259 | < 0.001 | 0.088 | 0.105 |
| **Right pallidum** | 0.192 | < 0.001 | 0.233 | < 0.001 | 0.250 | < 0.001 | 0.106 | 0.057 |
| **Right putamen** | 0.229 | < 0.001 | 0.244 | < 0.001 | 0.299 | < 0.001 | 0.187 | 0.002 |
| **Right thalamus** | 0.189 | < 0.001 | 0.255 | < 0.001 | 0.270 | < 0.001 | 0.210 | < 0.001 |

**Supplementary Table 7.** Effect size of disease duration in cortical thickness within male and female patients

| | TLE | | | | GGE | | | |
|---|---|---|---|---|---|---|---|---|
| | Male | | Female | | Male | | Female | |
| **Regions** | **d** | ***p* (FDR-corrected)** | **d** | ***p* (FDR-corrected)** | **d** | ***p* (FDR-corrected)** | **d** | ***p* (FDR-corrected)** |
| **L_bankssts** | -0.186 | < 0.001 | -0.140 | 0.002 | -0.373 | 0.051 | -0.202 | 0.076 |
| **L_caudalanteriorcingulate** | -0.073 | 0.138 | -0.009 | 0.851 | 0.040 | 0.848 | -0.175 | 0.120 |
| **L_caudalmiddlefrontal** | -0.296 | < 0.001 | -0.200 | < 0.001 | -0.266 | 0.130 | -0.299 | 0.011 |
| **L_cuneus** | -0.101 | 0.042 | -0.173 | < 0.001 | -0.041 | 0.848 | -0.231 | 0.043 |
| **L_entorhinal** | -0.101 | 0.042 | 0.025 | 0.595 | -0.208 | 0.287 | -0.075 | 0.499 |
| **L_fusiform** | -0.150 | 0.003 | -0.085 | 0.057 | -0.324 | 0.075 | -0.189 | 0.098 |
| **L_inferiorparietal** | -0.207 | < 0.001 | -0.187 | < 0.001 | -0.086 | 0.726 | -0.158 | 0.149 |
| **L_inferiortemporal** | -0.085 | 0.088 | -0.109 | 0.015 | -0.326 | 0.075 | -0.061 | 0.598 |
| **L_isthmuscingulate** | -0.217 | < 0.001 | -0.115 | 0.011 | -0.577 | 0.004 | -0.269 | 0.019 |
| **L_lateraloccipital** | -0.118 | 0.019 | -0.133 | 0.004 | -0.005 | 0.973 | -0.059 | 0.604 |
| **L_lateralorbitofrontal** | -0.140 | 0.005 | -0.064 | 0.153 | -0.047 | 0.848 | -0.177 | 0.115 |
| **L_lingual** | -0.135 | 0.007 | -0.165 | < 0.001 | -0.072 | 0.773 | -0.221 | 0.053 |
| **L_medialorbitofrontal** | -0.033 | 0.513 | -0.013 | 0.790 | -0.005 | 0.973 | -0.139 | 0.189 |
| **L_middletemporal** | -0.236 | < 0.001 | -0.121 | 0.008 | -0.419 | 0.044 | -0.320 | 0.007 |
| **L_parahippocampal** | -0.005 | 0.930 | -0.098 | 0.028 | 0.022 | 0.906 | 0.146 | 0.168 |
| **L_paracentral** | -0.167 | < 0.001 | -0.161 | < 0.001 | -0.076 | 0.773 | -0.129 | 0.224 |
| **L_parsopercularis** | -0.310 | < 0.001 | -0.248 | < 0.001 | -0.367 | 0.051 | -0.216 | 0.054 |
| **L_parsorbitalis** | -0.188 | < 0.001 | -0.097 | 0.031 | 0.056 | 0.834 | -0.127 | 0.224 |
| **L_parstriangularis** | -0.300 | < 0.001 | -0.119 | 0.008 | -0.344 | 0.063 | -0.296 | 0.011 |
| **L_pericalcarine** | -0.077 | 0.119 | -0.134 | 0.003 | 0.029 | 0.906 | -0.099 | 0.347 |
| **L_postcentral** | -0.139 | 0.005 | -0.144 | 0.002 | -0.176 | 0.377 | -0.148 | 0.167 |
| **L_posteriorcingulate** | -0.167 | < 0.001 | -0.129 | 0.004 | -0.148 | 0.471 | -0.263 | 0.021 |

| | | | | | | | | |
|---|---|---|---|---|---|---|---|---|
| **L_precentral** | -0.235 | < 0.001 | -0.225 | < 0.001 | -0.358 | 0.051 | -0.218 | 0.054 |
| **L_precuneus** | -0.205 | < 0.001 | -0.163 | < 0.001 | -0.094 | 0.714 | -0.180 | 0.113 |
| **L_rostralanteriorcingulate** | -0.037 | 0.458 | -0.016 | 0.742 | -0.174 | 0.377 | -0.271 | 0.019 |
| **L_rostralmiddlefrontal** | -0.227 | < 0.001 | -0.144 | 0.002 | -0.294 | 0.090 | -0.192 | 0.093 |
| **L_superiorfrontal** | -0.239 | < 0.001 | -0.156 | < 0.001 | -0.292 | 0.090 | -0.442 | < 0.001 |
| **L_superiorparietal** | -0.174 | < 0.001 | -0.159 | < 0.001 | 0.025 | 0.906 | -0.127 | 0.224 |
| **L_superiortemporal** | -0.225 | < 0.001 | -0.212 | < 0.001 | -0.362 | 0.051 | -0.319 | 0.007 |
| **L_supramarginal** | -0.254 | < 0.001 | -0.152 | 0.001 | -0.219 | 0.251 | -0.296 | 0.011 |
| **L_frontalpole** | -0.017 | 0.745 | -0.007 | 0.875 | 0.072 | 0.773 | 0.123 | 0.239 |
| **L_temporalpole** | -0.107 | 0.033 | -0.051 | 0.256 | -0.181 | 0.377 | -0.181 | 0.113 |
| **L_transversetemporal** | -0.216 | < 0.001 | -0.194 | < 0.001 | -0.302 | 0.090 | -0.220 | 0.053 |
| **L_insula** | -0.140 | 0.005 | -0.094 | 0.035 | -0.366 | 0.051 | -0.337 | 0.006 |
| **R_bankssts** | -0.195 | < 0.001 | -0.151 | 0.001 | -0.337 | 0.066 | -0.238 | 0.038 |
| **R_caudalanteriorcingulate** | -0.079 | 0.110 | -0.044 | 0.339 | -0.095 | 0.714 | -0.163 | 0.143 |
| **R_caudalmiddlefrontal** | -0.249 | < 0.001 | -0.137 | 0.003 | -0.101 | 0.712 | -0.236 | 0.038 |
| **R_cuneus** | -0.123 | 0.014 | -0.155 | < 0.001 | 0.120 | 0.628 | -0.169 | 0.129 |
| **R_entorhinal** | -0.116 | 0.020 | 0.034 | 0.469 | -0.015 | 0.939 | -0.034 | 0.747 |
| **R_fusiform** | -0.119 | 0.017 | -0.129 | 0.004 | 0.046 | 0.848 | -0.157 | 0.149 |
| **R_inferiorparietal** | -0.202 | < 0.001 | -0.213 | < 0.001 | -0.189 | 0.349 | -0.179 | 0.114 |
| **R_inferiortemporal** | -0.061 | 0.214 | -0.127 | 0.005 | -0.166 | 0.398 | -0.049 | 0.653 |
| **R_isthmuscingulate** | -0.226 | < 0.001 | -0.111 | 0.014 | -0.541 | 0.005 | -0.319 | 0.007 |
| **R_lateraloccipital** | -0.071 | 0.149 | -0.199 | < 0.001 | -0.044 | 0.848 | -0.025 | 0.818 |
| **R_lateralorbitofrontal** | -0.080 | 0.108 | -0.081 | 0.067 | -0.166 | 0.398 | -0.151 | 0.167 |
| **R_lingual** | -0.130 | 0.010 | -0.162 | < 0.001 | -0.098 | 0.714 | -0.292 | 0.011 |
| **R_medialorbitofrontal** | -0.060 | 0.222 | -0.084 | 0.057 | -0.250 | 0.162 | -0.051 | 0.645 |
| **R_middletemporal** | -0.174 | < 0.001 | -0.111 | 0.014 | -0.293 | 0.090 | -0.240 | 0.038 |
| **R_parahippocampal** | -0.103 | 0.038 | -0.106 | 0.018 | 0.057 | 0.834 | 0.107 | 0.310 |
| **R_paracentral** | -0.096 | 0.052 | -0.221 | < 0.001 | -0.109 | 0.688 | -0.149 | 0.167 |

| | | | | | | | | |
|---|---|---|---|---|---|---|---|---|
| **R_parsopercularis** | -0.273 | < 0.001 | -0.184 | < 0.001 | -0.312 | 0.084 | -0.270 | 0.019 |
| **R_parsorbitalis** | -0.248 | < 0.001 | -0.103 | 0.022 | -0.090 | 0.725 | -0.052 | 0.644 |
| **R_parstriangularis** | -0.166 | < 0.001 | -0.181 | < 0.001 | -0.292 | 0.090 | -0.216 | 0.054 |
| **R_pericalcarine** | -0.111 | 0.026 | -0.096 | 0.031 | 0.085 | 0.726 | -0.170 | 0.129 |
| **R_postcentral** | -0.113 | 0.023 | -0.133 | 0.003 | -0.027 | 0.906 | 0.004 | 0.974 |
| **R_posteriorcingulate** | -0.103 | 0.038 | -0.121 | 0.008 | -0.305 | 0.089 | -0.375 | 0.002 |
| **R_precentral** | -0.242 | < 0.001 | -0.213 | < 0.001 | -0.392 | 0.051 | -0.162 | 0.143 |
| **R_precuneus** | -0.166 | < 0.001 | -0.235 | < 0.001 | -0.315 | 0.083 | -0.272 | 0.019 |
| **R_rostralanteriorcingulate** | -0.077 | 0.119 | -0.091 | 0.041 | -0.157 | 0.429 | -0.372 | 0.002 |
| **R_rostralmiddlefrontal** | -0.198 | < 0.001 | -0.138 | 0.003 | -0.045 | 0.848 | -0.159 | 0.149 |
| **R_superiorfrontal** | -0.254 | < 0.001 | -0.150 | 0.001 | -0.174 | 0.377 | -0.401 | 0.001 |
| **R_superiorparietal** | -0.160 | 0.001 | -0.211 | < 0.001 | -0.067 | 0.787 | -0.035 | 0.747 |
| **R_superiortemporal** | -0.206 | < 0.001 | -0.156 | < 0.001 | -0.419 | 0.044 | -0.321 | 0.007 |
| **R_supramarginal** | -0.231 | < 0.001 | -0.145 | 0.002 | -0.360 | 0.051 | -0.260 | 0.022 |
| **R_frontalpole** | -0.021 | 0.677 | -0.019 | 0.692 | 0.105 | 0.701 | 0.003 | 0.974 |
| **R_temporalpole** | 0.004 | 0.930 | -0.070 | 0.116 | -0.199 | 0.313 | -0.144 | 0.172 |
| **R_transversetemporal** | -0.176 | < 0.001 | -0.080 | 0.070 | -0.269 | 0.127 | -0.148 | 0.167 |
| **R_insula** | -0.139 | 0.005 | -0.108 | 0.016 | -0.387 | 0.051 | -0.301 | 0.011 |

**Supplementary Table 8.** Effect size of disease duration in subcortical volume within male and female patients

| | TLE | | | | GGE | | | |
|---|---|---|---|---|---|---|---|---|
| | Male | | Female | | Male | | Female | |
| **Regions** | **d** | ***p* (FDR-corrected)** | **d** | ***p* (FDR-corrected)** | **d** | ***p* (FDR-corrected)** | **d** | ***p* (FDR-corrected)** |
| **Left accumbens** | -0.288 | < 0.001 | -0.239 | < 0.001 | -0.273 | 0.052 | -0.387 | < 0.001 |
| **Left amygdala** | -0.189 | < 0.001 | -0.158 | < 0.001 | -0.229 | 0.095 | -0.346 | < 0.001 |
| **Left caudate** | -0.219 | < 0.001 | -0.187 | < 0.001 | -0.474 | 0.002 | -0.628 | < 0.001 |
| **Left hippocampus** | -0.277 | < 0.001 | -0.242 | < 0.001 | -0.403 | 0.006 | -0.349 | < 0.001 |
| **Left pallidum** | -0.127 | 0.008 | -0.141 | 0.001 | -0.420 | 0.004 | -0.334 | < 0.001 |
| **Left putamen** | -0.243 | < 0.001 | -0.247 | < 0.001 | -0.609 | < 0.001 | -0.577 | < 0.001 |
| **Left thalamus** | -0.349 | < 0.001 | -0.344 | < 0.001 | -0.771 | < 0.001 | -0.496 | < 0.001 |
| **Right accumbens** | -0.212 | < 0.001 | -0.185 | < 0.001 | -0.436 | 0.004 | -0.351 | < 0.001 |
| **Right amygdala** | -0.101 | 0.032 | -0.150 | < 0.001 | -0.292 | 0.040 | -0.385 | < 0.001 |
| **Right caudate** | -0.201 | < 0.001 | -0.179 | < 0.001 | -0.445 | 0.003 | -0.625 | < 0.001 |
| **Right hippocampus** | -0.154 | 0.001 | -0.254 | < 0.001 | -0.225 | 0.095 | -0.255 | 0.008 |
| **Right pallidum** | -0.114 | 0.017 | -0.109 | 0.010 | -0.377 | 0.009 | -0.360 | < 0.001 |
| **Right putamen** | -0.226 | < 0.001 | -0.243 | < 0.001 | -0.621 | < 0.001 | -0.532 | < 0.001 |
| **Right thalamus** | -0.313 | < 0.001 | -0.311 | < 0.001 | -0.692 | < 0.001 | -0.483 | < 0.001 |

**Supplementary Table 9.** Effect size of age of onset in cortical thickness within male and female patients

| | TLE | | | | GGE | | | |
|---|---|---|---|---|---|---|---|---|
| | Male | | Female | | Male | | Female | |
| **Regions** | **d** | ***p* (FDR-corrected)** | **d** | ***p* (FDR-corrected)** | **d** | ***p* (FDR-corrected)** | **d** | ***p* (FDR-corrected)** |
| **L_bankssts** | -0.023 | 0.618 | -0.046 | 0.413 | -0.211 | 0.508 | -0.081 | 0.899 |
| **L_caudalanteriorcingulate** | -0.111 | 0.051 | -0.052 | 0.344 | -0.069 | 0.813 | 0.177 | 0.582 |
| **L_caudalmiddlefrontal** | -0.090 | 0.114 | -0.104 | 0.043 | -0.179 | 0.542 | 0.019 | 0.899 |
| **L_cuneus** | -0.069 | 0.186 | 0.027 | 0.688 | -0.166 | 0.544 | -0.026 | 0.899 |
| **L_entorhinal** | 0.089 | 0.114 | 0.024 | 0.688 | 0.199 | 0.508 | -0.068 | 0.899 |
| **L_fusiform** | -0.043 | 0.386 | -0.047 | 0.405 | -0.071 | 0.813 | 0.045 | 0.899 |
| **L_inferiorparietal** | -0.128 | 0.034 | -0.069 | 0.180 | -0.136 | 0.598 | -0.019 | 0.899 |
| **L_inferiortemporal** | -0.124 | 0.034 | -0.038 | 0.524 | -0.027 | 0.935 | -0.018 | 0.899 |
| **L_isthmuscingulate** | -0.088 | 0.114 | -0.138 | 0.009 | -0.024 | 0.935 | 0.053 | 0.899 |
| **L_lateraloccipital** | -0.088 | 0.114 | 0.007 | 0.927 | -0.118 | 0.640 | -0.198 | 0.582 |
| **L_lateralorbitofrontal** | -0.079 | 0.144 | -0.174 | < 0.001 | -0.152 | 0.554 | -0.018 | 0.899 |
| **L_lingual** | -0.098 | 0.079 | -0.071 | 0.168 | -0.293 | 0.265 | 0.059 | 0.899 |
| **L_medialorbitofrontal** | -0.104 | 0.065 | -0.188 | < 0.001 | -0.142 | 0.578 | 0.158 | 0.582 |
| **L_middletemporal** | -0.127 | 0.034 | -0.106 | 0.040 | -0.124 | 0.615 | 0.097 | 0.899 |
| **L_parahippocampal** | -0.056 | 0.261 | -0.006 | 0.927 | 0.002 | 0.989 | -0.070 | 0.899 |
| **L_paracentral** | -0.077 | 0.148 | 0.023 | 0.688 | -0.236 | 0.422 | -0.122 | 0.833 |
| **L_parsopercularis** | -0.086 | 0.114 | -0.111 | 0.032 | -0.095 | 0.756 | 0.046 | 0.899 |
| **L_parsorbitalis** | -0.053 | 0.284 | -0.100 | 0.049 | -0.145 | 0.575 | 0.017 | 0.899 |
| **L_parstriangularis** | -0.058 | 0.246 | -0.135 | 0.009 | -0.129 | 0.615 | 0.045 | 0.899 |
| **L_pericalcarine** | -0.077 | 0.148 | -0.023 | 0.688 | -0.294 | 0.265 | -0.025 | 0.899 |
| **L_postcentral** | -0.067 | 0.195 | -0.071 | 0.171 | -0.271 | 0.305 | -0.022 | 0.899 |
| **L_posteriorcingulate** | -0.201 | 0.001 | -0.112 | 0.032 | -0.039 | 0.935 | 0.133 | 0.829 |

| | | | | | | | | |
|---|---|---|---|---|---|---|---|---|
| **L_precentral** | -0.086 | 0.114 | -0.026 | 0.688 | -0.177 | 0.542 | -0.081 | 0.899 |
| **L_precuneus** | -0.114 | 0.045 | -0.076 | 0.139 | -0.151 | 0.554 | -0.071 | 0.899 |
| **L_rostralanteriorcingulate** | -0.105 | 0.065 | -0.145 | 0.006 | -0.078 | 0.813 | 0.223 | 0.582 |
| **L_rostralmiddlefrontal** | -0.124 | 0.034 | -0.116 | 0.025 | -0.275 | 0.305 | 0.085 | 0.899 |
| **L_superiorfrontal** | -0.141 | 0.024 | -0.175 | < 0.001 | -0.248 | 0.378 | 0.031 | 0.899 |
| **L_superiorparietal** | -0.127 | 0.034 | -0.003 | 0.949 | -0.194 | 0.508 | -0.096 | 0.899 |
| **L_superiortemporal** | -0.085 | 0.114 | -0.037 | 0.525 | -0.314 | 0.265 | 0.029 | 0.899 |
| **L_supramarginal** | -0.117 | 0.043 | -0.086 | 0.092 | -0.072 | 0.813 | 0.024 | 0.899 |
| **L_frontalpole** | -0.059 | 0.241 | -0.149 | 0.006 | -0.017 | 0.935 | -0.041 | 0.899 |
| **L_temporalpole** | 0.078 | 0.148 | 0.086 | 0.092 | -0.108 | 0.694 | 0.184 | 0.582 |
| **L_transversetemporal** | -0.045 | 0.364 | -0.057 | 0.282 | -0.319 | 0.265 | 0.048 | 0.899 |
| **L_insula** | -0.103 | 0.065 | -0.181 | < 0.001 | -0.175 | 0.542 | 0.124 | 0.833 |
| **R_bankssts** | -0.070 | 0.183 | -0.079 | 0.125 | -0.194 | 0.508 | -0.163 | 0.582 |
| **R_caudalanteriorcingulate** | -0.105 | 0.065 | -0.074 | 0.156 | 0.024 | 0.935 | 0.090 | 0.899 |
| **R_caudalmiddlefrontal** | -0.085 | 0.114 | -0.105 | 0.040 | -0.172 | 0.542 | 0.021 | 0.899 |
| **R_cuneus** | -0.061 | 0.229 | -0.019 | 0.737 | -0.182 | 0.542 | -0.088 | 0.899 |
| **R_entorhinal** | 0.064 | 0.215 | 0.020 | 0.719 | -0.045 | 0.935 | -0.101 | 0.899 |
| **R_fusiform** | -0.116 | 0.043 | -0.038 | 0.524 | -0.132 | 0.608 | 0.012 | 0.909 |
| **R_inferiorparietal** | -0.141 | 0.024 | -0.068 | 0.188 | -0.021 | 0.935 | -0.185 | 0.582 |
| **R_inferiortemporal** | -0.174 | 0.008 | -0.011 | 0.859 | -0.125 | 0.615 | -0.033 | 0.899 |
| **R_isthmuscingulate** | -0.041 | 0.412 | -0.135 | 0.009 | 0.068 | 0.813 | 0.014 | 0.904 |
| **R_lateraloccipital** | -0.143 | 0.024 | 0.022 | 0.688 | -0.016 | 0.935 | -0.161 | 0.582 |
| **R_lateralorbitofrontal** | -0.117 | 0.043 | -0.101 | 0.049 | -0.028 | 0.935 | -0.044 | 0.899 |
| **R_lingual** | -0.144 | 0.024 | -0.090 | 0.076 | -0.197 | 0.508 | 0.029 | 0.899 |
| **R_medialorbitofrontal** | -0.124 | 0.034 | -0.124 | 0.017 | -0.040 | 0.935 | -0.042 | 0.899 |
| **R_middletemporal** | -0.146 | 0.024 | -0.135 | 0.009 | -0.324 | 0.265 | 0.038 | 0.899 |
| **R_parahippocampal** | -0.076 | 0.148 | 0.000 | 0.994 | -0.028 | 0.935 | 0.031 | 0.899 |
| **R_paracentral** | -0.062 | 0.224 | -0.005 | 0.942 | -0.159 | 0.554 | -0.062 | 0.899 |

| | | | | | | | | |
|---|---|---|---|---|---|---|---|---|
| **R_parsopercularis** | -0.083 | 0.127 | -0.100 | 0.049 | 0.002 | 0.989 | 0.122 | 0.833 |
| **R_parsorbitalis** | -0.036 | 0.459 | -0.065 | 0.207 | -0.156 | 0.554 | -0.001 | 0.987 |
| **R_parstriangularis** | -0.152 | 0.024 | -0.121 | 0.020 | -0.035 | 0.935 | 0.057 | 0.899 |
| **R_pericalcarine** | -0.027 | 0.586 | -0.027 | 0.688 | -0.200 | 0.508 | 0.066 | 0.899 |
| **R_postcentral** | -0.087 | 0.114 | -0.064 | 0.214 | -0.266 | 0.305 | -0.163 | 0.582 |
| **R_posteriorcingulate** | -0.100 | 0.075 | -0.144 | 0.006 | -0.022 | 0.935 | 0.149 | 0.646 |
| **R_precentral** | -0.076 | 0.148 | -0.023 | 0.688 | -0.155 | 0.554 | -0.081 | 0.899 |
| **R_precuneus** | -0.071 | 0.178 | -0.029 | 0.678 | 0.105 | 0.697 | -0.058 | 0.899 |
| **R_rostralanteriorcingulate** | -0.104 | 0.065 | -0.091 | 0.076 | -0.088 | 0.783 | 0.305 | 0.100 |
| **R_rostralmiddlefrontal** | -0.103 | 0.065 | -0.090 | 0.076 | -0.326 | 0.265 | 0.105 | 0.899 |
| **R_superiorfrontal** | -0.135 | 0.028 | -0.131 | 0.011 | -0.230 | 0.427 | 0.067 | 0.899 |
| **R_superiorparietal** | -0.114 | 0.045 | 0.013 | 0.834 | -0.078 | 0.813 | -0.166 | 0.582 |
| **R_superiortemporal** | -0.119 | 0.043 | -0.095 | 0.061 | -0.306 | 0.265 | 0.058 | 0.899 |
| **R_supramarginal** | -0.135 | 0.028 | -0.116 | 0.025 | 0.016 | 0.935 | -0.091 | 0.899 |
| **R_frontalpole** | -0.073 | 0.165 | -0.108 | 0.037 | -0.068 | 0.813 | -0.047 | 0.899 |
| **R_temporalpole** | -0.065 | 0.213 | 0.024 | 0.688 | 0.035 | 0.935 | 0.054 | 0.899 |
| **R_transversetemporal** | -0.023 | 0.618 | -0.084 | 0.099 | -0.299 | 0.265 | -0.073 | 0.899 |
| **R_insula** | -0.125 | 0.034 | -0.147 | 0.006 | -0.169 | 0.544 | 0.107 | 0.899 |

**Supplementary Table 10.** Effect size of age of onset in subcortical volume within male and female patients

| | TLE | | | | GGE | | | |
|---|---|---|---|---|---|---|---|---|
| | Male | | Female | | Male | | Female | |
| **Regions** | **d** | ***p* (FDR-corrected)** | **d** | ***p* (FDR-corrected)** | **d** | ***p* (FDR-corrected)** | **d** | ***p* (FDR-corrected)** |
| **Left accumbens** | -0.042 | 0.894 | -0.048 | 0.365 | -0.006 | 0.990 | 0.117 | 0.824 |
| **Left amygdala** | 0.022 | 0.894 | 0.057 | 0.344 | 0.037 | 0.990 | 0.062 | 0.824 |
| **Left caudate** | -0.046 | 0.894 | -0.096 | 0.083 | 0.083 | 0.990 | 0.128 | 0.824 |
| **Left hippocampus** | 0.128 | 0.094 | 0.201 | < 0.001 | 0.176 | 0.990 | 0.050 | 0.824 |
| **Left pallidum** | 0.025 | 0.894 | 0.055 | 0.344 | -0.085 | 0.990 | -0.022 | 0.948 |
| **Left putamen** | -0.021 | 0.894 | -0.034 | 0.458 | 0.002 | 0.990 | 0.088 | 0.824 |
| **Left thalamus** | -0.006 | 0.950 | 0.072 | 0.244 | 0.017 | 0.990 | -0.044 | 0.824 |
| **Right accumbens** | -0.088 | 0.421 | -0.067 | 0.269 | -0.029 | 0.990 | 0.043 | 0.824 |
| **Right amygdala** | -0.026 | 0.894 | 0.034 | 0.458 | 0.108 | 0.990 | 0.114 | 0.824 |
| **Right caudate** | -0.029 | 0.894 | -0.100 | 0.083 | 0.067 | 0.990 | 0.106 | 0.824 |
| **Right hippocampus** | 0.006 | 0.950 | 0.129 | 0.017 | 0.089 | 0.990 | 0.079 | 0.824 |
| **Right pallidum** | 0.018 | 0.894 | 0.050 | 0.365 | -0.131 | 0.990 | 0.014 | 0.954 |
| **Right putamen** | -0.040 | 0.894 | -0.042 | 0.407 | 0.065 | 0.990 | 0.076 | 0.824 |
| **Right thalamus** | -0.003 | 0.950 | 0.024 | 0.572 | 0.024 | 0.990 | 0.002 | 0.984 |

**Supplementary Table 11.** Summary of effect size of sex interaction effects in cortical thickness

| | TLE | | | | | | GGE | | | | | |
|---|---|---|---|---|---|---|---|---|---|---|---|---|
| | Sex-diagnosis interaction | | Sex-duration interaction | | Sex-onset interaction | | Sex-diagnosis interaction | | Sex-duration interaction | | Sex-onset interaction | |
| **Regions** | **d** | ***p* (FDR-corrected)** | **d** | ***p* (FDR-corrected)** | **d** | ***p* (FDR-corrected)** | **d** | ***p* (FDR-corrected)** | **d** | ***p* (FDR-corrected)** | **d** | ***p* (FDR-corrected)** |
| **L_bankssts** | -0.004 | 0.938 | -0.018 | 0.927 | 0.012 | 0.961 | 0.030 | 0.763 | -0.057 | 0.980 | -0.078 | 0.705 |
| **L_caudalanteriorcingulate** | 0.002 | 0.938 | -0.034 | 0.810 | -0.033 | 0.903 | 0.024 | 0.808 | 0.108 | 0.980 | -0.100 | 0.595 |
| **L_caudalmiddlefrontal** | -0.018 | 0.938 | -0.046 | 0.639 | 0.005 | 0.961 | 0.009 | 0.933 | 0.049 | 0.980 | -0.092 | 0.656 |
| **L_cuneus** | -0.019 | 0.938 | 0.031 | 0.834 | -0.048 | 0.903 | 0.008 | 0.933 | 0.100 | 0.980 | -0.074 | 0.705 |
| **L_entorhinal** | 0.032 | 0.938 | -0.063 | 0.591 | 0.033 | 0.903 | 0.071 | 0.454 | -0.029 | 0.980 | 0.106 | 0.537 |
| **L_fusiform** | 0.014 | 0.938 | -0.032 | 0.834 | 0.002 | 0.961 | -0.005 | 0.970 | -0.034 | 0.980 | -0.052 | 0.763 |
| **L_inferiorparietal** | -0.012 | 0.938 | -0.005 | 0.942 | -0.028 | 0.903 | 0.055 | 0.505 | 0.049 | 0.980 | -0.057 | 0.744 |
| **L_inferiortemporal** | 0.024 | 0.938 | 0.010 | 0.927 | -0.045 | 0.903 | 0.039 | 0.706 | -0.094 | 0.980 | -0.006 | 0.969 |
| **L_isthmuscingulate** | 0.015 | 0.938 | -0.047 | 0.639 | 0.025 | 0.903 | 0.004 | 0.979 | -0.078 | 0.980 | -0.031 | 0.842 |
| **L_lateraloccipital** | 0.008 | 0.938 | 0.005 | 0.942 | -0.048 | 0.903 | 0.060 | 0.454 | 0.028 | 0.980 | 0.009 | 0.969 |
| **L_lateralorbitofrontal** | 0.004 | 0.938 | -0.035 | 0.796 | 0.049 | 0.903 | 0.060 | 0.454 | 0.076 | 0.980 | -0.063 | 0.734 |
| **L_lingual** | -0.037 | 0.938 | 0.011 | 0.927 | -0.016 | 0.961 | -0.062 | 0.454 | 0.079 | 0.980 | -0.172 | 0.454 |
| **L_medialorbitofrontal** | 0.019 | 0.938 | -0.011 | 0.927 | 0.035 | 0.903 | 0.032 | 0.763 | 0.074 | 0.980 | -0.124 | 0.454 |
| **L_middletemporal** | 0.009 | 0.938 | -0.059 | 0.591 | -0.014 | 0.961 | 0.077 | 0.454 | -0.006 | 0.980 | -0.098 | 0.602 |
| **L_parahippocampal** | -0.002 | 0.938 | 0.048 | 0.639 | -0.024 | 0.903 | 0.074 | 0.454 | -0.068 | 0.980 | 0.027 | 0.858 |
| **L_paracentral** | -0.003 | 0.938 | -0.005 | 0.942 | -0.051 | 0.903 | 0.011 | 0.931 | 0.035 | 0.980 | -0.069 | 0.705 |
| **L_parsopercularis** | -0.019 | 0.938 | -0.035 | 0.796 | 0.009 | 0.961 | 0.065 | 0.454 | -0.022 | 0.980 | -0.060 | 0.744 |
| **L_parsorbitalis** | -0.011 | 0.938 | -0.042 | 0.691 | 0.024 | 0.903 | 0.039 | 0.706 | 0.090 | 0.980 | -0.086 | 0.705 |
| **L_parstriangularis** | 0.007 | 0.938 | -0.083 | 0.561 | 0.038 | 0.903 | 0.098 | 0.270 | 0.021 | 0.980 | -0.077 | 0.705 |
| **L_pericalcarine** | 0.003 | 0.938 | 0.028 | 0.877 | -0.027 | 0.903 | -0.001 | 0.979 | 0.065 | 0.980 | -0.128 | 0.454 |

| | | | | | | | | | | | | |
|---|---|---|---|---|---|---|---|---|---|---|---|---|
| **L_postcentral** | -0.030 | 0.938 | 0.006 | 0.942 | 0.003 | 0.961 | -0.034 | 0.740 | 0.001 | 0.990 | -0.125 | 0.454 |
| **L_posteriorcingulate** | 0.021 | 0.938 | -0.012 | 0.927 | -0.039 | 0.903 | 0.078 | 0.454 | 0.068 | 0.980 | -0.069 | 0.705 |
| **L_precentral** | -0.008 | 0.938 | -0.011 | 0.927 | -0.032 | 0.903 | -0.038 | 0.706 | -0.070 | 0.980 | -0.075 | 0.705 |
| **L_precuneus** | -0.007 | 0.938 | -0.023 | 0.926 | -0.021 | 0.961 | 0.023 | 0.808 | 0.060 | 0.980 | -0.042 | 0.765 |
| **L_rostralanteriorcingulate** | -0.023 | 0.938 | -0.012 | 0.927 | 0.013 | 0.961 | 0.062 | 0.454 | 0.071 | 0.980 | -0.121 | 0.454 |
| **L_rostralmiddlefrontal** | 0.017 | 0.938 | -0.043 | 0.665 | -0.007 | 0.961 | 0.103 | 0.270 | -0.001 | 0.990 | -0.149 | 0.454 |
| **L_superiorfrontal** | -0.005 | 0.938 | -0.044 | 0.665 | 0.012 | 0.961 | 0.023 | 0.808 | 0.109 | 0.980 | -0.127 | 0.454 |
| **L_superiorparietal** | -0.028 | 0.938 | -0.009 | 0.938 | -0.063 | 0.791 | -0.020 | 0.839 | 0.078 | 0.980 | -0.055 | 0.744 |
| **L_superiortemporal** | -0.009 | 0.938 | -0.009 | 0.927 | -0.025 | 0.903 | 0.049 | 0.544 | 0.039 | 0.980 | -0.145 | 0.454 |
| **L_supramarginal** | -0.005 | 0.938 | -0.050 | 0.639 | -0.017 | 0.961 | 0.030 | 0.763 | 0.078 | 0.980 | -0.041 | 0.765 |
| **L_frontalpole** | 0.025 | 0.938 | -0.006 | 0.942 | 0.037 | 0.903 | 0.060 | 0.454 | -0.030 | 0.980 | 0.006 | 0.969 |
| **L_temporalpole** | 0.013 | 0.938 | -0.023 | 0.926 | -0.007 | 0.961 | 0.003 | 0.979 | 0.033 | 0.980 | -0.119 | 0.454 |
| **L_transversetemporal** | -0.014 | 0.938 | -0.013 | 0.927 | 0.004 | 0.961 | -0.070 | 0.454 | -0.013 | 0.980 | -0.172 | 0.454 |
| **L_insula** | 0.007 | 0.938 | -0.023 | 0.926 | 0.037 | 0.903 | 0.062 | 0.454 | 0.036 | 0.980 | -0.128 | 0.454 |
| **R_bankssts** | 0.032 | 0.938 | -0.022 | 0.926 | 0.004 | 0.961 | 0.020 | 0.839 | -0.026 | 0.980 | -0.042 | 0.765 |
| **R_caudalanteriorcingulate** | 0.009 | 0.938 | -0.017 | 0.927 | -0.016 | 0.961 | -0.049 | 0.544 | 0.040 | 0.980 | -0.020 | 0.873 |
| **R_caudalmiddlefrontal** | -0.011 | 0.938 | -0.050 | 0.639 | 0.011 | 0.961 | 0.025 | 0.808 | 0.095 | 0.980 | -0.078 | 0.705 |
| **R_cuneus** | -0.017 | 0.938 | 0.010 | 0.927 | -0.023 | 0.903 | -0.059 | 0.454 | 0.140 | 0.980 | -0.055 | 0.744 |
| **R_entorhinal** | 0.004 | 0.938 | -0.075 | 0.561 | 0.023 | 0.903 | 0.105 | 0.270 | 0.014 | 0.980 | 0.022 | 0.873 |
| **R_fusiform** | -0.020 | 0.938 | 0.004 | 0.942 | -0.040 | 0.903 | 0.053 | 0.505 | 0.102 | 0.980 | -0.068 | 0.705 |
| **R_inferiorparietal** | -0.005 | 0.938 | 0.016 | 0.927 | -0.031 | 0.903 | 0.011 | 0.931 | 0.013 | 0.980 | 0.058 | 0.744 |
| **R_inferiortemporal** | -0.016 | 0.938 | 0.036 | 0.796 | -0.077 | 0.476 | 0.049 | 0.544 | -0.041 | 0.980 | -0.047 | 0.763 |
| **R_isthmuscingulate** | 0.002 | 0.938 | -0.059 | 0.591 | 0.044 | 0.903 | 0.035 | 0.740 | -0.041 | 0.980 | 0.028 | 0.858 |
| **R_lateraloccipital** | 0.008 | 0.938 | 0.064 | 0.591 | -0.082 | 0.476 | -0.002 | 0.979 | -0.006 | 0.980 | 0.051 | 0.763 |
| **R_lateralorbitofrontal** | -0.019 | 0.938 | 0.002 | 0.972 | -0.006 | 0.961 | 0.024 | 0.808 | 0.005 | 0.980 | 0.002 | 0.985 |
| **R_lingual** | -0.028 | 0.938 | 0.012 | 0.927 | -0.030 | 0.903 | -0.069 | 0.454 | 0.102 | 0.980 | -0.112 | 0.488 |
| **R_medialorbitofrontal** | 0.041 | 0.938 | 0.018 | 0.927 | 0.008 | 0.961 | 0.026 | 0.808 | -0.065 | 0.980 | -0.001 | 0.985 |
| **R_middletemporal** | 0.006 | 0.938 | -0.025 | 0.915 | -0.002 | 0.961 | 0.069 | 0.454 | 0.015 | 0.980 | -0.158 | 0.454 |

| | | | | | | | | | | | | |
|---|---|---|---|---|---|---|---|---|---|---|---|---|
| **R_parahippocampal** | 0.033 | 0.938 | 0.000 | 0.999 | -0.039 | 0.903 | 0.115 | 0.270 | -0.029 | 0.980 | -0.026 | 0.858 |
| **R_paracentral** | -0.012 | 0.938 | 0.056 | 0.622 | -0.030 | 0.903 | -0.055 | 0.505 | 0.039 | 0.980 | -0.046 | 0.763 |
| **R_parsopercularis** | -0.005 | 0.938 | -0.040 | 0.712 | 0.008 | 0.961 | 0.015 | 0.876 | 0.025 | 0.980 | -0.046 | 0.763 |
| **R_parsorbitalis** | -0.021 | 0.938 | -0.065 | 0.591 | 0.015 | 0.961 | 0.018 | 0.853 | -0.008 | 0.980 | -0.072 | 0.705 |
| **R_parstriangularis** | 0.022 | 0.938 | 0.007 | 0.942 | -0.017 | 0.961 | 0.012 | 0.931 | -0.011 | 0.980 | -0.039 | 0.775 |
| **R_pericalcarine** | -0.019 | 0.938 | -0.013 | 0.927 | -0.002 | 0.961 | -0.074 | 0.454 | 0.126 | 0.980 | -0.128 | 0.454 |
| **R_postcentral** | -0.052 | 0.938 | 0.012 | 0.927 | -0.011 | 0.961 | -0.034 | 0.740 | -0.014 | 0.980 | -0.081 | 0.705 |
| **R_posteriorcingulate** | 0.023 | 0.938 | -0.002 | 0.972 | 0.009 | 0.961 | 0.068 | 0.454 | 0.076 | 0.980 | -0.067 | 0.705 |
| **R_precentral** | -0.020 | 0.938 | -0.014 | 0.927 | -0.027 | 0.903 | -0.017 | 0.853 | -0.078 | 0.980 | -0.048 | 0.763 |
| **R_precuneus** | -0.037 | 0.938 | 0.022 | 0.926 | -0.024 | 0.903 | 0.053 | 0.505 | 0.026 | 0.980 | 0.070 | 0.705 |
| **R_rostralanteriorcingulate** | 0.007 | 0.938 | 0.010 | 0.927 | -0.004 | 0.961 | 0.008 | 0.933 | 0.142 | 0.980 | -0.157 | 0.454 |
| **R_rostralmiddlefrontal** | 0.004 | 0.938 | -0.029 | 0.857 | -0.007 | 0.961 | 0.039 | 0.706 | 0.068 | 0.980 | -0.183 | 0.454 |
| **R_superiorfrontal** | 0.012 | 0.938 | -0.047 | 0.639 | -0.002 | 0.961 | 0.031 | 0.763 | 0.154 | 0.980 | -0.121 | 0.454 |
| **R_superiorparietal** | -0.024 | 0.938 | 0.026 | 0.915 | -0.063 | 0.791 | 0.001 | 0.979 | -0.010 | 0.980 | 0.024 | 0.873 |
| **R_superiortemporal** | 0.003 | 0.938 | -0.018 | 0.927 | -0.009 | 0.961 | 0.034 | 0.740 | 0.007 | 0.980 | -0.162 | 0.454 |
| **R_supramarginal** | -0.033 | 0.938 | -0.031 | 0.834 | -0.003 | 0.961 | -0.028 | 0.789 | -0.008 | 0.980 | 0.042 | 0.765 |
| **R_frontalpole** | -0.018 | 0.938 | -0.001 | 0.991 | 0.017 | 0.961 | 0.018 | 0.853 | 0.047 | 0.980 | -0.020 | 0.873 |
| **R_temporalpole** | 0.024 | 0.938 | 0.037 | 0.796 | -0.044 | 0.903 | 0.010 | 0.933 | 0.014 | 0.980 | -0.008 | 0.969 |
| **R_transversetemporal** | -0.012 | 0.938 | -0.047 | 0.639 | 0.030 | 0.903 | -0.070 | 0.454 | -0.039 | 0.980 | -0.120 | 0.454 |
| **R_insula** | -0.044 | 0.938 | -0.014 | 0.927 | 0.010 | 0.961 | 0.096 | 0.270 | 0.015 | 0.980 | -0.117 | 0.454 |

**Supplementary Table 12.** Summary of effect size of sex interaction effects in subcortical volume

| | TLE | | | | | | GGE | | | | | |
|---|---|---|---|---|---|---|---|---|---|---|---|---|
| | Sex-diagnosis interaction | | Sex-duration interaction | | Sex-onset interaction | | Sex-diagnosis interaction | | Sex-duration interaction | | Sex-onset interaction | |
| **Regions** | **d** | ***p* (FDR-corrected)** | **d** | ***p* (FDR-corrected)** | **d** | ***p* (FDR-corrected)** | **d** | ***p* (FDR-corrected)** | **d** | ***p* (FDR-corrected)** | **d** | ***p* (FDR-corrected)** |
| **Left accumbens** | -0.046 | 0.404 | -0.051 | 0.853 | 0.003 | 0.968 | -0.050 | 0.449 | 0.075 | 0.829 | -0.047 | 0.984 |
| **Left amygdala** | 0.045 | 0.404 | -0.018 | 0.853 | -0.015 | 0.951 | 0.042 | 0.449 | 0.092 | 0.803 | -0.005 | 0.984 |
| **Left caudate** | -0.016 | 0.880 | -0.029 | 0.853 | 0.022 | 0.951 | 0.046 | 0.449 | 0.099 | 0.803 | -0.003 | 0.984 |
| **Left hippocampus** | 0.022 | 0.830 | -0.027 | 0.853 | -0.026 | 0.951 | 0.056 | 0.449 | 0.028 | 0.902 | 0.067 | 0.984 |
| **Left pallidum** | -0.017 | 0.880 | 0.001 | 0.965 | -0.013 | 0.951 | 0.022 | 0.751 | 0.022 | 0.902 | -0.031 | 0.984 |
| **Left putamen** | 0.007 | 0.904 | -0.014 | 0.853 | 0.007 | 0.951 | 0.041 | 0.449 | 0.046 | 0.849 | -0.031 | 0.984 |
| **Left thalamus** | -0.003 | 0.904 | -0.019 | 0.853 | -0.035 | 0.951 | 0.000 | 0.993 | -0.027 | 0.902 | 0.027 | 0.984 |
| **Right accumbens** | -0.030 | 0.569 | -0.030 | 0.853 | -0.009 | 0.951 | 0.019 | 0.751 | 0.003 | 0.967 | -0.032 | 0.984 |
| **Right amygdala** | 0.033 | 0.569 | 0.017 | 0.853 | -0.026 | 0.951 | 0.057 | 0.449 | 0.093 | 0.803 | 0.006 | 0.984 |
| **Right caudate** | -0.004 | 0.904 | -0.024 | 0.853 | 0.034 | 0.951 | 0.052 | 0.449 | 0.113 | 0.803 | -0.004 | 0.984 |
| **Right hippocampus** | 0.034 | 0.569 | 0.042 | 0.853 | -0.058 | 0.935 | 0.053 | 0.449 | 0.055 | 0.829 | 0.012 | 0.984 |
| **Right pallidum** | 0.007 | 0.904 | -0.007 | 0.958 | -0.013 | 0.951 | 0.059 | 0.449 | 0.070 | 0.829 | -0.058 | 0.984 |
| **Right putamen** | 0.010 | 0.904 | -0.002 | 0.965 | 0.001 | 0.968 | 0.051 | 0.449 | 0.059 | 0.829 | 0.002 | 0.984 |
| **Right thalamus** | -0.005 | 0.904 | -0.013 | 0.853 | -0.010 | 0.951 | -0.006 | 0.943 | 0.014 | 0.921 | 0.013 | 0.984 |

**Supplementary Table 13.** Summary of effect size of sex interaction effects in cortical thickness in left and right TLE

| | Left TLE | | | | | | Right TLE | | | | | |
|---|---|---|---|---|---|---|---|---|---|---|---|---|
| | Sex-diagnosis interaction | | Sex-duration interaction | | Sex-onset interaction | | Sex-diagnosis interaction | | Sex-duration interaction | | Sex-onset interaction | |
| **Regions** | **d** | ***p* (FDR-corrected)** | **d** | ***p* (FDR-corrected)** | **d** | ***p* (FDR-corrected)** | **d** | ***p* (FDR-corrected)** | **d** | ***p* (FDR-corrected)** | **d** | ***p* (FDR-corrected)** |
| **L_bankssts** | 0.003 | 0.954 | -0.004 | 0.981 | -0.037 | 0.986 | -0.014 | 0.900 | -0.034 | 0.872 | 0.077 | 0.988 |
| **L_caudalanteriorcingulate** | 0.032 | 0.944 | -0.056 | 0.981 | -0.011 | 0.986 | -0.039 | 0.900 | -0.001 | 0.989 | -0.057 | 0.988 |
| **L_caudalmiddlefrontal** | -0.010 | 0.945 | -0.051 | 0.981 | 0.011 | 0.986 | -0.028 | 0.900 | -0.035 | 0.872 | -0.003 | 0.988 |
| **L_cuneus** | -0.019 | 0.944 | 0.061 | 0.981 | -0.075 | 0.763 | -0.020 | 0.900 | -0.017 | 0.956 | -0.012 | 0.988 |
| **L_entorhinal** | 0.047 | 0.944 | -0.091 | 0.358 | 0.068 | 0.763 | 0.011 | 0.900 | 0.006 | 0.980 | -0.020 | 0.988 |
| **L_fusiform** | 0.030 | 0.944 | -0.068 | 0.829 | 0.010 | 0.986 | -0.008 | 0.900 | 0.041 | 0.870 | -0.010 | 0.988 |
| **L_inferiorparietal** | -0.003 | 0.954 | 0.001 | 0.992 | -0.036 | 0.986 | -0.026 | 0.900 | -0.022 | 0.937 | -0.013 | 0.988 |
| **L_inferiortemporal** | 0.048 | 0.944 | 0.004 | 0.981 | -0.034 | 0.986 | -0.010 | 0.900 | 0.030 | 0.905 | -0.058 | 0.988 |
| **L_isthmuscingulate** | 0.024 | 0.944 | -0.040 | 0.981 | 0.013 | 0.986 | 0.001 | 0.975 | -0.048 | 0.870 | 0.040 | 0.988 |
| **L_lateraloccipital** | 0.019 | 0.944 | 0.006 | 0.981 | -0.061 | 0.763 | -0.007 | 0.900 | 0.003 | 0.980 | -0.027 | 0.988 |
| **L_lateralorbitofrontal** | 0.008 | 0.952 | -0.034 | 0.981 | 0.060 | 0.763 | -0.001 | 0.975 | -0.025 | 0.937 | 0.033 | 0.988 |
| **L_lingual** | -0.048 | 0.944 | 0.012 | 0.981 | -0.017 | 0.986 | -0.022 | 0.900 | 0.013 | 0.979 | -0.019 | 0.988 |
| **L_medialorbitofrontal** | 0.013 | 0.944 | -0.013 | 0.981 | 0.049 | 0.911 | 0.028 | 0.900 | -0.001 | 0.989 | 0.017 | 0.988 |
| **L_middletemporal** | 0.021 | 0.944 | -0.041 | 0.981 | -0.033 | 0.986 | -0.009 | 0.900 | -0.070 | 0.870 | 0.019 | 0.988 |
| **L_parahippocampal** | 0.000 | 0.998 | 0.014 | 0.981 | 0.009 | 0.986 | -0.004 | 0.923 | 0.125 | 0.655 | -0.076 | 0.988 |
| **L_paracentral** | 0.022 | 0.944 | 0.017 | 0.981 | -0.055 | 0.763 | -0.038 | 0.900 | -0.038 | 0.870 | -0.039 | 0.988 |
| **L_parsopercularis** | -0.005 | 0.954 | -0.017 | 0.981 | 0.019 | 0.986 | -0.038 | 0.900 | -0.062 | 0.870 | -0.003 | 0.988 |
| **L_parsorbitalis** | -0.012 | 0.944 | -0.024 | 0.981 | 0.015 | 0.986 | -0.008 | 0.900 | -0.078 | 0.870 | 0.036 | 0.988 |
| **L_parstriangularis** | 0.019 | 0.944 | -0.068 | 0.829 | 0.037 | 0.986 | -0.010 | 0.900 | -0.104 | 0.655 | 0.041 | 0.988 |
| **L_pericalcarine** | 0.003 | 0.954 | 0.016 | 0.981 | -0.041 | 0.986 | 0.003 | 0.947 | 0.037 | 0.870 | -0.009 | 0.988 |

| | | | | | | | | | | | | |
|---|---|---|---|---|---|---|---|---|---|---|---|---|
| **L_postcentral** | -0.009 | 0.952 | -0.003 | 0.981 | 0.023 | 0.986 | -0.061 | 0.900 | 0.022 | 0.937 | -0.024 | 0.988 |
| **L_posteriorcingulate** | 0.049 | 0.944 | 0.018 | 0.981 | -0.064 | 0.763 | -0.018 | 0.900 | -0.057 | 0.870 | 0.001 | 0.991 |
| **L_precentral** | 0.004 | 0.954 | 0.011 | 0.981 | -0.047 | 0.931 | -0.021 | 0.900 | -0.047 | 0.870 | -0.010 | 0.988 |
| **L_precuneus** | 0.003 | 0.954 | 0.000 | 0.996 | -0.055 | 0.763 | -0.021 | 0.900 | -0.058 | 0.870 | 0.028 | 0.988 |
| **L_rostralanteriorcingulate** | -0.013 | 0.944 | -0.020 | 0.981 | 0.028 | 0.986 | -0.036 | 0.900 | 0.004 | 0.980 | -0.004 | 0.988 |
| **L_rostralmiddlefrontal** | 0.016 | 0.944 | -0.051 | 0.981 | 0.001 | 0.986 | 0.019 | 0.900 | -0.040 | 0.870 | -0.019 | 0.988 |
| **L_superiorfrontal** | 0.003 | 0.954 | -0.044 | 0.981 | 0.017 | 0.986 | -0.014 | 0.900 | -0.040 | 0.870 | 0.005 | 0.988 |
| **L_superiorparietal** | -0.018 | 0.944 | 0.005 | 0.981 | -0.058 | 0.763 | -0.043 | 0.900 | -0.034 | 0.872 | -0.066 | 0.988 |
| **L_superiortemporal** | 0.015 | 0.944 | -0.015 | 0.981 | -0.022 | 0.986 | -0.045 | 0.900 | 0.019 | 0.939 | -0.029 | 0.988 |
| **L_supramarginal** | 0.017 | 0.944 | -0.047 | 0.981 | -0.018 | 0.986 | -0.038 | 0.900 | -0.059 | 0.870 | -0.010 | 0.988 |
| **L_frontalpole** | 0.035 | 0.944 | 0.003 | 0.981 | 0.046 | 0.931 | 0.011 | 0.900 | -0.020 | 0.937 | 0.029 | 0.988 |
| **L_temporalpole** | 0.040 | 0.944 | -0.032 | 0.981 | -0.001 | 0.986 | -0.025 | 0.900 | 0.016 | 0.956 | -0.013 | 0.988 |
| **L_transversetemporal** | -0.018 | 0.944 | 0.007 | 0.981 | -0.016 | 0.986 | -0.008 | 0.900 | -0.053 | 0.870 | 0.033 | 0.988 |
| **L_insula** | 0.022 | 0.944 | -0.010 | 0.981 | 0.006 | 0.986 | -0.015 | 0.900 | -0.044 | 0.870 | 0.085 | 0.988 |
| **R_bankssts** | 0.033 | 0.944 | -0.008 | 0.981 | -0.008 | 0.986 | 0.030 | 0.900 | -0.050 | 0.870 | 0.020 | 0.988 |
| **R_caudalanteriorcingulate** | 0.035 | 0.944 | -0.001 | 0.992 | 0.002 | 0.986 | -0.026 | 0.900 | -0.044 | 0.870 | -0.033 | 0.988 |
| **R_caudalmiddlefrontal** | -0.003 | 0.954 | -0.041 | 0.981 | 0.006 | 0.986 | -0.023 | 0.900 | -0.076 | 0.870 | 0.020 | 0.988 |
| **R_cuneus** | -0.023 | 0.944 | 0.040 | 0.981 | -0.065 | 0.763 | -0.010 | 0.900 | -0.033 | 0.877 | 0.028 | 0.988 |
| **R_entorhinal** | 0.014 | 0.944 | -0.107 | 0.318 | 0.055 | 0.763 | -0.008 | 0.900 | -0.059 | 0.870 | -0.006 | 0.988 |
| **R_fusiform** | -0.018 | 0.944 | -0.018 | 0.981 | -0.020 | 0.986 | -0.023 | 0.900 | 0.023 | 0.937 | -0.064 | 0.988 |
| **R_inferiorparietal** | -0.012 | 0.944 | 0.014 | 0.981 | -0.060 | 0.763 | 0.006 | 0.903 | 0.005 | 0.980 | 0.004 | 0.988 |
| **R_inferiortemporal** | -0.011 | 0.944 | 0.035 | 0.981 | -0.076 | 0.763 | -0.025 | 0.900 | 0.029 | 0.905 | -0.077 | 0.988 |
| **R_isthmuscingulate** | 0.024 | 0.944 | -0.106 | 0.318 | 0.036 | 0.986 | -0.024 | 0.900 | -0.016 | 0.956 | 0.060 | 0.988 |
| **R_lateraloccipital** | 0.009 | 0.952 | 0.097 | 0.318 | -0.104 | 0.759 | 0.009 | 0.900 | 0.009 | 0.980 | -0.052 | 0.988 |
| **R_lateralorbitofrontal** | -0.016 | 0.944 | 0.018 | 0.981 | -0.022 | 0.986 | -0.025 | 0.900 | -0.021 | 0.937 | 0.016 | 0.988 |
| **R_lingual** | -0.039 | 0.944 | -0.005 | 0.981 | -0.032 | 0.986 | -0.012 | 0.900 | 0.044 | 0.870 | -0.032 | 0.988 |
| **R_medialorbitofrontal** | 0.036 | 0.944 | 0.028 | 0.981 | 0.007 | 0.986 | 0.049 | 0.900 | 0.005 | 0.980 | 0.009 | 0.988 |
| **R_middletemporal** | 0.006 | 0.954 | -0.021 | 0.981 | -0.003 | 0.986 | 0.007 | 0.900 | -0.043 | 0.870 | 0.001 | 0.991 |

| | | | | | | | | | | | | |
|---|---|---|---|---|---|---|---|---|---|---|---|---|
| **R_parahippocampal** | 0.032 | 0.944 | -0.034 | 0.981 | 0.004 | 0.986 | 0.036 | 0.900 | 0.042 | 0.870 | -0.094 | 0.988 |
| **R_paracentral** | -0.003 | 0.954 | 0.097 | 0.318 | -0.062 | 0.763 | -0.023 | 0.900 | -0.006 | 0.980 | 0.013 | 0.988 |
| **R_parsopercularis** | 0.005 | 0.954 | -0.032 | 0.981 | 0.018 | 0.986 | -0.019 | 0.900 | -0.052 | 0.870 | -0.004 | 0.988 |
| **R_parsorbitalis** | -0.023 | 0.944 | -0.041 | 0.981 | 0.021 | 0.986 | -0.018 | 0.900 | -0.101 | 0.655 | 0.006 | 0.988 |
| **R_parstriangularis** | 0.019 | 0.944 | 0.027 | 0.981 | 0.001 | 0.986 | 0.026 | 0.900 | -0.029 | 0.905 | -0.044 | 0.988 |
| **R_pericalcarine** | -0.011 | 0.944 | -0.014 | 0.981 | -0.010 | 0.986 | -0.028 | 0.900 | -0.011 | 0.980 | 0.010 | 0.988 |
| **R_postcentral** | -0.037 | 0.944 | 0.052 | 0.981 | -0.013 | 0.986 | -0.072 | 0.900 | -0.051 | 0.870 | -0.003 | 0.988 |
| **R_posteriorcingulate** | 0.052 | 0.944 | -0.011 | 0.981 | -0.016 | 0.986 | -0.014 | 0.900 | -0.004 | 0.980 | 0.044 | 0.988 |
| **R_precentral** | -0.016 | 0.944 | 0.004 | 0.981 | -0.029 | 0.986 | -0.026 | 0.900 | -0.045 | 0.870 | -0.024 | 0.988 |
| **R_precuneus** | -0.024 | 0.944 | 0.049 | 0.981 | -0.071 | 0.763 | -0.053 | 0.900 | -0.025 | 0.937 | 0.034 | 0.988 |
| **R_rostralanteriorcingulate** | 0.022 | 0.944 | 0.014 | 0.981 | 0.004 | 0.986 | -0.014 | 0.900 | 0.005 | 0.980 | -0.011 | 0.988 |
| **R_rostralmiddlefrontal** | 0.010 | 0.945 | -0.005 | 0.981 | -0.025 | 0.986 | -0.006 | 0.903 | -0.076 | 0.870 | 0.021 | 0.988 |
| **R_superiorfrontal** | 0.013 | 0.944 | -0.035 | 0.981 | 0.010 | 0.986 | 0.012 | 0.900 | -0.076 | 0.870 | -0.018 | 0.988 |
| **R_superiorparietal** | -0.026 | 0.944 | 0.027 | 0.981 | -0.073 | 0.763 | -0.020 | 0.900 | 0.010 | 0.980 | -0.049 | 0.988 |
| **R_superiortemporal** | 0.000 | 0.998 | -0.028 | 0.981 | 0.010 | 0.986 | 0.007 | 0.900 | -0.014 | 0.971 | -0.033 | 0.988 |
| **R_supramarginal** | -0.030 | 0.944 | -0.007 | 0.981 | -0.011 | 0.986 | -0.037 | 0.900 | -0.066 | 0.870 | 0.007 | 0.988 |
| **R_frontalpole** | -0.024 | 0.944 | 0.070 | 0.829 | -0.018 | 0.986 | -0.009 | 0.900 | -0.102 | 0.655 | 0.066 | 0.988 |
| **R_temporalpole** | 0.022 | 0.944 | 0.008 | 0.981 | -0.004 | 0.986 | 0.028 | 0.900 | 0.066 | 0.870 | -0.085 | 0.988 |
| **R_transversetemporal** | -0.012 | 0.944 | -0.049 | 0.981 | 0.022 | 0.986 | -0.012 | 0.900 | -0.038 | 0.870 | 0.038 | 0.988 |
| **R_insula** | -0.044 | 0.944 | 0.009 | 0.981 | 0.004 | 0.986 | -0.047 | 0.900 | -0.045 | 0.870 | 0.018 | 0.988 |

**Supplementary Table 14.** Summary of effect size of sex interaction effects in subcortical volume in left and right TLE

| | Left TLE | | | | | | Right TLE | | | | | |
|---|---|---|---|---|---|---|---|---|---|---|---|---|
| | Sex-diagnosis interaction | | Sex-duration interaction | | Sex-onset interaction | | Sex-diagnosis interaction | | Sex-duration interaction | | Sex-onset interaction | |
| **Regions** | **d** | ***p* (FDR-corrected)** | **d** | ***p* (FDR-corrected)** | **d** | ***p* (FDR-corrected)** | **d** | ***p* (FDR-corrected)** | **d** | ***p* (FDR-corrected)** | **d** | ***p* (FDR-corrected)** |
| **Left accumbens** | -0.042 | 0.582 | -0.034 | 0.960 | -0.008 | 0.915 | -0.049 | 0.373 | -0.071 | 0.746 | 0.020 | 0.975 |
| **Left amygdala** | 0.052 | 0.582 | 0.017 | 0.960 | -0.027 | 0.843 | 0.041 | 0.483 | -0.052 | 0.746 | 0.003 | 0.975 |
| **Left caudate** | -0.025 | 0.882 | -0.034 | 0.960 | 0.028 | 0.843 | -0.003 | 0.967 | -0.018 | 0.813 | 0.014 | 0.975 |
| **Left hippocampus** | 0.013 | 0.892 | 0.028 | 0.960 | -0.065 | 0.524 | 0.048 | 0.373 | -0.058 | 0.746 | 0.033 | 0.975 |
| **Left pallidum** | -0.019 | 0.882 | 0.024 | 0.960 | -0.025 | 0.843 | -0.013 | 0.908 | -0.015 | 0.813 | 0.003 | 0.975 |
| **Left putamen** | 0.006 | 0.892 | -0.012 | 0.960 | 0.004 | 0.924 | 0.008 | 0.919 | -0.005 | 0.914 | 0.010 | 0.975 |
| **Left thalamus** | -0.006 | 0.892 | 0.010 | 0.960 | -0.045 | 0.624 | 0.001 | 0.967 | -0.040 | 0.813 | -0.026 | 0.975 |
| **Right accumbens** | -0.045 | 0.582 | 0.007 | 0.960 | -0.048 | 0.624 | -0.011 | 0.908 | -0.076 | 0.746 | 0.039 | 0.975 |
| **Right amygdala** | 0.012 | 0.892 | 0.057 | 0.960 | -0.082 | 0.318 | 0.064 | 0.373 | -0.049 | 0.746 | 0.035 | 0.975 |
| **Right caudate** | -0.018 | 0.882 | -0.027 | 0.960 | 0.046 | 0.624 | 0.016 | 0.908 | -0.022 | 0.813 | 0.020 | 0.975 |
| **Right hippocampus** | 0.029 | 0.882 | 0.042 | 0.960 | -0.088 | 0.318 | 0.053 | 0.373 | -0.017 | 0.813 | -0.034 | 0.975 |
| **Right pallidum** | -0.008 | 0.892 | 0.008 | 0.960 | -0.019 | 0.843 | 0.027 | 0.862 | -0.026 | 0.813 | -0.004 | 0.975 |
| **Right putamen** | 0.004 | 0.892 | 0.002 | 0.960 | -0.012 | 0.889 | 0.019 | 0.908 | -0.018 | 0.813 | 0.020 | 0.975 |
| **Right thalamus** | -0.022 | 0.882 | 0.002 | 0.960 | -0.018 | 0.843 | 0.018 | 0.908 | -0.051 | 0.746 | 0.002 | 0.975 |

**Supplementary Table 15.** Summary of effect size of sex interaction effects in cortical thickness in MTS and non-lesional TLE patients

| | MTS | | | | | | Non-lesional TLE | | | | | |
|---|---|---|---|---|---|---|---|---|---|---|---|---|
| | Sex-diagnosis interaction | | Sex-duration interaction | | Sex-onset interaction | | Sex-diagnosis interaction | | Sex-duration interaction | | Sex-onset interaction | |
| **Regions** | **d** | ***p* (FDR-corrected)** | **d** | ***p* (FDR-corrected)** | **d** | ***p* (FDR-corrected)** | **d** | ***p* (FDR-corrected)** | **d** | ***p* (FDR-corrected)** | **d** | ***p* (FDR-corrected)** |
| **L_bankssts** | 0.016 | 0.971 | -0.029 | 0.916 | 0.039 | 0.960 | -0.030 | 0.955 | -0.043 | 0.855 | 0.013 | 0.986 |
| **L_caudalanteriorcingulate** | -0.003 | 0.971 | -0.063 | 0.901 | -0.032 | 0.960 | 0.008 | 0.999 | 0.023 | 0.888 | -0.044 | 0.986 |
| **L_caudalmiddlefrontal** | -0.002 | 0.971 | -0.041 | 0.916 | 0.038 | 0.960 | -0.037 | 0.803 | -0.086 | 0.837 | -0.017 | 0.986 |
| **L_cuneus** | 0.003 | 0.971 | -0.019 | 0.916 | -0.023 | 0.960 | -0.049 | 0.739 | 0.079 | 0.837 | -0.049 | 0.986 |
| **L_entorhinal** | 0.055 | 0.858 | -0.139 | 0.059 | 0.090 | 0.960 | 0.003 | 0.999 | 0.001 | 0.992 | 0.005 | 0.986 |
| **L_fusiform** | 0.026 | 0.971 | -0.096 | 0.495 | 0.042 | 0.960 | 0.000 | 0.999 | 0.039 | 0.855 | -0.030 | 0.986 |
| **L_inferiorparietal** | 0.020 | 0.971 | -0.018 | 0.916 | -0.009 | 0.970 | -0.053 | 0.736 | -0.040 | 0.855 | -0.007 | 0.986 |
| **L_inferiortemporal** | 0.047 | 0.971 | -0.015 | 0.916 | -0.002 | 0.982 | -0.004 | 0.999 | 0.019 | 0.906 | -0.085 | 0.986 |
| **L_isthmuscingulate** | 0.019 | 0.971 | -0.057 | 0.901 | 0.030 | 0.960 | 0.010 | 0.999 | -0.028 | 0.882 | 0.016 | 0.986 |
| **L_lateraloccipital** | 0.035 | 0.971 | -0.005 | 0.942 | -0.022 | 0.960 | -0.024 | 0.999 | -0.028 | 0.882 | -0.041 | 0.986 |
| **L_lateralorbitofrontal** | -0.005 | 0.971 | -0.023 | 0.916 | 0.024 | 0.960 | 0.014 | 0.999 | -0.033 | 0.882 | 0.065 | 0.986 |
| **L_lingual** | -0.025 | 0.971 | 0.003 | 0.955 | 0.005 | 0.982 | -0.054 | 0.736 | 0.008 | 0.955 | -0.030 | 0.986 |
| **L_medialorbitofrontal** | -0.005 | 0.971 | -0.047 | 0.916 | 0.077 | 0.960 | 0.052 | 0.736 | 0.114 | 0.780 | -0.053 | 0.986 |
| **L_middletemporal** | 0.028 | 0.971 | -0.100 | 0.495 | 0.027 | 0.960 | -0.014 | 0.999 | -0.030 | 0.882 | -0.047 | 0.986 |
| **L_parahippocampal** | 0.033 | 0.971 | 0.019 | 0.916 | -0.017 | 0.970 | -0.047 | 0.739 | 0.026 | 0.882 | 0.022 | 0.986 |
| **L_paracentral** | 0.011 | 0.971 | 0.021 | 0.916 | -0.078 | 0.960 | -0.023 | 0.999 | -0.077 | 0.837 | 0.003 | 0.986 |
| **L_parsopercularis** | -0.015 | 0.971 | -0.027 | 0.916 | 0.001 | 0.982 | -0.024 | 0.999 | -0.050 | 0.855 | 0.021 | 0.986 |
| **L_parsorbitalis** | -0.022 | 0.971 | -0.030 | 0.916 | -0.021 | 0.960 | 0.004 | 0.999 | -0.037 | 0.855 | 0.063 | 0.986 |
| **L_parstriangularis** | 0.012 | 0.971 | -0.076 | 0.792 | 0.049 | 0.960 | 0.000 | 0.999 | -0.109 | 0.780 | 0.035 | 0.986 |
| **L_pericalcarine** | -0.005 | 0.971 | 0.040 | 0.916 | -0.056 | 0.960 | 0.014 | 0.999 | 0.035 | 0.869 | -0.006 | 0.986 |

| | | | | | | | | | | | | |
|---|---|---|---|---|---|---|---|---|---|---|---|---|
| **L_postcentral** | -0.013 | 0.971 | 0.048 | 0.916 | -0.035 | 0.960 | -0.053 | 0.736 | -0.088 | 0.837 | 0.075 | 0.986 |
| **L_posteriorcingulate** | 0.030 | 0.971 | -0.067 | 0.901 | -0.012 | 0.970 | 0.011 | 0.999 | 0.069 | 0.837 | -0.065 | 0.986 |
| **L_precentral** | 0.002 | 0.971 | 0.007 | 0.933 | -0.020 | 0.960 | -0.019 | 0.999 | -0.057 | 0.855 | -0.033 | 0.986 |
| **L_precuneus** | 0.017 | 0.971 | -0.035 | 0.916 | -0.025 | 0.960 | -0.037 | 0.803 | -0.046 | 0.855 | 0.022 | 0.986 |
| **L_rostralanteriorcingulate** | -0.041 | 0.971 | -0.018 | 0.916 | 0.028 | 0.960 | 0.000 | 0.999 | 0.040 | 0.855 | -0.029 | 0.986 |
| **L_rostralmiddlefrontal** | 0.014 | 0.971 | -0.020 | 0.916 | -0.015 | 0.970 | 0.021 | 0.999 | -0.075 | 0.837 | -0.005 | 0.986 |
| **L_superiorfrontal** | -0.012 | 0.971 | -0.030 | 0.916 | 0.001 | 0.982 | 0.005 | 0.999 | -0.047 | 0.855 | 0.013 | 0.986 |
| **L_superiorparietal** | -0.008 | 0.971 | -0.009 | 0.933 | -0.066 | 0.960 | -0.055 | 0.736 | -0.045 | 0.855 | -0.027 | 0.986 |
| **L_superiortemporal** | -0.014 | 0.971 | -0.039 | 0.916 | 0.005 | 0.982 | -0.003 | 0.999 | 0.054 | 0.855 | -0.081 | 0.986 |
| **L_supramarginal** | 0.002 | 0.971 | -0.058 | 0.901 | -0.029 | 0.960 | -0.014 | 0.999 | -0.049 | 0.855 | 0.006 | 0.986 |
| **L_frontalpole** | -0.002 | 0.971 | 0.036 | 0.916 | 0.034 | 0.960 | 0.060 | 0.736 | -0.011 | 0.932 | -0.008 | 0.986 |
| **L_temporalpole** | 0.018 | 0.971 | -0.030 | 0.916 | -0.032 | 0.960 | 0.007 | 0.999 | -0.025 | 0.882 | 0.031 | 0.986 |
| **L_transversetemporal** | -0.012 | 0.971 | 0.023 | 0.916 | 0.005 | 0.982 | -0.017 | 0.999 | -0.070 | 0.837 | 0.001 | 0.995 |
| **L_insula** | 0.010 | 0.971 | -0.032 | 0.916 | 0.036 | 0.960 | 0.002 | 0.999 | -0.007 | 0.959 | 0.042 | 0.986 |
| **R_bankssts** | 0.026 | 0.971 | -0.002 | 0.962 | 0.002 | 0.982 | 0.040 | 0.803 | -0.045 | 0.855 | 0.002 | 0.995 |
| **R_caudalanteriorcingulate** | 0.008 | 0.971 | -0.005 | 0.942 | -0.045 | 0.960 | 0.010 | 0.999 | -0.027 | 0.882 | 0.019 | 0.986 |
| **R_caudalmiddlefrontal** | 0.003 | 0.971 | -0.058 | 0.901 | 0.038 | 0.960 | -0.029 | 0.955 | -0.065 | 0.837 | -0.006 | 0.986 |
| **R_cuneus** | -0.027 | 0.971 | 0.013 | 0.925 | -0.025 | 0.960 | -0.006 | 0.999 | 0.029 | 0.882 | -0.030 | 0.986 |
| **R_entorhinal** | -0.008 | 0.971 | -0.089 | 0.557 | 0.056 | 0.960 | 0.019 | 0.999 | -0.042 | 0.855 | -0.031 | 0.986 |
| **R_fusiform** | -0.012 | 0.971 | 0.006 | 0.933 | -0.035 | 0.960 | -0.032 | 0.955 | -0.011 | 0.932 | -0.040 | 0.986 |
| **R_inferiorparietal** | 0.007 | 0.971 | 0.027 | 0.916 | -0.014 | 0.970 | -0.019 | 0.999 | -0.016 | 0.932 | -0.040 | 0.986 |
| **R_inferiortemporal** | 0.010 | 0.971 | 0.018 | 0.916 | -0.048 | 0.960 | -0.050 | 0.736 | 0.022 | 0.888 | -0.088 | 0.986 |
| **R_isthmuscingulate** | -0.029 | 0.971 | -0.016 | 0.916 | 0.049 | 0.960 | 0.041 | 0.803 | -0.067 | 0.837 | 0.000 | 0.999 |
| **R_lateraloccipital** | 0.031 | 0.971 | 0.054 | 0.901 | -0.052 | 0.960 | -0.020 | 0.999 | 0.038 | 0.855 | -0.078 | 0.986 |
| **R_lateralorbitofrontal** | -0.016 | 0.971 | 0.016 | 0.916 | -0.056 | 0.960 | -0.025 | 0.999 | -0.021 | 0.888 | 0.063 | 0.986 |
| **R_lingual** | -0.026 | 0.971 | 0.020 | 0.916 | -0.032 | 0.960 | -0.031 | 0.955 | 0.008 | 0.955 | -0.027 | 0.986 |
| **R_medialorbitofrontal** | 0.060 | 0.858 | 0.012 | 0.925 | 0.020 | 0.960 | 0.018 | 0.999 | -0.004 | 0.983 | 0.030 | 0.986 |
| **R_middletemporal** | 0.004 | 0.971 | -0.028 | 0.916 | 0.017 | 0.970 | 0.009 | 0.999 | -0.012 | 0.932 | -0.028 | 0.986 |

| | | | | | | | | | | | | |
|---|---|---|---|---|---|---|---|---|---|---|---|---|
| **R_parahippocampal** | 0.056 | 0.858 | 0.008 | 0.933 | -0.050 | 0.960 | 0.005 | 0.999 | -0.055 | 0.855 | 0.013 | 0.986 |
| **R_paracentral** | -0.007 | 0.971 | 0.082 | 0.691 | -0.027 | 0.960 | -0.019 | 0.999 | 0.013 | 0.932 | -0.028 | 0.986 |
| **R_parsopercularis** | -0.005 | 0.971 | -0.057 | 0.901 | 0.023 | 0.960 | -0.003 | 0.999 | -0.002 | 0.991 | -0.025 | 0.986 |
| **R_parsorbitalis** | -0.028 | 0.971 | -0.060 | 0.901 | -0.002 | 0.982 | -0.012 | 0.999 | -0.067 | 0.837 | 0.028 | 0.986 |
| **R_parstriangularis** | 0.030 | 0.971 | -0.012 | 0.925 | -0.009 | 0.970 | 0.012 | 0.999 | 0.030 | 0.882 | -0.024 | 0.986 |
| **R_pericalcarine** | -0.034 | 0.971 | 0.007 | 0.933 | -0.015 | 0.970 | 0.000 | 0.999 | -0.014 | 0.932 | -0.015 | 0.986 |
| **R_postcentral** | -0.057 | 0.858 | 0.055 | 0.901 | -0.048 | 0.960 | -0.044 | 0.752 | -0.040 | 0.855 | 0.032 | 0.986 |
| **R_posteriorcingulate** | 0.010 | 0.971 | 0.014 | 0.920 | -0.001 | 0.982 | 0.039 | 0.803 | -0.003 | 0.991 | 0.013 | 0.986 |
| **R_precentral** | -0.016 | 0.971 | -0.010 | 0.933 | 0.013 | 0.970 | -0.025 | 0.999 | -0.021 | 0.888 | -0.074 | 0.986 |
| **R_precuneus** | -0.021 | 0.971 | 0.023 | 0.916 | -0.032 | 0.960 | -0.060 | 0.736 | 0.000 | 0.992 | 0.004 | 0.986 |
| **R_rostralanteriorcingulate** | 0.001 | 0.975 | 0.039 | 0.916 | -0.033 | 0.960 | 0.015 | 0.999 | -0.013 | 0.932 | 0.029 | 0.986 |
| **R_rostralmiddlefrontal** | 0.008 | 0.971 | -0.025 | 0.916 | -0.011 | 0.970 | -0.002 | 0.999 | -0.037 | 0.855 | -0.004 | 0.986 |
| **R_superiorfrontal** | 0.018 | 0.971 | -0.033 | 0.916 | -0.012 | 0.970 | 0.005 | 0.999 | -0.076 | 0.837 | 0.013 | 0.986 |
| **R_superiorparietal** | -0.013 | 0.971 | 0.024 | 0.916 | -0.059 | 0.960 | -0.038 | 0.803 | 0.015 | 0.932 | -0.048 | 0.986 |
| **R_superiortemporal** | 0.008 | 0.971 | -0.019 | 0.916 | 0.012 | 0.970 | -0.003 | 0.999 | -0.021 | 0.888 | -0.029 | 0.986 |
| **R_supramarginal** | -0.036 | 0.971 | -0.024 | 0.916 | -0.004 | 0.982 | -0.030 | 0.955 | -0.027 | 0.882 | -0.016 | 0.986 |
| **R_frontalpole** | -0.033 | 0.971 | -0.030 | 0.916 | 0.024 | 0.960 | 0.001 | 0.999 | 0.085 | 0.837 | -0.021 | 0.986 |
| **R_temporalpole** | 0.024 | 0.971 | 0.032 | 0.916 | -0.010 | 0.970 | 0.025 | 0.999 | 0.056 | 0.855 | -0.092 | 0.986 |
| **R_transversetemporal** | -0.025 | 0.971 | -0.030 | 0.916 | 0.049 | 0.960 | 0.005 | 0.999 | -0.052 | 0.855 | -0.007 | 0.986 |
| **R_insula** | -0.045 | 0.971 | 0.007 | 0.933 | 0.022 | 0.960 | -0.045 | 0.752 | -0.043 | 0.855 | -0.007 | 0.986 |

**Supplementary Table 16.** Summary of effect size of sex interaction effects in subcortical volume in MTS and non-lesional TLE patients

| | MTS | | | | | | Non-lesional TLE | | | | | |
|---|---|---|---|---|---|---|---|---|---|---|---|---|
| | Sex-diagnosis interaction | | Sex-duration interaction | | Sex-onset interaction | | Sex-diagnosis interaction | | Sex-duration interaction | | Sex-onset interaction | |
| **Regions** | **d** | ***p* (FDR-corrected)** | **d** | ***p* (FDR-corrected)** | **d** | ***p* (FDR-corrected)** | **d** | ***p* (FDR-corrected)** | **d** | ***p* (FDR-corrected)** | **d** | ***p* (FDR-corrected)** |
| **Left accumbens** | -0.050 | 0.402 | -0.058 | 0.775 | 0.022 | 0.828 | -0.042 | 0.872 | -0.031 | 0.713 | -0.036 | 0.972 |
| **Left amygdala** | 0.073 | 0.122 | -0.077 | 0.449 | 0.049 | 0.774 | 0.009 | 0.872 | 0.021 | 0.713 | -0.065 | 0.972 |
| **Left caudate** | 0.002 | 0.948 | -0.048 | 0.882 | 0.051 | 0.774 | -0.042 | 0.872 | -0.036 | 0.713 | 0.002 | 0.972 |
| **Left hippocampus** | 0.024 | 0.805 | -0.022 | 0.995 | -0.044 | 0.774 | 0.022 | 0.872 | -0.061 | 0.713 | 0.020 | 0.972 |
| **Left pallidum** | -0.012 | 0.815 | 0.002 | 0.995 | -0.016 | 0.828 | -0.027 | 0.872 | -0.010 | 0.837 | -0.008 | 0.972 |
| **Left putamen** | 0.018 | 0.805 | -0.017 | 0.995 | 0.026 | 0.828 | -0.010 | 0.872 | -0.033 | 0.713 | -0.003 | 0.972 |
| **Left thalamus** | 0.018 | 0.805 | -0.016 | 0.995 | -0.026 | 0.828 | -0.033 | 0.872 | -0.073 | 0.713 | -0.023 | 0.972 |
| **Right accumbens** | -0.042 | 0.471 | -0.003 | 0.995 | -0.027 | 0.828 | -0.017 | 0.872 | -0.050 | 0.713 | -0.010 | 0.972 |
| **Right amygdala** | 0.048 | 0.402 | 0.022 | 0.995 | -0.004 | 0.969 | 0.014 | 0.872 | -0.022 | 0.713 | -0.032 | 0.972 |
| **Right caudate** | 0.011 | 0.815 | -0.037 | 0.995 | 0.068 | 0.725 | -0.025 | 0.872 | -0.033 | 0.713 | 0.007 | 0.972 |
| **Right hippocampus** | 0.038 | 0.485 | 0.082 | 0.449 | -0.073 | 0.725 | 0.034 | 0.872 | -0.059 | 0.713 | -0.002 | 0.972 |
| **Right pallidum** | 0.011 | 0.815 | -0.003 | 0.995 | 0.002 | 0.969 | -0.001 | 0.984 | -0.026 | 0.713 | -0.034 | 0.972 |
| **Right putamen** | 0.020 | 0.805 | 0.000 | 0.995 | 0.040 | 0.774 | -0.005 | 0.939 | -0.023 | 0.713 | -0.039 | 0.972 |
| **Right thalamus** | 0.002 | 0.948 | 0.013 | 0.995 | -0.017 | 0.828 | -0.016 | 0.872 | -0.076 | 0.713 | 0.011 | 0.972 |

**Supplementary Table 17.** Effect size of age in cortical thickness in male and female patients with TLE and GGE

| | TLE | | | | | | GGE | | | | | |
|---|---|---|---|---|---|---|---|---|---|---|---|---|
| | Male | | Female | | Age-sex interaction | | Male | | Female | | Age-sex interaction | |
| **Regions** | **d** | ***p* (FDR-corrected)** | **d** | ***p* (FDR-corrected)** | **d** | ***p* (FDR-corrected)** | **d** | ***p* (FDR-corrected)** | **d** | ***p* (FDR-corrected)** | **d** | ***p* (FDR-corrected)** |
| **L_bankssts** | -0.282 | < 0.001 | -0.232 | < 0.001 | -0.012 | 0.920 | -0.537 | 0.002 | -0.299 | 0.008 | -0.065 | 0.894 |
| **L_caudalanteriorcingulate** | -0.206 | < 0.001 | -0.082 | 0.057 | -0.063 | 0.705 | 0.008 | 0.991 | -0.073 | 0.458 | 0.042 | 0.894 |
| **L_caudalmiddlefrontal** | -0.508 | < 0.001 | -0.425 | < 0.001 | -0.027 | 0.796 | -0.387 | 0.017 | -0.303 | 0.008 | 0.010 | 0.918 |
| **L_cuneus** | -0.189 | < 0.001 | -0.195 | < 0.001 | 0.001 | 0.980 | -0.123 | 0.486 | -0.276 | 0.014 | 0.089 | 0.894 |
| **L_entorhinal** | 0.007 | 0.883 | 0.064 | 0.135 | -0.029 | 0.796 | -0.127 | 0.476 | -0.147 | 0.172 | 0.040 | 0.894 |
| **L_fusiform** | -0.230 | < 0.001 | -0.158 | < 0.001 | -0.030 | 0.796 | -0.390 | 0.017 | -0.181 | 0.093 | -0.060 | 0.894 |
| **L_inferiorparietal** | -0.413 | < 0.001 | -0.316 | < 0.001 | -0.029 | 0.796 | -0.155 | 0.356 | -0.190 | 0.079 | 0.040 | 0.894 |
| **L_inferiortemporal** | -0.250 | < 0.001 | -0.167 | < 0.001 | -0.041 | 0.705 | -0.366 | 0.024 | -0.084 | 0.408 | -0.094 | 0.894 |
| **L_isthmuscingulate** | -0.372 | < 0.001 | -0.293 | < 0.001 | -0.024 | 0.796 | -0.661 | < 0.001 | -0.252 | 0.024 | -0.106 | 0.894 |
| **L_lateraloccipital** | -0.231 | < 0.001 | -0.145 | < 0.001 | -0.042 | 0.705 | -0.059 | 0.759 | -0.212 | 0.051 | 0.082 | 0.894 |
| **L_lateralorbitofrontal** | -0.278 | < 0.001 | -0.282 | < 0.001 | 0.014 | 0.907 | -0.121 | 0.489 | -0.215 | 0.050 | 0.069 | 0.894 |
| **L_lingual** | -0.291 | < 0.001 | -0.305 | < 0.001 | 0.006 | 0.950 | -0.211 | 0.226 | -0.204 | 0.059 | 0.010 | 0.918 |
| **L_medialorbitofrontal** | -0.183 | < 0.001 | -0.220 | < 0.001 | 0.016 | 0.899 | -0.071 | 0.711 | -0.030 | 0.749 | -0.010 | 0.918 |
| **L_middletemporal** | -0.475 | < 0.001 | -0.281 | < 0.001 | -0.084 | 0.530 | -0.535 | 0.002 | -0.277 | 0.014 | -0.062 | 0.894 |
| **L_parahippocampal** | -0.081 | 0.091 | -0.122 | 0.005 | 0.025 | 0.796 | 0.028 | 0.883 | 0.099 | 0.334 | -0.042 | 0.894 |
| **L_paracentral** | -0.296 | < 0.001 | -0.179 | < 0.001 | -0.054 | 0.705 | -0.194 | 0.242 | -0.234 | 0.035 | 0.041 | 0.894 |
| **L_parsopercularis** | -0.519 | < 0.001 | -0.500 | < 0.001 | -0.010 | 0.925 | -0.460 | 0.004 | -0.207 | 0.057 | -0.053 | 0.894 |
| **L_parsorbitalis** | -0.287 | < 0.001 | -0.231 | < 0.001 | -0.017 | 0.899 | -0.004 | 0.991 | -0.126 | 0.229 | 0.061 | 0.894 |
| **L_parstriangularis** | -0.473 | < 0.001 | -0.334 | < 0.001 | -0.046 | 0.705 | -0.450 | 0.005 | -0.299 | 0.008 | -0.011 | 0.918 |
| **L_pericalcarine** | -0.172 | < 0.001 | -0.194 | < 0.001 | 0.014 | 0.907 | -0.101 | 0.578 | -0.130 | 0.227 | 0.028 | 0.918 |
| **L_postcentral** | -0.246 | < 0.001 | -0.268 | < 0.001 | 0.021 | 0.832 | -0.328 | 0.043 | -0.177 | 0.099 | -0.043 | 0.894 |

| | | | | | | | | | | | | |
|---|---|---|---|---|---|---|---|---|---|---|---|---|
| **L_posteriorcingulate** | -0.476 | < 0.001 | -0.292 | < 0.001 | -0.058 | 0.705 | -0.178 | 0.284 | -0.183 | 0.090 | 0.018 | 0.918 |
| **L_precentral** | -0.423 | < 0.001 | -0.334 | < 0.001 | -0.042 | 0.705 | -0.501 | 0.002 | -0.314 | 0.007 | -0.084 | 0.894 |
| **L_precuneus** | -0.401 | < 0.001 | -0.306 | < 0.001 | -0.041 | 0.705 | -0.170 | 0.301 | -0.262 | 0.021 | 0.074 | 0.894 |
| **L_rostralanteriorcingulate** | -0.158 | < 0.001 | -0.189 | < 0.001 | 0.011 | 0.920 | -0.226 | 0.193 | -0.127 | 0.227 | -0.022 | 0.918 |
| **L_rostralmiddlefrontal** | -0.444 | < 0.001 | -0.342 | < 0.001 | -0.044 | 0.705 | -0.474 | 0.004 | -0.138 | 0.200 | -0.085 | 0.894 |
| **L_superiorfrontal** | -0.491 | < 0.001 | -0.440 | < 0.001 | -0.021 | 0.832 | -0.458 | 0.004 | -0.449 | < 0.001 | 0.058 | 0.894 |
| **L_superiorparietal** | -0.358 | < 0.001 | -0.216 | < 0.001 | -0.062 | 0.705 | -0.061 | 0.756 | -0.214 | 0.050 | 0.090 | 0.894 |
| **L_superiortemporal** | -0.401 | < 0.001 | -0.333 | < 0.001 | -0.028 | 0.796 | -0.591 | < 0.001 | -0.344 | 0.003 | -0.012 | 0.918 |
| **L_supramarginal** | -0.477 | < 0.001 | -0.317 | < 0.001 | -0.064 | 0.705 | -0.273 | 0.100 | -0.313 | 0.007 | 0.071 | 0.894 |
| **L_frontalpole** | -0.110 | 0.021 | -0.165 | < 0.001 | 0.021 | 0.832 | 0.072 | 0.711 | 0.106 | 0.306 | -0.023 | 0.918 |
| **L_temporalpole** | -0.034 | 0.472 | 0.028 | 0.507 | -0.031 | 0.796 | -0.248 | 0.146 | -0.057 | 0.559 | -0.057 | 0.894 |
| **L_transversetemporal** | -0.345 | < 0.001 | -0.329 | < 0.001 | -0.006 | 0.950 | -0.512 | 0.002 | -0.217 | 0.050 | -0.087 | 0.894 |
| **L_insula** | -0.322 | < 0.001 | -0.343 | < 0.001 | 0.016 | 0.899 | -0.503 | 0.002 | -0.278 | 0.014 | -0.035 | 0.918 |
| **R_bankssts** | -0.329 | < 0.001 | -0.282 | < 0.001 | -0.017 | 0.899 | -0.484 | 0.003 | -0.430 | < 0.001 | 0.011 | 0.918 |
| **R_caudalanteriorcingulate** | -0.220 | < 0.001 | -0.142 | < 0.001 | -0.033 | 0.796 | -0.091 | 0.623 | -0.113 | 0.275 | 0.018 | 0.918 |
| **R_caudalmiddlefrontal** | -0.443 | < 0.001 | -0.300 | < 0.001 | -0.050 | 0.705 | -0.192 | 0.242 | -0.231 | 0.036 | 0.063 | 0.894 |
| **R_cuneus** | -0.197 | < 0.001 | -0.221 | < 0.001 | 0.007 | 0.950 | 0.046 | 0.800 | -0.254 | 0.024 | 0.155 | 0.894 |
| **R_entorhinal** | -0.042 | 0.377 | 0.078 | 0.070 | -0.060 | 0.705 | -0.034 | 0.862 | -0.121 | 0.246 | 0.057 | 0.894 |
| **R_fusiform** | -0.296 | < 0.001 | -0.206 | < 0.001 | -0.039 | 0.745 | -0.006 | 0.991 | -0.172 | 0.109 | 0.090 | 0.894 |
| **R_inferiorparietal** | -0.417 | < 0.001 | -0.345 | < 0.001 | -0.005 | 0.950 | -0.214 | 0.225 | -0.351 | 0.003 | 0.091 | 0.894 |
| **R_inferiortemporal** | -0.281 | < 0.001 | -0.169 | < 0.001 | -0.041 | 0.705 | -0.238 | 0.165 | -0.081 | 0.423 | -0.051 | 0.894 |
| **R_isthmuscingulate** | -0.340 | < 0.001 | -0.290 | < 0.001 | -0.024 | 0.796 | -0.553 | 0.002 | -0.354 | 0.003 | -0.024 | 0.918 |
| **R_lateraloccipital** | -0.234 | < 0.001 | -0.223 | < 0.001 | 0.001 | 0.980 | -0.054 | 0.774 | -0.148 | 0.172 | 0.054 | 0.894 |
| **R_lateralorbitofrontal** | -0.239 | < 0.001 | -0.228 | < 0.001 | 0.004 | 0.950 | -0.193 | 0.242 | -0.214 | 0.050 | 0.027 | 0.918 |
| **R_lingual** | -0.311 | < 0.001 | -0.297 | < 0.001 | -0.008 | 0.950 | -0.198 | 0.242 | -0.303 | 0.008 | 0.065 | 0.894 |
| **R_medialorbitofrontal** | -0.223 | < 0.001 | -0.243 | < 0.001 | 0.031 | 0.796 | -0.292 | 0.077 | -0.099 | 0.334 | -0.051 | 0.894 |
| **R_middletemporal** | -0.387 | < 0.001 | -0.305 | < 0.001 | -0.017 | 0.899 | -0.498 | 0.002 | -0.244 | 0.028 | -0.048 | 0.894 |
| **R_parahippocampal** | -0.189 | < 0.001 | -0.137 | 0.001 | -0.025 | 0.796 | 0.050 | 0.790 | 0.139 | 0.197 | -0.049 | 0.894 |

| | | | | | | | | | | | | |
|---|---|---|---|---|---|---|---|---|---|---|---|---|
| **R_paracentral** | -0.176 | < 0.001 | -0.297 | < 0.001 | 0.055 | 0.705 | -0.194 | 0.242 | -0.220 | 0.049 | 0.047 | 0.894 |
| **R_parsopercularis** | -0.485 | < 0.001 | -0.381 | < 0.001 | -0.032 | 0.796 | -0.342 | 0.036 | -0.203 | 0.060 | -0.018 | 0.918 |
| **R_parsorbitalis** | -0.342 | < 0.001 | -0.211 | < 0.001 | -0.047 | 0.705 | -0.171 | 0.301 | -0.073 | 0.458 | -0.028 | 0.918 |
| **R_parstriangularis** | -0.434 | < 0.001 | -0.380 | < 0.001 | -0.016 | 0.899 | -0.338 | 0.037 | -0.197 | 0.067 | -0.036 | 0.918 |
| **R_pericalcarine** | -0.144 | 0.003 | -0.162 | < 0.001 | 0.003 | 0.960 | 0.001 | 0.991 | -0.128 | 0.227 | 0.067 | 0.894 |
| **R_postcentral** | -0.236 | < 0.001 | -0.229 | < 0.001 | 0.006 | 0.950 | -0.153 | 0.359 | -0.118 | 0.257 | -0.009 | 0.918 |
| **R_posteriorcingulate** | -0.253 | < 0.001 | -0.318 | < 0.001 | 0.007 | 0.950 | -0.341 | 0.036 | -0.293 | 0.009 | 0.026 | 0.918 |
| **R_precentral** | -0.413 | < 0.001 | -0.296 | < 0.001 | -0.045 | 0.705 | -0.531 | 0.002 | -0.247 | 0.027 | -0.080 | 0.894 |
| **R_precuneus** | -0.289 | < 0.001 | -0.348 | < 0.001 | 0.014 | 0.907 | -0.281 | 0.091 | -0.352 | 0.003 | 0.082 | 0.894 |
| **R_rostralanteriorcingulate** | -0.221 | < 0.001 | -0.217 | < 0.001 | 0.010 | 0.925 | -0.208 | 0.229 | -0.165 | 0.123 | 0.016 | 0.918 |
| **R_rostralmiddlefrontal** | -0.378 | < 0.001 | -0.282 | < 0.001 | -0.037 | 0.781 | -0.193 | 0.242 | -0.087 | 0.393 | -0.026 | 0.918 |
| **R_superiorfrontal** | -0.508 | < 0.001 | -0.368 | < 0.001 | -0.045 | 0.705 | -0.304 | 0.065 | -0.372 | 0.003 | 0.098 | 0.894 |
| **R_superiorparietal** | -0.331 | < 0.001 | -0.245 | < 0.001 | -0.033 | 0.796 | -0.107 | 0.554 | -0.168 | 0.116 | 0.045 | 0.894 |
| **R_superiortemporal** | -0.421 | < 0.001 | -0.323 | < 0.001 | -0.025 | 0.796 | -0.667 | < 0.001 | -0.319 | 0.007 | -0.064 | 0.894 |
| **R_supramarginal** | -0.455 | < 0.001 | -0.327 | < 0.001 | -0.025 | 0.796 | -0.383 | 0.018 | -0.367 | 0.003 | 0.040 | 0.894 |
| **R_frontalpole** | -0.116 | 0.015 | -0.132 | 0.002 | 0.012 | 0.920 | 0.085 | 0.647 | -0.039 | 0.689 | 0.059 | 0.894 |
| **R_temporalpole** | -0.053 | 0.267 | -0.057 | 0.184 | 0.004 | 0.950 | -0.195 | 0.242 | -0.128 | 0.227 | 0.008 | 0.918 |
| **R_transversetemporal** | -0.267 | < 0.001 | -0.206 | < 0.001 | -0.024 | 0.796 | -0.458 | 0.004 | -0.234 | 0.035 | -0.063 | 0.894 |
| **R_insula** | -0.335 | < 0.001 | -0.327 | < 0.001 | 0.005 | 0.950 | -0.525 | 0.002 | -0.257 | 0.023 | -0.047 | 0.894 |

**Supplementary Table 18.** Effect size of age in subcortical volume in male and female patients with TLE and GGE

| | TLE | | | | | | GGE | | | | | |
|---|---|---|---|---|---|---|---|---|---|---|---|---|
| | Male | | Female | | Age-sex interaction | | Male | | Female | | Age-sex interaction | |
| **Regions** | **d** | ***p* (FDR-corrected)** | **d** | ***p* (FDR-corrected)** | **d** | ***p* (FDR-corrected)** | **d** | ***p* (FDR-corrected)** | **d** | ***p* (FDR-corrected)** | **d** | ***p* (FDR-corrected)** |
| Left accumbens | -0.425 | < 0.001 | -0.355 | < 0.001 | -0.058 | 0.665 | -0.305 | 0.032 | -0.314 | 0.001 | 0.030 | 0.843 |
| Left amygdala | -0.220 | < 0.001 | -0.149 | < 0.001 | -0.033 | 0.822 | -0.231 | 0.093 | -0.346 | < 0.001 | 0.096 | 0.843 |
| Left caudate | -0.361 | < 0.001 | -0.356 | < 0.001 | -0.016 | 0.920 | -0.469 | 0.001 | -0.554 | < 0.001 | 0.078 | 0.843 |
| Left hippocampus | -0.192 | < 0.001 | -0.088 | 0.039 | -0.052 | 0.665 | -0.345 | 0.017 | -0.355 | < 0.001 | 0.059 | 0.843 |
| Left pallidum | -0.124 | 0.009 | -0.122 | 0.005 | -0.004 | 0.920 | -0.520 | < 0.001 | -0.389 | < 0.001 | 0.021 | 0.843 |
| Left putamen | -0.339 | < 0.001 | -0.382 | < 0.001 | 0.006 | 0.920 | -0.686 | < 0.001 | -0.556 | < 0.001 | 0.022 | 0.843 |
| Left thalamus | -0.436 | < 0.001 | -0.371 | < 0.001 | -0.037 | 0.822 | -0.866 | < 0.001 | -0.609 | < 0.001 | 0.007 | 0.929 |
| Right accumbens | -0.371 | < 0.001 | -0.308 | < 0.001 | -0.041 | 0.822 | -0.506 | < 0.001 | -0.342 | < 0.001 | -0.022 | 0.843 |
| Right amygdala | -0.134 | 0.005 | -0.159 | < 0.001 | 0.009 | 0.920 | -0.262 | 0.062 | -0.342 | < 0.001 | 0.087 | 0.843 |
| Right caudate | -0.314 | < 0.001 | -0.348 | < 0.001 | 0.003 | 0.920 | -0.447 | 0.002 | -0.578 | < 0.001 | 0.099 | 0.843 |
| Right hippocampus | -0.187 | < 0.001 | -0.175 | < 0.001 | -0.008 | 0.920 | -0.204 | 0.130 | -0.228 | 0.017 | 0.054 | 0.843 |
| Right pallidum | -0.107 | 0.022 | -0.088 | 0.039 | -0.011 | 0.920 | -0.500 | < 0.001 | -0.393 | < 0.001 | 0.057 | 0.843 |
| Right putamen | -0.337 | < 0.001 | -0.379 | < 0.001 | 0.012 | 0.920 | -0.646 | < 0.001 | -0.519 | < 0.001 | 0.054 | 0.843 |
| Right thalamus | -0.383 | < 0.001 | -0.376 | < 0.001 | -0.008 | 0.920 | -0.761 | < 0.001 | -0.541 | < 0.001 | 0.033 | 0.843 |

**Supplementary Table 19.** Summary of effect size of sex interaction effects in cortical thickness in TLE in ENIGMA dataset

| | Sex-diagnosis interaction | | Sex-duration interaction | | Sex-onset interaction | |
|---|---|---|---|---|---|---|
| **Regions** | **d** | ***p* (FDR-corrected)** | **d** | ***p* (FDR-corrected)** | **d** | ***p* (FDR-corrected)** |
| **L_bankssts** | -0.003 | 0.993 | -0.029 | 0.768 | 0.002 | 0.973 |
| **L_caudalanteriorcingulate** | -0.002 | 0.993 | -0.014 | 0.945 | -0.073 | 0.417 |
| **L_caudalmiddlefrontal** | -0.006 | 0.993 | -0.043 | 0.604 | -0.005 | 0.960 |
| **L_cuneus** | -0.012 | 0.936 | 0.024 | 0.897 | -0.058 | 0.628 |
| **L_entorhinal** | 0.039 | 0.936 | -0.065 | 0.604 | 0.022 | 0.886 |
| **L_fusiform** | 0.027 | 0.936 | -0.056 | 0.604 | 0.013 | 0.942 |
| **L_inferiorparietal** | -0.008 | 0.993 | -0.006 | 0.945 | -0.035 | 0.860 |
| **L_inferiortemporal** | 0.023 | 0.936 | -0.012 | 0.945 | -0.049 | 0.831 |
| **L_isthmuscingulate** | 0.004 | 0.993 | -0.059 | 0.604 | 0.020 | 0.886 |
| **L_lateraloccipital** | 0.023 | 0.936 | 0.000 | 0.993 | -0.066 | 0.418 |
| **L_lateralorbitofrontal** | -0.003 | 0.993 | -0.030 | 0.768 | 0.030 | 0.860 |
| **L_lingual** | -0.017 | 0.936 | 0.008 | 0.945 | -0.021 | 0.886 |
| **L_medialorbitofrontal** | 0.018 | 0.936 | -0.039 | 0.604 | 0.049 | 0.831 |
| **L_middletemporal** | 0.014 | 0.936 | -0.082 | 0.349 | -0.022 | 0.886 |
| **L_parahippocampal** | -0.015 | 0.936 | 0.062 | 0.604 | -0.034 | 0.860 |
| **L_paracentral** | -0.002 | 0.993 | -0.004 | 0.965 | -0.070 | 0.417 |
| **L_parsopercularis** | -0.011 | 0.936 | -0.050 | 0.604 | 0.005 | 0.960 |
| **L_parsorbitalis** | -0.027 | 0.936 | -0.050 | 0.604 | 0.010 | 0.960 |
| **L_parstriangularis** | 0.005 | 0.993 | -0.093 | 0.349 | 0.029 | 0.860 |
| **L_pericalcarine** | 0.011 | 0.936 | 0.039 | 0.604 | -0.047 | 0.848 |
| **L_postcentral** | -0.033 | 0.936 | 0.017 | 0.945 | -0.039 | 0.860 |
| **L_posteriorcingulate** | 0.019 | 0.936 | -0.035 | 0.664 | -0.032 | 0.860 |
| **L_precentral** | -0.012 | 0.936 | -0.012 | 0.945 | -0.045 | 0.848 |

| | | | | | | |
|---|---|---|---|---|---|---|
| **L_precuneus** | -0.012 | 0.936 | -0.027 | 0.836 | -0.035 | 0.860 |
| **L_rostralanteriorcingulate** | -0.038 | 0.936 | -0.007 | 0.945 | 0.003 | 0.960 |
| **L_rostralmiddlefrontal** | 0.034 | 0.936 | -0.049 | 0.604 | -0.019 | 0.887 |
| **L_superiorfrontal** | -0.016 | 0.936 | -0.047 | 0.604 | 0.001 | 0.979 |
| **L_superiorparietal** | -0.038 | 0.936 | 0.001 | 0.993 | -0.085 | 0.417 |
| **L_superiortemporal** | -0.012 | 0.936 | -0.033 | 0.722 | -0.025 | 0.886 |
| **L_supramarginal** | -0.015 | 0.936 | -0.055 | 0.604 | -0.015 | 0.915 |
| **L_frontalpole** | 0.019 | 0.936 | -0.007 | 0.945 | 0.034 | 0.860 |
| **L_temporalpole** | -0.025 | 0.936 | -0.011 | 0.945 | -0.025 | 0.886 |
| **L_transversetemporal** | -0.019 | 0.936 | -0.014 | 0.945 | -0.005 | 0.960 |
| **L_insula** | 0.006 | 0.993 | -0.038 | 0.623 | 0.027 | 0.860 |
| **R_bankssts** | 0.026 | 0.936 | -0.040 | 0.604 | 0.006 | 0.960 |
| **R_caudalanteriorcingulate** | -0.004 | 0.993 | -0.010 | 0.945 | -0.041 | 0.860 |
| **R_caudalmiddlefrontal** | -0.021 | 0.936 | -0.048 | 0.604 | -0.003 | 0.960 |
| **R_cuneus** | -0.018 | 0.936 | 0.011 | 0.945 | -0.023 | 0.886 |
| **R_entorhinal** | 0.001 | 0.997 | -0.083 | 0.349 | 0.022 | 0.886 |
| **R_fusiform** | -0.014 | 0.936 | -0.011 | 0.945 | -0.043 | 0.860 |
| **R_inferiorparietal** | 0.006 | 0.993 | 0.006 | 0.945 | -0.040 | 0.860 |
| **R_inferiortemporal** | -0.012 | 0.936 | 0.015 | 0.945 | -0.075 | 0.417 |
| **R_isthmuscingulate** | -0.020 | 0.936 | -0.045 | 0.604 | 0.030 | 0.860 |
| **R_lateraloccipital** | 0.025 | 0.936 | 0.055 | 0.604 | -0.077 | 0.417 |
| **R_lateralorbitofrontal** | -0.029 | 0.936 | 0.009 | 0.945 | -0.030 | 0.860 |
| **R_lingual** | -0.026 | 0.936 | 0.016 | 0.945 | -0.045 | 0.848 |
| **R_medialorbitofrontal** | 0.030 | 0.936 | 0.018 | 0.945 | -0.006 | 0.960 |
| **R_middletemporal** | 0.007 | 0.993 | -0.045 | 0.604 | -0.017 | 0.915 |
| **R_parahippocampal** | 0.036 | 0.936 | 0.017 | 0.945 | -0.067 | 0.418 |
| **R_paracentral** | -0.012 | 0.936 | 0.050 | 0.604 | -0.028 | 0.860 |
| **R_parsopercularis** | -0.015 | 0.936 | -0.054 | 0.604 | -0.011 | 0.960 |

| | | | | | | |
|---|---|---|---|---|---|---|
| **R_parsorbitalis** | -0.048 | 0.936 | -0.065 | 0.604 | 0.010 | 0.960 |
| **R_parstriangularis** | 0.000 | 0.997 | -0.010 | 0.945 | -0.021 | 0.886 |
| **R_pericalcarine** | -0.018 | 0.936 | -0.009 | 0.945 | -0.005 | 0.960 |
| **R_postcentral** | -0.057 | 0.936 | 0.015 | 0.945 | -0.034 | 0.860 |
| **R_posteriorcingulate** | 0.005 | 0.993 | -0.003 | 0.987 | -0.005 | 0.960 |
| **R_precentral** | -0.026 | 0.936 | -0.018 | 0.945 | -0.029 | 0.860 |
| **R_precuneus** | -0.035 | 0.936 | 0.010 | 0.945 | -0.029 | 0.860 |
| **R_rostralanteriorcingulate** | -0.011 | 0.936 | 0.001 | 0.993 | -0.003 | 0.960 |
| **R_rostralmiddlefrontal** | -0.016 | 0.936 | -0.040 | 0.604 | -0.020 | 0.886 |
| **R_superiorfrontal** | 0.003 | 0.993 | -0.051 | 0.604 | -0.015 | 0.915 |
| **R_superiorparietal** | -0.024 | 0.936 | 0.029 | 0.768 | -0.082 | 0.417 |
| **R_superiortemporal** | 0.003 | 0.993 | -0.040 | 0.604 | -0.016 | 0.915 |
| **R_supramarginal** | -0.042 | 0.936 | -0.045 | 0.604 | -0.016 | 0.915 |
| **R_frontalpole** | -0.032 | 0.936 | 0.000 | 0.993 | 0.005 | 0.960 |
| **R_temporalpole** | 0.000 | 0.997 | 0.021 | 0.945 | -0.053 | 0.810 |
| **R_transversetemporal** | -0.031 | 0.936 | -0.041 | 0.604 | 0.003 | 0.960 |
| **R_insula** | -0.055 | 0.936 | -0.020 | 0.945 | -0.006 | 0.960 |

**Supplementary Table 20.** Summary of effect size of sex interaction effects in subcortical volume in TLE in ENIGMA dataset

| | Sex-diagnosis interaction | | Sex-duration interaction | | Sex-onset interaction | |
|---|---|---|---|---|---|---|
| **Regions** | **d** | ***p* (FDR-corrected)** | **d** | ***p* (FDR-corrected)** | **d** | ***p* (FDR-corrected)** |
| **Left accumbens** | -0.038 | 0.624 | -0.071 | 0.401 | -0.003 | 0.962 |
| **Left amygdala** | 0.046 | 0.624 | -0.041 | 0.401 | 0.012 | 0.923 |
| **Left caudate** | -0.016 | 0.624 | -0.045 | 0.401 | 0.038 | 0.823 |
| **Left hippocampus** | 0.018 | 0.624 | -0.037 | 0.401 | -0.002 | 0.962 |
| **Left pallidum** | -0.028 | 0.624 | -0.033 | 0.411 | 0.028 | 0.823 |
| **Left putamen** | 0.022 | 0.624 | -0.038 | 0.401 | 0.020 | 0.923 |
| **Left thalamus** | -0.023 | 0.624 | -0.038 | 0.401 | -0.029 | 0.823 |
| **Right accumbens** | -0.031 | 0.624 | -0.052 | 0.401 | -0.016 | 0.923 |
| **Right amygdala** | 0.035 | 0.624 | -0.010 | 0.768 | -0.007 | 0.962 |
| **Right caudate** | -0.008 | 0.812 | -0.037 | 0.401 | 0.043 | 0.823 |
| **Right hippocampus** | 0.029 | 0.624 | 0.023 | 0.525 | -0.037 | 0.823 |
| **Right pallidum** | 0.004 | 0.886 | -0.036 | 0.401 | 0.033 | 0.823 |
| **Right putamen** | 0.022 | 0.624 | -0.036 | 0.401 | 0.028 | 0.823 |
| **Right thalamus** | -0.019 | 0.624 | -0.029 | 0.459 | -0.012 | 0.923 |

**Supplementary Table 21.** Summary of effect size of sex interaction effects in cortical thickness in TLE in ECP and MNI datasets

| | ECP | | | | | | MNI | | | | | |
|---|---|---|---|---|---|---|---|---|---|---|---|---|
| | Sex-diagnosis interaction | | Sex-duration interaction | | Sex-onset interaction | | Sex-diagnosis interaction | | Sex-duration interaction | | Sex-onset interaction | |
| **Regions** | **d** | ***p* (FDR-corrected)** | **d** | ***p* (FDR-corrected)** | **d** | ***p* (FDR-corrected)** | **d** | ***p* (FDR-corrected)** | **d** | ***p* (FDR-corrected)** | **d** | ***p* (FDR-corrected)** |
| **L_bankssts** | 0.091 | 0.821 | -0.111 | 0.808 | 0.232 | 0.945 | -0.115 | 0.910 | 0.158 | 0.970 | -0.014 | 0.975 |
| **L_caudalanteriorcingulate** | 0.064 | 0.821 | -0.176 | 0.808 | 0.163 | 0.945 | -0.006 | 0.987 | -0.096 | 0.970 | 0.132 | 0.805 |
| **L_caudalmiddlefrontal** | 0.022 | 0.909 | -0.038 | 0.929 | 0.162 | 0.945 | -0.092 | 0.910 | 0.029 | 0.970 | -0.038 | 0.963 |
| **L_cuneus** | -0.162 | 0.662 | 0.041 | 0.924 | -0.015 | 0.974 | 0.040 | 0.987 | 0.219 | 0.970 | -0.038 | 0.963 |
| **L_entorhinal** | 0.069 | 0.821 | 0.120 | 0.808 | 0.058 | 0.945 | 0.049 | 0.987 | -0.206 | 0.970 | 0.175 | 0.780 |
| **L_fusiform** | 0.012 | 0.989 | 0.101 | 0.808 | -0.061 | 0.945 | -0.054 | 0.987 | 0.047 | 0.970 | 0.037 | 0.963 |
| **L_inferiorparietal** | -0.079 | 0.821 | -0.013 | 0.980 | 0.130 | 0.945 | 0.038 | 0.987 | -0.006 | 0.970 | -0.108 | 0.881 |
| **L_inferiortemporal** | 0.039 | 0.846 | 0.231 | 0.807 | -0.087 | 0.945 | 0.009 | 0.987 | -0.094 | 0.970 | 0.244 | 0.568 |
| **L_isthmuscingulate** | -0.006 | 0.989 | 0.101 | 0.808 | 0.005 | 0.974 | 0.157 | 0.910 | -0.134 | 0.970 | 0.138 | 0.805 |
| **L_lateraloccipital** | -0.064 | 0.821 | 0.067 | 0.838 | 0.054 | 0.947 | -0.047 | 0.987 | 0.024 | 0.970 | 0.045 | 0.963 |
| **L_lateralorbitofrontal** | 0.068 | 0.821 | -0.065 | 0.838 | 0.005 | 0.974 | 0.002 | 0.987 | -0.088 | 0.970 | 0.337 | 0.416 |
| **L_lingual** | -0.116 | 0.751 | -0.009 | 0.994 | -0.030 | 0.974 | -0.108 | 0.910 | 0.092 | 0.970 | 0.143 | 0.805 |
| **L_medialorbitofrontal** | 0.044 | 0.838 | 0.144 | 0.808 | -0.049 | 0.952 | -0.047 | 0.987 | 0.196 | 0.970 | 0.128 | 0.805 |
| **L_middletemporal** | 0.051 | 0.821 | 0.089 | 0.808 | 0.075 | 0.945 | 0.023 | 0.987 | -0.061 | 0.970 | 0.113 | 0.881 |
| **L_parahippocampal** | 0.000 | 0.996 | -0.063 | 0.838 | 0.064 | 0.945 | 0.107 | 0.910 | 0.114 | 0.970 | -0.108 | 0.881 |
| **L_paracentral** | 0.117 | 0.751 | 0.094 | 0.808 | -0.039 | 0.974 | -0.070 | 0.987 | -0.048 | 0.970 | -0.006 | 0.976 |
| **L_parsopercularis** | 0.064 | 0.821 | 0.035 | 0.929 | 0.092 | 0.945 | -0.113 | 0.910 | 0.019 | 0.970 | 0.093 | 0.881 |
| **L_parsorbitalis** | 0.205 | 0.662 | 0.085 | 0.808 | -0.022 | 0.974 | -0.098 | 0.910 | -0.061 | 0.970 | 0.263 | 0.557 |
| **L_parstriangularis** | 0.115 | 0.751 | -0.071 | 0.838 | 0.089 | 0.945 | -0.049 | 0.987 | 0.040 | 0.970 | 0.088 | 0.881 |
| **L_pericalcarine** | 0.061 | 0.821 | -0.029 | 0.929 | -0.029 | 0.974 | -0.161 | 0.910 | -0.103 | 0.970 | 0.168 | 0.805 |

| | | | | | | | | | | | | |
|---|---|---|---|---|---|---|---|---|---|---|---|---|
| **L_postcentral** | 0.062 | 0.821 | 0.051 | 0.881 | 0.096 | 0.945 | -0.004 | 0.987 | 0.020 | 0.970 | 0.146 | 0.805 |
| **L_posteriorcingulate** | 0.060 | 0.821 | -0.050 | 0.881 | 0.018 | 0.974 | -0.018 | 0.987 | 0.204 | 0.970 | -0.011 | 0.975 |
| **L_precentral** | 0.112 | 0.755 | 0.062 | 0.838 | -0.012 | 0.974 | -0.047 | 0.987 | 0.084 | 0.970 | 0.062 | 0.963 |
| **L_precuneus** | 0.089 | 0.821 | -0.030 | 0.929 | 0.159 | 0.945 | 0.003 | 0.987 | 0.085 | 0.970 | -0.089 | 0.881 |
| **L_rostralanteriorcingulate** | 0.036 | 0.847 | -0.180 | 0.808 | 0.147 | 0.945 | -0.018 | 0.987 | 0.167 | 0.970 | 0.017 | 0.975 |
| **L_rostralmiddlefrontal** | 0.085 | 0.821 | -0.093 | 0.808 | 0.121 | 0.945 | -0.131 | 0.910 | -0.046 | 0.970 | 0.130 | 0.805 |
| **L_superiorfrontal** | 0.169 | 0.662 | -0.015 | 0.980 | 0.169 | 0.945 | -0.002 | 0.987 | 0.063 | 0.970 | -0.044 | 0.963 |
| **L_superiorparietal** | -0.008 | 0.989 | 0.068 | 0.838 | 0.071 | 0.945 | 0.057 | 0.987 | -0.107 | 0.970 | -0.069 | 0.963 |
| **L_superiortemporal** | 0.034 | 0.849 | 0.204 | 0.808 | -0.016 | 0.974 | -0.017 | 0.987 | 0.046 | 0.970 | 0.097 | 0.881 |
| **L_supramarginal** | 0.143 | 0.662 | 0.115 | 0.808 | -0.021 | 0.974 | 0.048 | 0.987 | 0.009 | 0.970 | -0.153 | 0.805 |
| **L_frontalpole** | 0.116 | 0.751 | 0.181 | 0.808 | -0.204 | 0.945 | -0.032 | 0.987 | -0.198 | 0.970 | 0.352 | 0.416 |
| **L_temporalpole** | 0.145 | 0.662 | -0.030 | 0.929 | 0.012 | 0.974 | 0.138 | 0.910 | -0.103 | 0.970 | 0.219 | 0.576 |
| **L_transversetemporal** | 0.038 | 0.846 | 0.074 | 0.838 | -0.035 | 0.974 | -0.046 | 0.987 | -0.005 | 0.970 | 0.087 | 0.881 |
| **L_insula** | -0.125 | 0.751 | 0.099 | 0.808 | 0.081 | 0.945 | 0.171 | 0.910 | 0.063 | 0.970 | 0.262 | 0.557 |
| **R_bankssts** | 0.170 | 0.662 | 0.143 | 0.808 | -0.061 | 0.945 | -0.015 | 0.987 | -0.016 | 0.970 | 0.103 | 0.881 |
| **R_caudalanteriorcingulate** | 0.052 | 0.821 | -0.144 | 0.808 | 0.134 | 0.945 | 0.064 | 0.987 | 0.114 | 0.970 | 0.057 | 0.963 |
| **R_caudalmiddlefrontal** | 0.068 | 0.821 | 0.001 | 0.994 | 0.121 | 0.945 | 0.059 | 0.987 | 0.008 | 0.970 | -0.023 | 0.975 |
| **R_cuneus** | -0.145 | 0.662 | -0.007 | 0.994 | -0.083 | 0.945 | 0.044 | 0.987 | 0.063 | 0.970 | -0.009 | 0.976 |
| **R_entorhinal** | 0.042 | 0.846 | 0.104 | 0.808 | -0.004 | 0.974 | 0.101 | 0.910 | -0.137 | 0.970 | 0.024 | 0.975 |
| **R_fusiform** | -0.051 | 0.821 | 0.151 | 0.808 | -0.073 | 0.945 | 0.019 | 0.987 | 0.095 | 0.970 | 0.043 | 0.963 |
| **R_inferiorparietal** | 0.031 | 0.849 | 0.158 | 0.808 | 0.032 | 0.974 | -0.073 | 0.987 | -0.014 | 0.970 | -0.002 | 0.991 |
| **R_inferiortemporal** | -0.032 | 0.849 | 0.249 | 0.751 | -0.095 | 0.945 | 0.035 | 0.987 | 0.210 | 0.970 | -0.040 | 0.963 |
| **R_isthmuscingulate** | -0.032 | 0.849 | -0.067 | 0.838 | 0.140 | 0.945 | 0.139 | 0.910 | -0.132 | 0.970 | 0.037 | 0.963 |
| **R_lateraloccipital** | -0.006 | 0.989 | 0.132 | 0.808 | -0.110 | 0.945 | -0.112 | 0.910 | 0.118 | 0.970 | -0.144 | 0.805 |
| **R_lateralorbitofrontal** | -0.011 | 0.989 | -0.048 | 0.881 | 0.060 | 0.945 | -0.090 | 0.910 | -0.100 | 0.970 | 0.305 | 0.488 |
| **R_lingual** | -0.049 | 0.821 | 0.003 | 0.994 | -0.066 | 0.945 | -0.001 | 0.987 | 0.182 | 0.970 | 0.055 | 0.963 |
| **R_medialorbitofrontal** | 0.039 | 0.846 | -0.119 | 0.808 | 0.053 | 0.947 | -0.037 | 0.987 | -0.016 | 0.970 | 0.292 | 0.488 |
| **R_middletemporal** | 0.109 | 0.755 | 0.112 | 0.808 | 0.074 | 0.945 | -0.041 | 0.987 | 0.011 | 0.970 | 0.217 | 0.576 |

| | | | | | | | | | | | | |
|---|---|---|---|---|---|---|---|---|---|---|---|---|
| **R_parahippocampal** | -0.050 | 0.821 | -0.059 | 0.846 | 0.125 | 0.945 | 0.100 | 0.910 | 0.040 | 0.970 | -0.060 | 0.963 |
| **R_paracentral** | 0.008 | 0.989 | 0.168 | 0.808 | -0.132 | 0.945 | -0.008 | 0.987 | 0.111 | 0.970 | -0.030 | 0.975 |
| **R_parsopercularis** | 0.166 | 0.662 | 0.015 | 0.980 | 0.210 | 0.945 | -0.046 | 0.987 | 0.066 | 0.970 | 0.130 | 0.805 |
| **R_parsorbitalis** | 0.074 | 0.821 | -0.019 | 0.980 | -0.040 | 0.974 | 0.045 | 0.987 | -0.042 | 0.970 | 0.132 | 0.805 |
| **R_parstriangularis** | 0.098 | 0.821 | 0.254 | 0.751 | -0.068 | 0.945 | 0.095 | 0.910 | 0.027 | 0.970 | 0.131 | 0.805 |
| **R_pericalcarine** | 0.002 | 0.995 | -0.096 | 0.808 | -0.090 | 0.945 | -0.115 | 0.910 | 0.134 | 0.970 | 0.029 | 0.975 |
| **R_postcentral** | -0.055 | 0.821 | 0.204 | 0.808 | 0.006 | 0.974 | 0.024 | 0.987 | 0.027 | 0.970 | 0.019 | 0.975 |
| **R_posteriorcingulate** | 0.065 | 0.821 | -0.090 | 0.808 | 0.172 | 0.945 | 0.109 | 0.910 | 0.033 | 0.970 | 0.101 | 0.881 |
| **R_precentral** | 0.045 | 0.838 | 0.109 | 0.808 | -0.052 | 0.947 | -0.032 | 0.987 | -0.016 | 0.970 | 0.051 | 0.963 |
| **R_precuneus** | -0.057 | 0.821 | -0.003 | 0.994 | 0.109 | 0.945 | -0.031 | 0.987 | 0.163 | 0.970 | -0.072 | 0.963 |
| **R_rostralanteriorcingulate** | 0.029 | 0.856 | -0.092 | 0.808 | 0.062 | 0.945 | 0.044 | 0.987 | 0.148 | 0.970 | -0.023 | 0.975 |
| **R_rostralmiddlefrontal** | 0.071 | 0.821 | 0.028 | 0.929 | 0.044 | 0.974 | 0.053 | 0.987 | 0.044 | 0.970 | 0.089 | 0.881 |
| **R_superiorfrontal** | 0.106 | 0.759 | -0.079 | 0.838 | 0.151 | 0.945 | 0.011 | 0.987 | 0.020 | 0.970 | -0.012 | 0.975 |
| **R_superiorparietal** | -0.054 | 0.821 | 0.112 | 0.808 | 0.034 | 0.974 | 0.023 | 0.987 | -0.015 | 0.970 | -0.055 | 0.963 |
| **R_superiortemporal** | 0.069 | 0.821 | 0.171 | 0.808 | 0.009 | 0.974 | -0.080 | 0.987 | 0.039 | 0.970 | 0.188 | 0.702 |
| **R_supramarginal** | 0.139 | 0.662 | 0.159 | 0.808 | 0.129 | 0.945 | -0.028 | 0.987 | 0.170 | 0.970 | -0.210 | 0.585 |
| **R_frontalpole** | 0.086 | 0.821 | 0.086 | 0.808 | -0.082 | 0.945 | -0.035 | 0.987 | -0.035 | 0.970 | 0.199 | 0.645 |
| **R_temporalpole** | 0.087 | 0.821 | 0.315 | 0.516 | -0.198 | 0.945 | 0.088 | 0.910 | 0.081 | 0.970 | 0.234 | 0.568 |
| **R_transversetemporal** | 0.138 | 0.662 | 0.156 | 0.808 | -0.084 | 0.945 | 0.007 | 0.987 | -0.178 | 0.970 | 0.239 | 0.568 |
| **R_insula** | -0.004 | 0.995 | 0.090 | 0.808 | 0.105 | 0.945 | 0.014 | 0.987 | -0.048 | 0.970 | 0.229 | 0.568 |

**Supplementary Table 22.** Summary of effect size of sex interaction effects in subcortical volume in TLE in ECP and MNI datasets

| | ECP | | | | | | MNI | | | | | |
|---|---|---|---|---|---|---|---|---|---|---|---|---|
| | Sex-diagnosis interaction | | Sex-duration interaction | | Sex-onset interaction | | Sex-diagnosis interaction | | Sex-duration interaction | | Sex-onset interaction | |
| **Regions** | **d** | ***p* (FDR-corrected)** | **d** | ***p* (FDR-corrected)** | **d** | ***p* (FDR-corrected)** | **d** | ***p* (FDR-corrected)** | **d** | ***p* (FDR-corrected)** | **d** | ***p* (FDR-corrected)** |
| **Left accumbens** | -0.032 | 0.924 | 0.081 | 0.614 | -0.107 | 0.554 | -0.123 | 0.549 | -0.036 | 0.994 | 0.296 | 0.270 |
| **Left amygdala** | -0.108 | 0.906 | 0.077 | 0.614 | -0.132 | 0.443 | 0.113 | 0.549 | 0.137 | 0.994 | -0.012 | 0.929 |
| **Left caudate** | -0.085 | 0.924 | 0.073 | 0.614 | -0.240 | 0.357 | -0.027 | 0.903 | -0.053 | 0.994 | 0.129 | 0.394 |
| **Left hippocampus** | -0.063 | 0.924 | -0.116 | 0.614 | -0.055 | 0.741 | 0.002 | 0.980 | 0.210 | 0.790 | -0.235 | 0.270 |
| **Left pallidum** | -0.037 | 0.924 | 0.146 | 0.614 | -0.150 | 0.392 | -0.130 | 0.549 | 0.008 | 0.994 | -0.156 | 0.331 |
| **Left putamen** | -0.131 | 0.828 | 0.094 | 0.614 | -0.072 | 0.741 | -0.083 | 0.743 | -0.054 | 0.994 | 0.212 | 0.270 |
| **Left thalamus** | 0.022 | 0.924 | 0.055 | 0.682 | 0.019 | 0.925 | 0.023 | 0.903 | 0.098 | 0.994 | -0.126 | 0.394 |
| **Right accumbens** | -0.012 | 0.926 | 0.097 | 0.614 | -0.059 | 0.741 | -0.076 | 0.743 | 0.013 | 0.994 | 0.225 | 0.270 |
| **Right amygdala** | -0.041 | 0.924 | 0.084 | 0.614 | -0.208 | 0.357 | 0.003 | 0.980 | 0.084 | 0.994 | 0.188 | 0.270 |
| **Right caudate** | -0.044 | 0.924 | 0.106 | 0.614 | -0.158 | 0.392 | -0.028 | 0.903 | -0.065 | 0.994 | 0.199 | 0.270 |
| **Right hippocampus** | -0.028 | 0.924 | 0.037 | 0.749 | -0.182 | 0.392 | 0.071 | 0.743 | 0.251 | 0.790 | -0.165 | 0.326 |
| **Right pallidum** | -0.008 | 0.926 | 0.093 | 0.614 | -0.206 | 0.357 | -0.126 | 0.549 | -0.001 | 0.994 | -0.208 | 0.270 |
| **Right putamen** | -0.130 | 0.828 | 0.240 | 0.557 | -0.155 | 0.392 | -0.056 | 0.839 | -0.045 | 0.994 | 0.196 | 0.270 |
| **Right thalamus** | 0.033 | 0.924 | 0.087 | 0.614 | 0.011 | 0.925 | -0.024 | 0.903 | 0.017 | 0.994 | 0.032 | 0.870 |

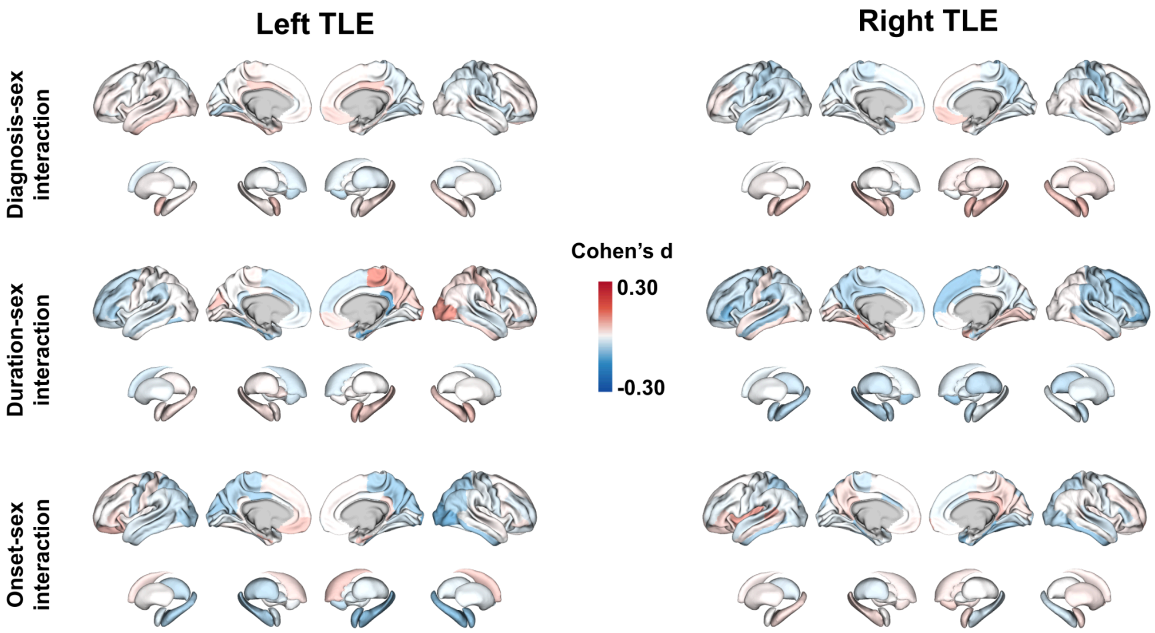


**Supplementary Figure 1. Subcohort analyses of sex differences across left and right TLE subgroups.** Effect sizes in the merged dataset are shown for the diagnosis-by-sex, disease-duration-by-sex, and age-of-onset-by-sex interactions in left and right TLE patients. Significant regions were outlined by dark lines with a threshold of $P_{\mathrm{FDR}} < 0.05$. For the upper panel, positive values (red) indicate greater cortical thickness or subcortical volume in male than female patients relative to their corresponding healthy controls, shown separately for left and right TLE. For middle and lower panels, positive values (red) indicate regions where male patients show stronger associations with disease duration or age of onset, shown separately for left and right TLE. TLE: temporal lobe epilepsy; Duration: disease duration; Onset: age of onset.

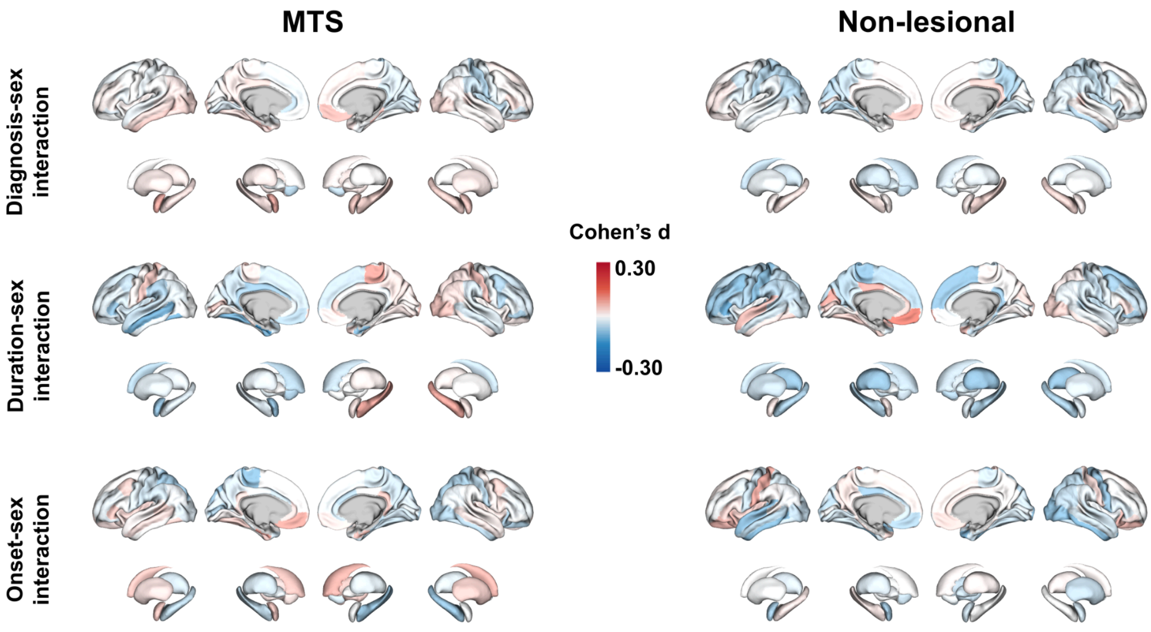


**Supplementary Figure 2. Sex differences in mesial temporal sclerosis (MTS) and non-lesional TLE patients.** Effect sizes in the merged dataset are shown for the diagnosis-by-sex, disease-duration-by-sex, and age-of-onset-by-sex interactions in MTS and non-lesional TLE patients. Significant regions were outlined by dark lines with a threshold of $P_{FDR} < 0.05$. For the upper panel, positive values (red) indicate greater cortical thickness or subcortical volume in male than female patients relative to their corresponding healthy controls, shown separately for MTS and Non-lesional subgroups. For middle and lower panels, positive values (red) indicate regions where male patients show stronger associations with disease duration or age of onset, shown separately for MTS and Non-lesional subgroups. TLE: temporal lobe epilepsy; Duration: disease duration; Onset: age of onset.

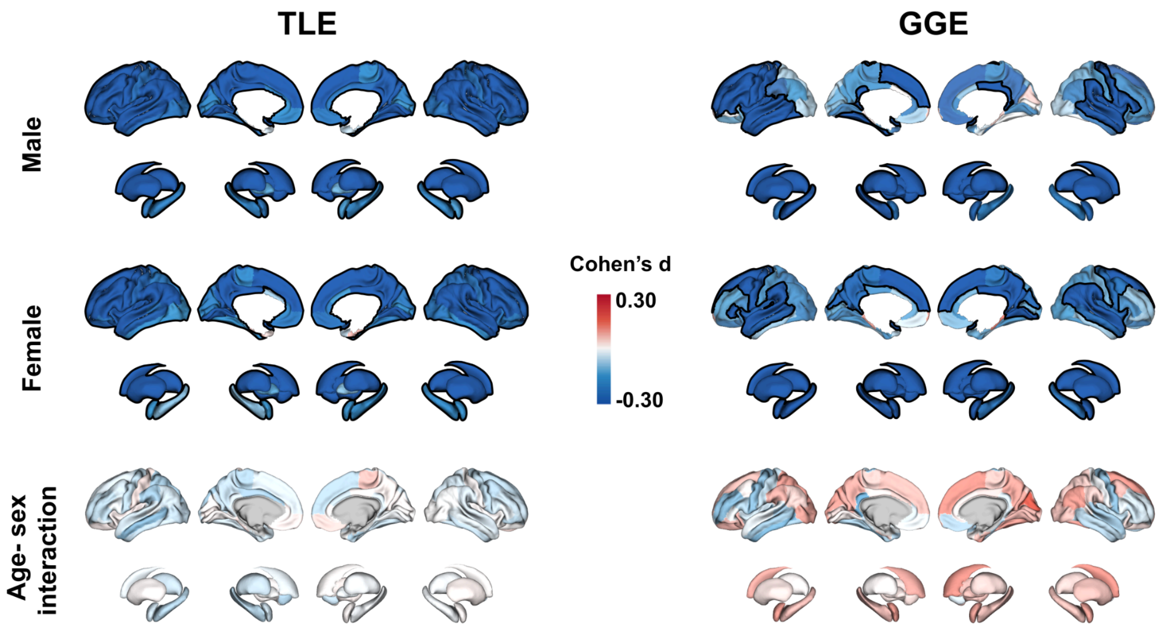


**Supplementary Figure 3. Sex differences in the effects of age in TLE and GGE.** Effect sizes in the merged dataset are shown for the main effect of age (upper and middle panels) and for the disease duration-by-sex interaction (lower panel). Significant regions were outlined by dark lines with a threshold of $P_{FDR} < 0.05$. Positive values (red) indicate regions where greater age corresponds to greater cortical thickness and subcortical volume, or where males exhibit stronger age effects. TLE: temporal lobe epilepsy; GGE: genetic generalized epilepsy; Duration: disease duration; Onset: age of onset.

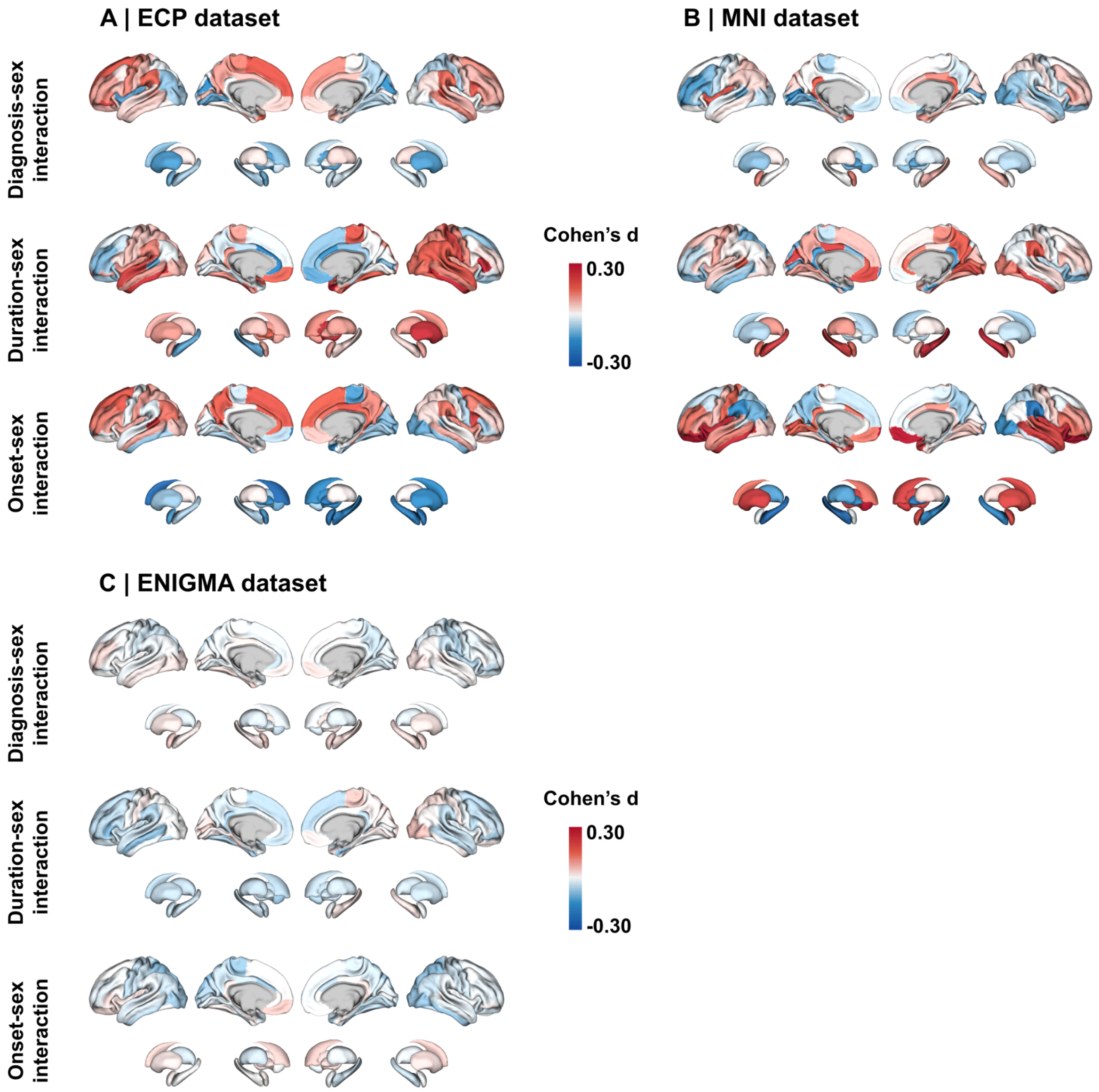


**Supplementary Figure 4. Consistency of sex interaction effects analyses in TLE samples across datasets.** **A**. Effect sizes in the ECP dataset are shown for the diagnosis-by-sex, disease-duration-by-sex, and age-of-onset-by-sex interactions. **B**. Effect sizes in the MNI dataset are shown for the diagnosis-by-sex, disease-duration-by-sex, and age-of-onset-by-sex interactions. **C**. Effect sizes in the ENIGMA dataset are shown for the diagnosis-by-sex, disease-duration-by-sex, and age-of-onset-by-sex interactions. Statistically significant regions are outlined by dark borders ($P_{FDR} < 0.05$). TLE: temporal lobe epilepsy; Duration: disease duration; Onset: age of onset.
